# Whole-cortex in situ sequencing reveals peripheral input-dependent cell type-defined area identity

**DOI:** 10.1101/2022.11.06.515380

**Authors:** Xiaoyin Chen, Stephan Fischer, Mara CP Rue, Aixin Zhang, Didhiti Mukherjee, Patrick O Kanold, Jesse Gillis, Anthony M Zador

## Abstract

The cortex is composed of neuronal types with diverse gene expression that are organized into specialized cortical areas. These areas, each with characteristic cytoarchitecture (Brodmann 1909; Vogt and Vogt 1919; Von Bonin 1947), connectivity (Zingg et al. 2014; Harris et al. 2019), and neuronal activity (Schwarz et al. 2008; Ferrarini et al. 2009; He et al. 2009; Meunier et al. 2010; Bertolero et al. 2015), are wired into modular networks (Zingg et al. 2014; Harris et al. 2019; Huang et al. 2020). However, it remains unclear whether cortical areas and their modular organization can be similarly defined by their transcriptomic signatures and how such signatures are established in development. Here we used BARseq, a high-throughput *in situ* sequencing technique, to interrogate the expression of 104 cell type marker genes in 10.3 million cells, including 4,194,658 cortical neurons over nine mouse forebrain hemispheres at cellular resolution. *De novo* clustering of gene expression in single neurons revealed transcriptomic types that were consistent with previous single-cell RNAseq studies(Yao et al. 2021a; Yao et al. 2021b). Gene expression and the distribution of fine-grained cell types vary along the contours of cortical areas, and the composition of transcriptomic types are highly predictive of cortical area identity. Moreover, areas with similar compositions of transcriptomic types, which we defined as cortical modules, overlap with areas that are highly connected, suggesting that the same modular organization is reflected in both transcriptomic signatures and connectivity. To explore how the transcriptomic profiles of cortical neurons depend on development, we compared the cell type distributions after neonatal binocular enucleation. Strikingly, binocular enucleation caused the cell type compositional profiles of visual areas to shift towards neighboring areas within the same cortical module, suggesting that peripheral inputs sharpen the distinct transcriptomic identities of areas within cortical modules. Enabled by the high-throughput, low-cost, and reproducibility of BARseq, our study provides a proof-of-principle for using large-scale *in situ* sequencing to reveal brain-wide molecular architecture and to understand its development.

## Main text

The vertebrate brain is organized into subregions that are specialized in function and distinct in cytoarchitecture and connectivity. This spatial specialization of function and structure is established by developmental processes involving intrinsic genetic programs and/or external signaling (Cadwell et al. 2019). Both intrinsic and extrinsic developmental processes drive the expression of specific sets of genes, which together specify the cell fates and establish specialized cellular properties (Hobert 2008). Although gene expression can change during cell maturation and remains dynamic in response to internal cellular conditions and external stimuli, a core transcriptional program that maintains the cellular identity usually remains steady in mature neurons (Zeng and Sanes 2017). Thus, resolving the expression of core sets of genes that distinguish different types of neurons provides insight into the functional and structural specialization of neurons.

Many large brain structures are spatially organized into divisions, or modules, within which neurons are more similar in morphology, connectivity, and activity. In the cortex, these modules usually involve a set of adjacent cortical areas that are highly interconnected (Zingg et al. 2014; Harris et al. 2019; Huang et al. 2020) and correlated in neuronal activity (Schwarz et al. 2008; Ferrarini et al. 2009; He et al. 2009; Meunier et al. 2010; Bertolero et al. 2015). This modular organization is perturbed in Alzheimer’s disease (Stam et al. 2007), mild cognitive impairment (Yao et al. 2010), schizophrenia (Lynall et al. 2010), and depression (Fingelkurts et al. 2007), suggesting that cortical modules are critical to normal functions. In contrast to this modular organization in activity and connectivity, many cortical areas share the same medium-grained and fine-grained transcriptomically defined neuronal types (Tasic et al. 2018; Yao et al. 2021b). Whether and how the areal and modular organization of cortical connectivity and activity is reflected in the transcriptomic signatures of areas is unknown.

To address this question, here we apply BARseq (Chen et al. 2021b; Sun et al. 2021) to interrogate gene expression and the distribution of excitatory neuron types across nine mouse forebrain hemispheres at high spatial resolution. BARseq is a form of *in situ* sequencing (Ke et al. 2013; Qian et al. 2020; Bugeon et al. 2022), in which Illumina sequencing-by-synthesis chemistry is used to achieve a robust readout of both endogenous mRNAs and synthetic RNA barcodes. These RNA barcodes are used to infer long-range projections of neurons. We have previously used BARseq to identify the projection patterns of different neuronal types defined either by gene expression (Chen et al. 2019; Sun et al. 2021) and/or their locations (Chen et al. 2021b; Munoz-Castaneda et al. 2021), and to identify genes that are associated with differences in projections within neuronal populations (Sun et al. 2021). Importantly, we showed that BARseq can resolve transcriptomically defined cell types of cortical neurons at cellular resolution by sequencing dozens of cell type markers (Sun et al. 2021). Because BARseq has high throughput and low cost compared to many other spatial techniques (Chen et al. 2015; Shah et al. 2016; Stahl et al. 2016; Codeluppi et al. 2018; Qian et al. 2020; Stickels et al. 2021; Chen et al. 2022), it is ideally suited for studying the spatial organization of gene expression at cellular resolution over whole brain structures, such as the cortex.

Here we use BARseq as a standalone technique for sequencing gene expression *in situ*, and scale it up to a brain-wide scale in nine animals, with or without binocular enucleation, to resolve the distribution of neuronal populations and gene expression across the cortex. We generate high-resolution maps of 10.3 million cells with detailed gene expression, including 4,194,658 cortical cells. We find that although most neuronal populations are found in multiple cortical areas, the composition of neuronal populations is distinct across areas. The neuronal compositions of highly connected areas are more similar, suggesting a modular organization of the cortex that matches cortical hierarchy and modules defined by connectivity in previous studies (Zingg et al. 2014; Harris et al. 2019; Huang et al. 2020). By comparing littermates with and without binocular enucleation, we then show that peripheral inputs play a critical role in shaping cortical gene expression and area-specific cell type compositional profiles.

### Cellular resolution BARseq captures brain-wide gene expression in situ

A number of recent single-cell transcriptomic studies (Zeisel et al. 2015; Paul et al. 2017; Tasic et al. 2018; Zeisel et al. 2018; Yao et al. 2021a) have classified neuronal types using hierarchical analyses, but these studies used different nomenclatures to refer to cell types at different levels of the hierarchy. To avoid confusion, we first define the cell type nomenclature we will use in this paper. The highest hierarchical level we use, or H1 type, divides neurons into excitatory neurons, inhibitory neurons, and other cells; this level is referred to as “class” level in many studies. Within each H1 type, we subdivide neurons into H2 types, which are sometimes referred to as “subclasses”(Yao et al. 2021a; Yao et al. 2021b). In particular, cortical excitatory neurons can be divided into nine H2 types that are shared across most cortical areas. This division refines the traditional projection-based IT (intra-telencephalic) /PT (pyramidal tract) /CT (corticothalamic) neuron (Harris and Shepherd 2015) classification as follows: PT and CT neurons correspond to L5 ET (extra-telencephalic neurons) and L6 CT neurons, respectively, whereas IT neurons are subdivided into L2/3 IT, L4/5 IT, L5 IT, L6 IT, NP (near-projecting neurons), Car3, and L6b. This division follows recent single-cell RNAseq studies but differs from the classical tripartite of IT/PT/CT neurons, which were defined largely based on long-range projections and failed to capture differences among some transcriptomically distinct cell types, such as NP, Car3, and L6b. Each H2 type can be further divided into H3 types, which correspond to “cluster” or “type” level in some studies (Yao et al. 2021a; Yao et al. 2021b). Previous studies have shown that H1 and H2 types are largely shared across most cortical areas, but the expression of many genes is localized to specific parts of the cortex both during development (O’Leary et al. 2007; Cadwell et al. 2019) and in the adult (Lein et al. 2007). Clusters at the H3 level appeared to be enriched in neurons dissected from different parts of the cortex in single-cell RNAseq studies (Tasic et al. 2018; Yao et al. 2021a). However, because these dissections could only be performed at a coarse resolution, the detailed distribution of neuronal populations at this higher granularity across cortical areas remains unclear.

To assess the distribution of neuronal populations across the cortex, we first generated a pilot dataset by applying BARseq to interrogate the expression of 104 cell type marker genes (**Supplementary Table 1**) in 40 hemi-brain coronal sections that cover the whole forebrain in one animal (**Fig. 1A, B**). We applied the same approach that we previously used to resolve cortical excitatory neuron types in the motor cortex (Sun et al. 2021). Briefly, we selected marker genes that were optimized for distinguishing excitatory neuronal types in the cortex (see **Supplementary Note 1**). We evaluated this gene panel using a comprehensive single-cell RNAseq dataset from the cortex (Yao et al. 2021b), and found that these genes performed similarly to the full transcriptome in distinguishing H2 (i.e. subclass-level) and H3 (i.e. cluster-level) types **(Fig. 1C; ED Fig. 1**; see **Supplementary Note 1**). We used up to 12 padlock probes to target each gene; each probe carried a 7-nt gene identification index (GII) that uniquely identified the gene. These GIIs were designed to have a minimum hamming distance of 3-nt to allow for error correction. We further included 5 blank GIIs that were not present in the padlock probes as negative controls when decoding the GIIs; these blank GIIs allowed us to control and evaluate false detection rates. We decoded the GIIs from seven rounds of sequencing using BarDensr (Chen et al. 2021a) while maintaining an optimal false detection rate (∼5% estimated using the blank GIIs). We registered our data to the Allen Mouse Brain Common Coordinate Framework v3 (CCF v3)(Wang et al. 2020) using a semi-manual procedure that utilized QuickNii, Visualign (Puchades et al. 2019), and custom python scripts (see **Data and code availability**). Three highly expressed high-level marker genes (*Slc17a7* for excitatory neurons, *Gad1* for inhibitory neurons, and *Slc30a3* for IT neurons) were detected by hybridization rather than by sequencing (**Fig. 1B**). We segmented cell bodies with Cellpose using DAPI as the nucleic channel in Cellpose and sequencing signals from all imaging channels as the cytoplasmic channel in Cellpose (Stringer et al. 2020) (**Fig. 1B**), which resulted in 2,167,762 cells. We then removed cells with insufficient number of reads (20 reads/cell and 5 genes/cell minimum), resulting in 1,259,256 cells after quality control (QC, see **Methods**) with a mean of 60 unique reads/cell and 27 genes/cell (**Fig. 1D, E**).

**Figure 1.**
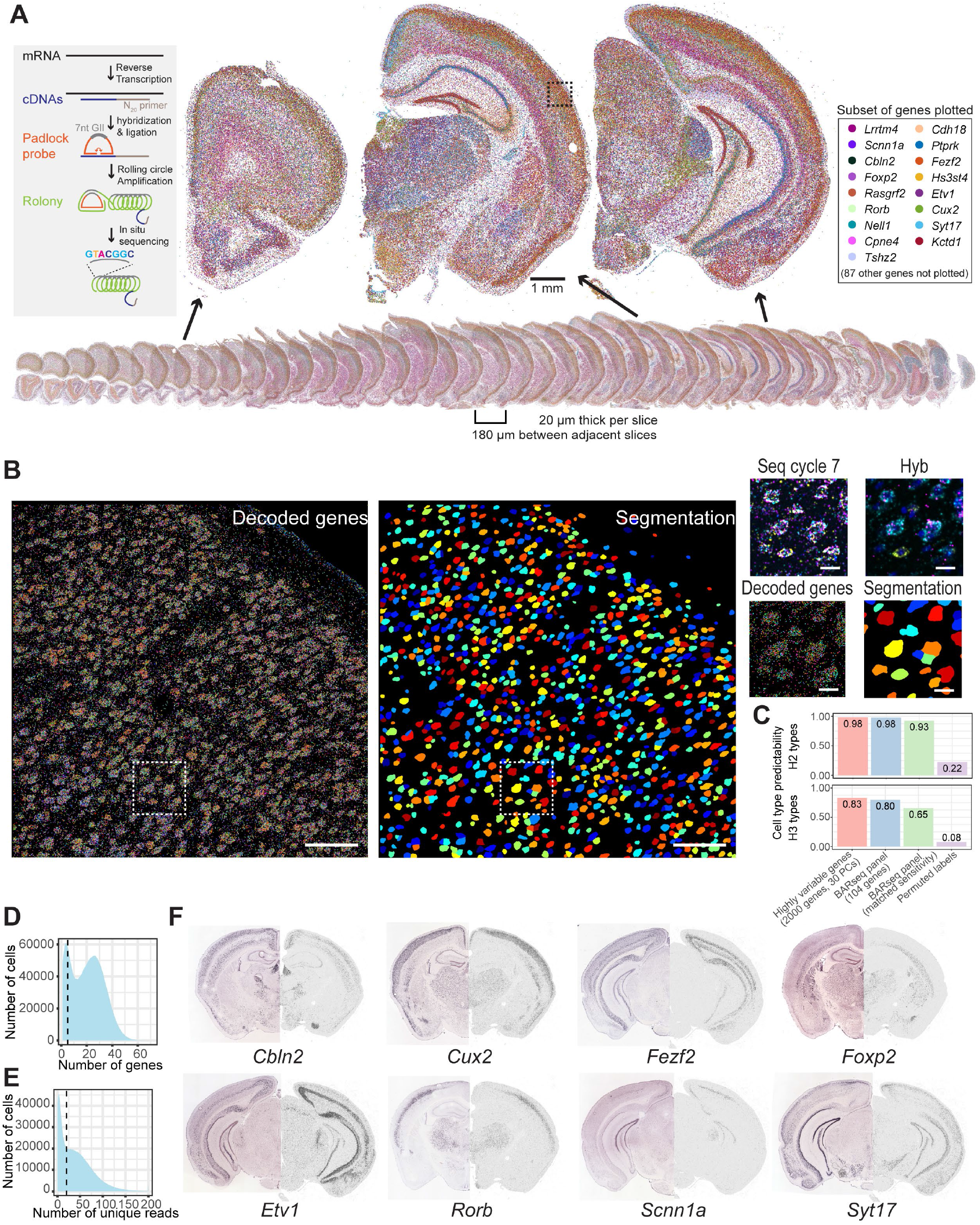
BARseq reveals brain-wide gene expression. (**A**) Images showing mRNA reads of all 40 slices (bottom) and close-up images of three representative slices (top). For clarity, only 17 out of 104 genes (indicated on the right) are plotted. Inset on the left shows illustration of mRNA detection using BARseq. Scale bar = 1 mm. (**B**) Decoded genes (left) and cell segmentations (middle) from a representative field-of-view that corresponds to the dashed box area in (A). Close up images of the dashed white box area showing the last sequencing cycle, hybridization cycle, decoded genes, and cell segmentation are shown on the right. Scale bars = 100 μm for full field-of-view images, and scale bars = 10 μm for the boxed area. (**C**) Single-cell cluster assignment performance using the full transcriptome, top principal components, and the 104-gene panel with or without subsampling to match sensitivity of BARseq for H2 (top) and H3 (bottom) clusters. (**D**) Gene counts per cell and (**E**) read counts per cell in the dataset. Quality control thresholds are indicated by dashed lines in both plots. The lower peaks in gene and read counts likely include non-neuronal cells that do not express the cortical neuronal markers in our gene panel and non-cellular particles that are fluorescent. (**F**) The expression patterns of representative genes in Allen Brain Atlas (left half) compared to the current dataset (right half).

At a gross anatomical level, many genes were differentially expressed across major brain structures and cortical layers (**Fig. 1A**). These expression patterns were consistent with the patterns of *in situ* hybridization in the Allen Brain Atlas (Lein et al. 2007)(**Fig. 1F**). For example, classical cortical layer-specific markers, including *Cux2*, *Fezf2*, and *Foxp2*, were expressed in layer 2/3, layer 5/layer 6, and layer 6, respectively. *Rorb*, a layer 4 marker, was seen in layer 4 throughout the cortex except in the motor cortex and medial areas, which lack a classically defined layer 4. *Scnn1a* was expressed mostly in the retrosplenial cortex and primary sensory areas, with the strongest expression in the primary visual cortex and primary somatosensory cortex. Thus, our pilot dataset recapitulated the known spatial distribution of gene expression.

### De novo clustering reveals neuronal subpopulations that are consistent with reference transcriptomic types

We next identified transcriptomic types of individual neurons based on single-cell gene expression. Generally, two approaches can be used to identify cell types in new transcriptomic datasets. In the first approach, we can map individual neurons in a new dataset directly to clusters in reference single-cell RNAseq datasets to determine cell type identities. This approach can match small datasets to cell types discovered in much larger, higher-resolution datasets (Cadwell et al. 2016; Bakken et al. 2021; Scala et al. 2021), but is prone to technique-specific variations when mapping data generated by different techniques (Liu et al. 2022). Alternatively, we can cluster the new dataset and map *clusters* to cell types in reference single-cell RNAseq datasets (Yao et al. 2021a; Zhang et al. 2021). This approach can better account for technique-specific variations and batch effects (Crow et al. 2018), but the ability to distinguish cell types is dependent on both data quality and sample size in the new dataset. Because our pilot dataset contained 1.2 million cells, which is comparable in size to many comprehensive single-cell RNAseq datasets (Saunders et al. 2018; Zeisel et al. 2018; Yao et al. 2021b), we reasoned that *de novo* clustering followed by assessment at the cluster level would be more easily interpretable.

We applied hierarchical clustering to separate neuronal populations at H1, H2, and H3 levels (**Fig. 2A**; see **Methods**). Clustering all cells resulted in 24 clusters, which we then combined into three H1 types (642,340 excitatory neurons, 427,939 inhibitory neurons, and 188,977 other cells) based on the expression of *Slc17a7* and *Gad1* (**ED Fig. 2A**). Of these 1.2 million cells, 517,428 were in the cortex and were the focus of the remainder of this study. Previous studies estimated the fraction of inhibitory neurons in the mouse cortex to be between 10% and 20% (Meyer et al. 2011; Sahara et al. 2012). Consistent with these estimates, 16% of neurons in the cortex were inhibitory neurons in our dataset (427,766 excitatory neurons, 83,394 inhibitory neurons, and 6,268 other cells). Because the excitatory and inhibitory neurons were defined by clustering on the expression of all genes, a small fraction of them did not have detectable *Slc17a7* (3,800 of 427,766 excitatory neurons, 0.9%) or *Gad1* expression (100 of 83,394 inhibitory neurons, 0.1%). Because we only sampled *Slc17a7* and *Gad1* for excitatory and inhibitory neuron markers, the excitatory neurons identified were dominated by cortical neurons, although we also saw neurons in the pons and the epithalamus in this group. The third group of cells, which expressed neither *Slc17a7* nor *Gad1*, included subpopulations of subcortical neurons (e.g., the midbrain and the thalamus) and non-neuronal cells (e.g., glial cells, immune cells, and epithelial cells). We expect this group of cells to be under-sampled, because these cells may not express cortical cell type marker genes that we probed at sufficient levels to pass quality control (**Fig. 1D, E**). Based on the fraction of excitatory neurons that express both *Slc17a7* and *Gad1*, we estimated that the probability of segmentation errors in which two neighboring cells were merged together, i.e. doublet rate, to be between 5% and 7% (**ED Fig. 2B, C**; see **Supplementary Note 2**). The 24 clusters, comprising the three H1 types, largely corresponded to coarse anatomical structures in the brain (**Fig. 2B**). For example, different clusters were enriched in the lateral group and ventral group of the thalamus, the intralaminar nuclei, the epithalamus, the medial, basolateral, and lateral nuclei of the amygdala, the striatum, and the globus pallidus (**Fig. 2B**). These results recapitulated the clear distinction of transcriptomic types across anatomically defined brain structures as observed in whole-brain single-cell RNAseq studies (Saunders et al. 2018; Zeisel et al. 2018; Langlieb et al. 2023; Yao et al. 2023; Zhang et al. 2023).

**Figure 2.**
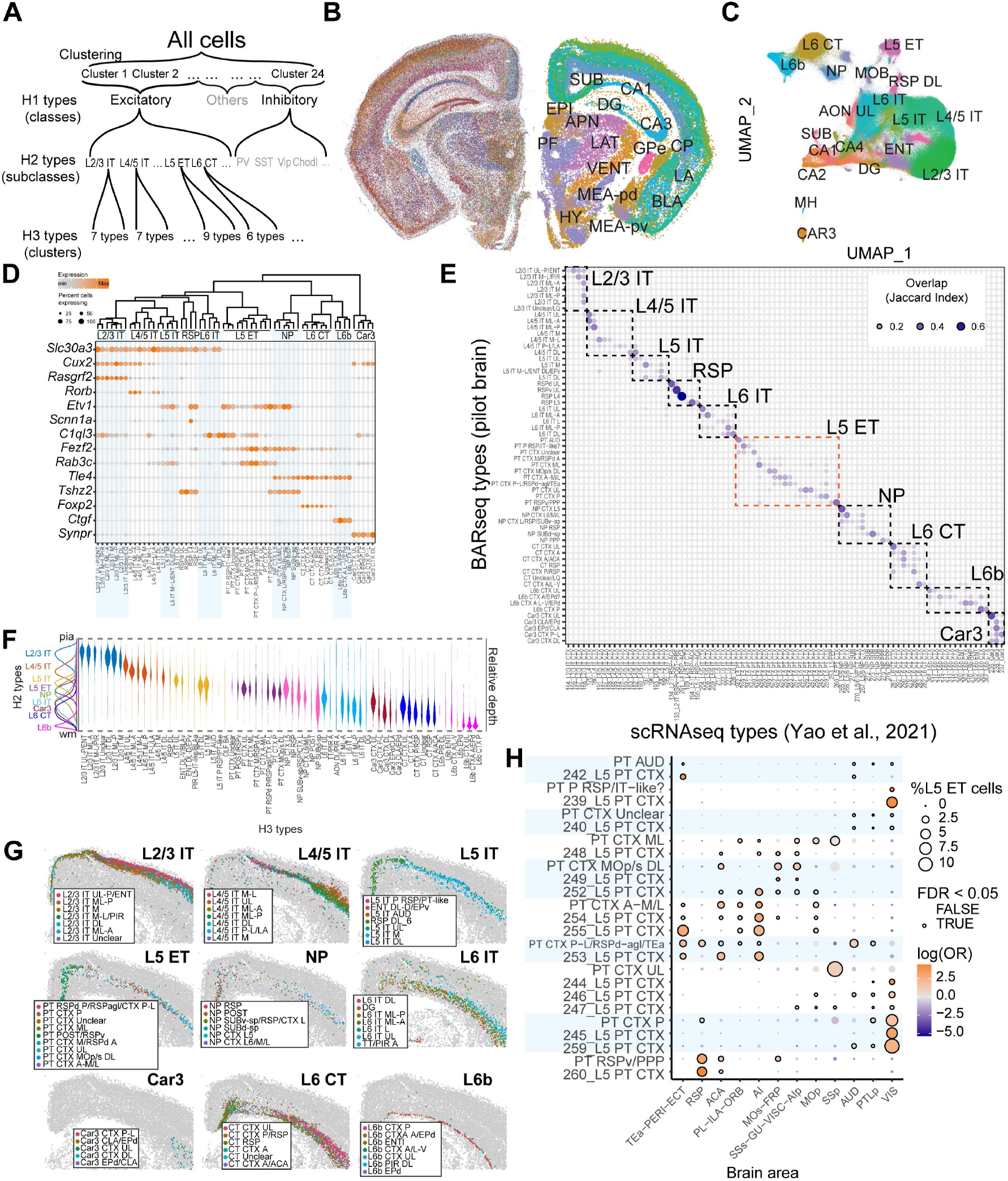
BARseq captures gene expression and spatial distribution of cortical excitatory cell types. (**A**) Workflow of hierarchical clustering. (**B**) Gene expression (left) and H1 clusters (right) in a representative slice. Major anatomical divisions that are distinguished by H1 clusters are labeled. SUB: subiculum, DG: dentate gyrus, CP: caudate putamen, GPe: Globus Pallidus, external segment, LA: lateral amygdala, BLA: basolateral amygdala, MEA-pd(pv): medial amygdalar nucleus, posterodorsal (posteroventral), EPI: epithalamus, APN: anterior pretectal nucleus, LAT: lateral group of the dorsal thalamus, VENT: ventral group of the dorsal thalamus, PF: parafascicular nucleus, HY: hypothalamus. (**C**) UMAP plot of gene expression of excitatory neurons colored by H2 types. (**D**) Marker gene expression in cortical excitatory H3 types. Colors indicate mean expression level and dot size indicates fraction of cells expressing the gene. The dendrogram on top shows hierarchical clustering of pooled gene expression within each H3 type. (**E**) Overlap (Jaccard index) between BARseq H3 types and scRNAseq cell types from Yao et al. (2021b). Dashed boxes indicate the parent H2 types. (**F**) The laminar distribution of H2 types (shown on the left) and each H3 type. H3 types are sorted by their median laminar position. (**G**) The distribution of H3 types in the dorsomedial portion of the cortex on a representative slice. The parent H2 types are indicated in each plot. (**H**) Distribution across CCF regions of matching L5 ET BARseq H3 types and L5 ET scRNAseq cell types (Jaccard Index > 0.1). Matched types are shown next to each other and share the same background color. The colors indicate log odds ratios and circle size indicates the fraction of cells among all L5 ET neurons.

We then re-clustered the excitatory and inhibitory neurons separately into H2 types (**Fig. 2C**; **ED Fig. 2D**) to improve the resolution of clustering. At this level, we recovered major inhibitory neuron subclasses (Pvalb, Sst, Vip/Sncg, Meis2-like, and Lamp5), all excitatory subclasses that are shared across the cortex (L2/3 IT, L4/5 IT, L5 IT, L6 IT, L5 ET, L6 CT, NP, Car3, L6b), and an excitatory subclass specific to the medial cortex (RSP) observed in previous cortical single-cell RNAseq datasets (Tasic et al. 2016; Tasic et al. 2018; Yao et al. 2021a; Yao et al. 2021b). The H2 types expressed known cell type markers and other highly differentially expressed genes (**Fig. 2D**). For example, *Cux2* is expressed mostly in superficial layer IT neurons and Car3 neurons; *Fezf2* is expressed in NP and L5 ET neurons; and *Foxp2* is expressed specifically in L6 CT neurons (see **Supplementary Note 3** for detailed description of all genes). Although we generated the full 40-section data in two batches (see **Methods**), we did not observe strong batch effects, because neurons from different slices across the two batches were intermingled in the UMAP plot for excitatory H2 types (**ED Fig. 2E**). Thus, the H2 types recapitulated known neuronal types at medium granularity that were identified in previous single-cell RNAseq datasets (Tasic et al. 2018; Yao et al. 2021a; Yao et al. 2021b).

We then re-clustered each excitatory H2 type into H3 types. To quantify how well H3 types corresponded to reference transcriptomic types identified in previous single-cell RNAseq studies, we used a k-nearest neighbor-based approach to match each H3 type to leaf-level clusters in a previous single-cell RNAseq dataset (Yao et al. 2021b) (see **Methods**). We found that cortical H2 types had a one-to-one correspondence to subclass-level cell types in the single-cell RNAseq data (**Fig. 2E**). Within each H2 type, the H3 types differentially mapped onto single or small subsets of leaf-level clusters in the single-cell RNAseq data (**Fig. 2E**; see **ED Fig. 2F** for matching of clusters outside of the cortex). These results demonstrate that our dataset resolved fine-grained neuronal subpopulations corresponding to clusters obtained previously using single-cell RNAseq data.

Both H2 types and H3 types were organized in an orderly fashion along the depth of the cortex. H2 types were concentrated in distinct layers, whereas H3 types within a H2 type were enriched in finer divisions within each layer (p < 1×10^-61^ using one-way ANOVA to compare the laminar positions of all H3 types within each H2 type after Bonferroni correction; **Fig. 2F**). For example, multiple H3 types of L2/3 IT, L4/5 IT, and L6 IT clearly occupied distinct sublayers in the somatosensory cortex (**Fig. 2G**). These results are consistent with previous studies using other spatial transcriptomic techniques (Codeluppi et al. 2018; Wang et al. 2018; Zhang et al. 2021) and with sublaminar differences in functional connectivity (Meng et al. 2017). Thus, our data recapitulated the laminar organization of cortical excitatory neurons.

Previous single-cell RNAseq studies, which used manual dissection to distinguish neurons from different cortical areas, showed that subpopulations of cortical neurons were also differentially distributed across large areas of the cortex (Tasic et al. 2018; Yao et al. 2021b). Consistent with these studies, H3 types and their matching clusters in single-cell RNAseq datasets were also found in similar cortical areas (**Fig. 2H**, **ED Fig. 2G, H, ED Fig. 3**). For example, the H3 type “PT AUD” and its corresponding single-cell RNAseq cluster (242_L5_PT CTX) were both enriched in the lateral cortical areas (TEa-PERI-ECT) and auditory cortex (AUD), whereas the H3 type “PT CTX P” and its corresponding single-cell RNAseq clusters (245_L5_PT CTX and 259_L5_PT CTX) were all enriched in the visual cortex (VIS). Thus, at this coarse spatial resolution, our data recapitulated previously observed areal distribution of cortical excitatory neurons.

To summarize, our pilot dataset resolved fine-grained transcriptomic types of cortical excitatory neurons that were consistent with previous single-cell RNAseq datasets (Yao et al. 2021b) and recapitulated their areal and laminar distribution (Tasic et al. 2018; Yao et al. 2021b; Zhang et al. 2021). Unlike these previous studies, the high resolution and the cortex-wide span of our dataset allow us to go beyond this coarse spatial resolution and resolve the spatial enrichment of gene expression and the distribution of neuronal subpopulations across the cortex at micron-level resolution. This high resolution matches those in previous connectivity studies (Zingg et al. 2014; Harris et al. 2019), and thus enables comparison of transcriptomic and connectomic organization of the cortex. In the next sections, we first examine how gene expression varies across the tangential plane of the cortex. We then examine the distribution of neuronal populations across cortical areas and identify modules of cortical areas with similar compositions of neuronal subpopulations. Finally, we explore how developmental processes contribute to the distinct transcriptomic identities of cortical areas and modules by collecting a larger dataset involving eight hemispheres of the forebrain from eight animals after neonatal enucleation.

### Gene expression varies within H2 types along interconnected cortical areas

Gene expression varies substantially across the whole cortex (Lein et al. 2007; Lu et al. 2021), but most cortical areas largely share the same H2 types, or subclasses, of excitatory neurons (Tasic et al. 2018; Yao et al. 2021b). Thus, it is unclear how differences in the organization of neuronal subpopulations lead to area-specific gene expression. Three sources of variation could contribute to the differences in gene expression across areas (**Fig. 3A**). First, the composition of H2 types may drive the differences in gene expression across the cortex (**Fig. 3Aa**, the cell type composition model). For example, the ratio of H2 type X to type Y might be high in visual cortex but low in motor cortex, so genes that are expressed more highly in X than in Y will be more highly expressed in visual cortex. Second, the expression of some genes may vary across space regardless of H2 type, i.e., they change consistently across space in multiple H2 types (**Fig. 3Ab**, the spatial gradient model). In this model, gene A may be more highly expressed in the visual cortex than in the motor cortex in both H2 types X and Y. Finally, the expression of some genes may vary across space in an H2-specific manner (**Fig. 3Ac**, the area-specialized cell type model). For example, gene A may be more highly expressed in the visual cortex than in the motor cortex in H2 type X. Our pilot dataset suggests that all three models contribute to the spatial variation of gene expression in the cortex, but the model that contributes most to the variation for each gene can vary.

**Figure 3.**
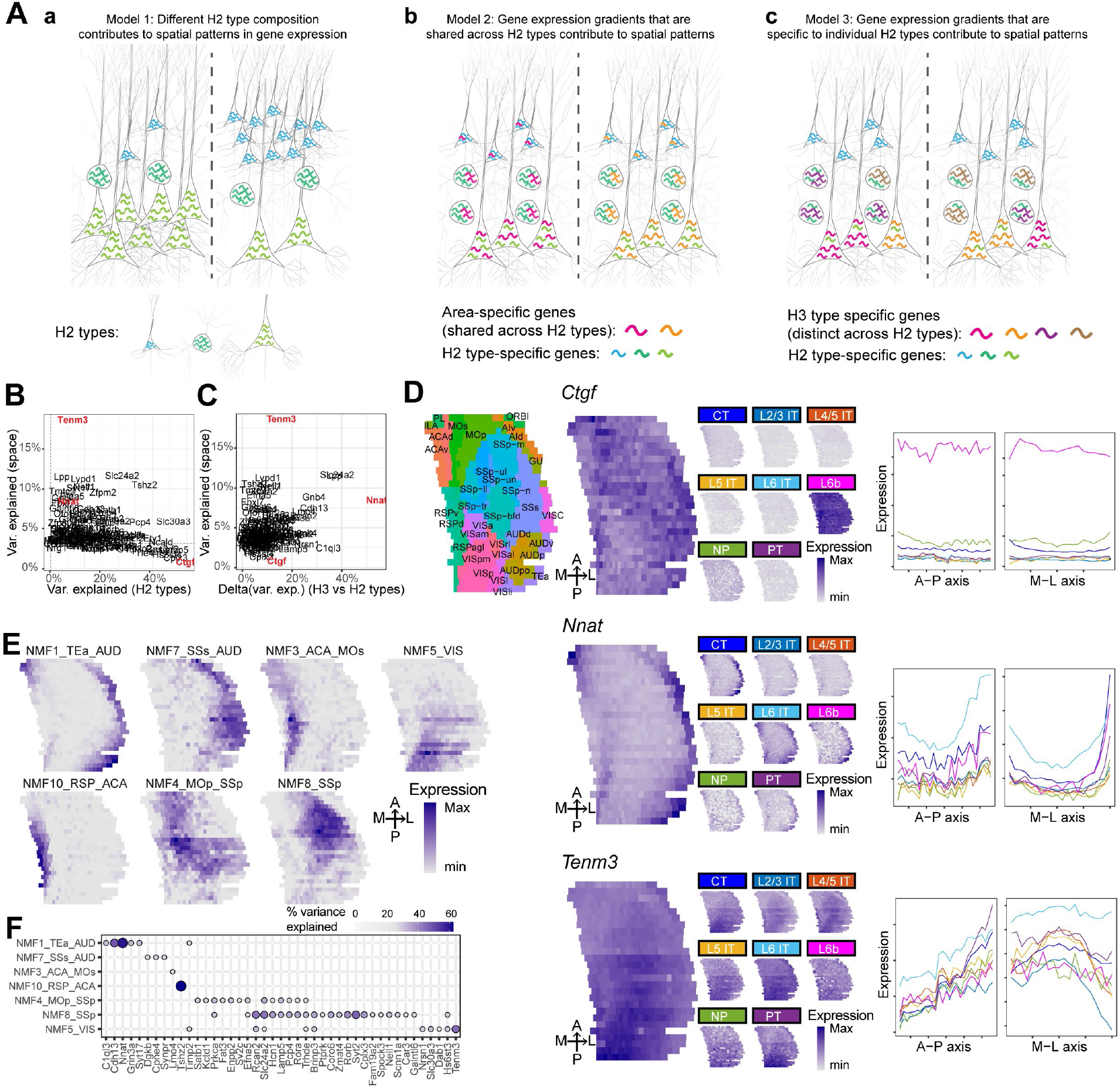
Spatial variations of gene expression across the cortex. (**A**)Three models of differential gene expression across cortical areas. (a) The relative proportion of each H2 type is different across areas. (b) The expression of genes varies across cortical areas consistently in different H2 types. (c) In each area, H2 types are enriched in different area-specialized H3 types. Different H2 types are indicated by neurons of different shapes. Cool-toned squiggles indicate H2 type-specific genes. Warm-toned squiggles indicate area-specific genes expression in (b) and H3 type-specific genes in (c). (**B**)(**C**) Variance in gene expression explained by space compared to that explained by H2 types (**B**) or additional variance explained by H3 types (**C**). In (B) dashed lines indicate threshold for p = 0.05 after Bonferroni correction. (**D**) The expression patterns of the indicated genes (Ctgf, Nnat, and Tenm3) plotted on flatmaps of the cortex in all cells (left column) or in each H2 type (center). The variations of gene expression in each H2 type along the AP axis and the ML axis are shown on the right. Line colors indicate H2 types as shown in the center plots. In the upper left plot, the same map is color-coded and labeled by cortical areas. (**E**) The expression of select spatially variant NMF modules plotted on cortical flatmaps. (**F**) The expression of select marker genes in each NMF module.

To determine the contribution of each source to the variation of gene expression across areas, we discretized the cortex on each coronal slice into 20 spatial bins. We “un-warped” the cortex on each slice (see **Methods** for details on unwarping), then divided the un-warped cortex into bins that spanned all cortical layers and had the same number of cells across bins in the same slice (**ED Fig. 4A**). We then assessed how much the variation in bulk gene expression across bins can be explained by space or by composition of H2 or H3 types using one-way ANOVA (**Fig. 3B, C**; see **Methods**). We found that variations in many genes were strongly explained by the composition of H2 types; these patterns were consistent with the cell type composition model (**Fig. 3Aa**). An example gene of this category is *Ctgf*, which is specifically expressed in L6b neurons at a consistent level across space (**Fig. 3D, *top***). Thus, variations in the expression (up to 80%) of *Ctgf* across space were largely explained by the fraction of L6b neurons in each bin rather than by variation in gene expression within cells. Other genes, in contrast, were largely explained by the composition of H3 types rather than the composition of H2 types (**Fig. 3C**, i.e. the area-specialized cell type model in **Fig. 3Ac**). Many genes in this category were also highly spatially variable (**Fig. 3C**, rho = 0.23), which suggest that H3 types were likely differentially distributed across the cortex (e.g. *Nnat*, marker of lateral areas, **Fig. 3D, *middle***). Finally, some genes displayed high spatial variability, but relatively low H2 and H3 variability. These genes were usually expressed in multiple H2 types (e.g. *Tenm3*, **Fig. 3D, *bottom***) and varied consistently in space across these H2 types, suggesting a general spatial gradient that is independent of H2 types. The spatial patterns of these genes were consistent with the spatial gradient model (**Fig. 3Ab**).

Because the spatial patterns of many genes were similar across genes and H2 types, we sought to extract basic spatial components that were shared across genes and H2 types using non-negative matrix factorization (NMF) (Lee and Seung 1999). Briefly, we performed NMF on the residuals of gene expression of spatial bins after accounting for differences in the composition of H2 types in each bin. We extracted ten NMF components, seven of which captured spatial variations in gene expression (the other three components captured slice-specific technical variability and were not used for subsequent analyses; see **Supplementary Note 4** and **ED Fig. 4B, C**). We found that the majority of NMF components were expressed not in broad gradients along major spatial axes, but rather in areas that were functionally related and highly interconnected (**Fig. 3E**; **ED Fig. 4D**). For example, NMF5 was expressed mostly in the visual areas, whereas NMF8 was expressed in somatosensory areas. Other NMF modules, including NMF1 (medial areas) and NMF10 (lateral areas), were expressed in combinations of areas that were functionally distinct but also highly interconnected (Zingg et al. 2014; Harris et al. 2019). Individual spatially variant genes were usually strongly associated with only one or two components (**Fig. 3F**; **ED Fig. 4E; ED Fig. 5**), and the association recapitulated known spatial patterns of these genes. For example, *Tenm3* was expressed mostly in posterior sensory areas, including the visual cortex, auditory cortex, and part of the somatosensory cortex (Lein et al. 2007)(**Fig. 3D, *bottom***); *Tenm3* was strongly associated with NMF5 (**Fig. 3F**), which was also expressed in the same sets of areas (**Fig. 3E**). Thus, the finding that gene expression varies along sets of interconnected areas suggests an intriguing link between gene expression and intra-cortical connectivity across areas.

**Figure 5.**
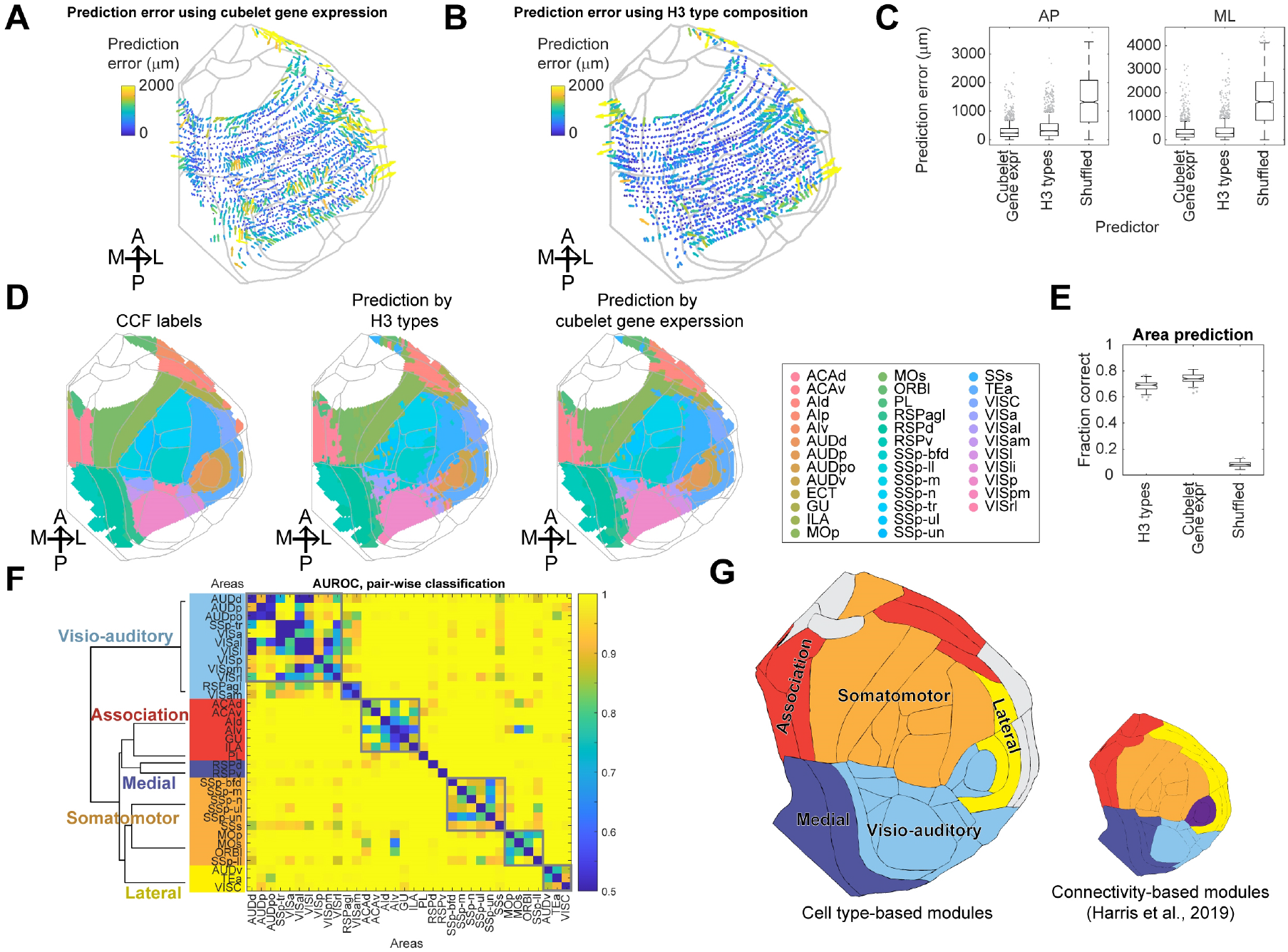
Modular and hierarchical organization of cortical areas by cell types. (**A**)(**B**) Heatmaps showing the errors in predicting cubelet locations using gene expression (**A**) or H3 type composition (**B**). Arrows indicate the directions of the errors and colors indicate the magnitudes of the prediction errors (in μm). The lengths of the arrows are proportional to the prediction error. (**C**) Box plots summarizing the prediction performance shown in (A) and (B). Boxes show median and quartiles and whiskers indicate range after excluding outliers. Dots indicate outliers. (**D**) Cortical areas defined in CCF (left) and those predicted by H3 types (center) and cubelet gene expression (right). (**E**) Fraction of correctly predicted cubelets using H3 type composition, cubelet gene expression, and shuffled control. (**F**) Matrix showing the AUROC of pair-wise classification between combinations of cortical areas. Areas are sorted by modules, which are color-coded on the left. Dendrogram is calculated using similarity of H3 type composition. Clusters obtained based on the matrix are shown in gray boxes. (**G**) Cortical flatmaps colored by cell type-based modules as defined in (F) (left) and by connectivity-based modules identified by Harris et al. (2019) (right). Areas in gray did not contain sufficient numbers of cubelets and were excluded from this analysis.

### Cortical areas have distinct compositional profiles of H3 types

Because the spatially varying NMF modules were obtained after controlling for variability in the composition of H2 types, but not H3 types, we hypothesized that these modules reflected differences in the composition of H3 types across cortical areas. Consistent with this hypothesis, each H3 type was enriched in a small subset of NMF modules **(Fig. 4A**; see **Methods**). All H2 types contained at least one H3 type that was associated with NMF modules that were expressed in the medial and lateral areas (NMF 1, 3, 7, 10), and one to four H3 types that were associated with NMF modules that were expressed in subsets of the dorsal cortex, including the motor, somatosensory, and visual areas (NMF 4, 5, 8). The associations between H3 types and the NMF modules were different across H2 types, suggesting that different H2 types were specialized to different degrees at the H3 level. For example, the H3 types of L5 ET neurons had strong associations with individual NMFs, whereas H3 types of NP and L6b neurons showed little specialization within the dorsal cortex.

**Figure 4.**
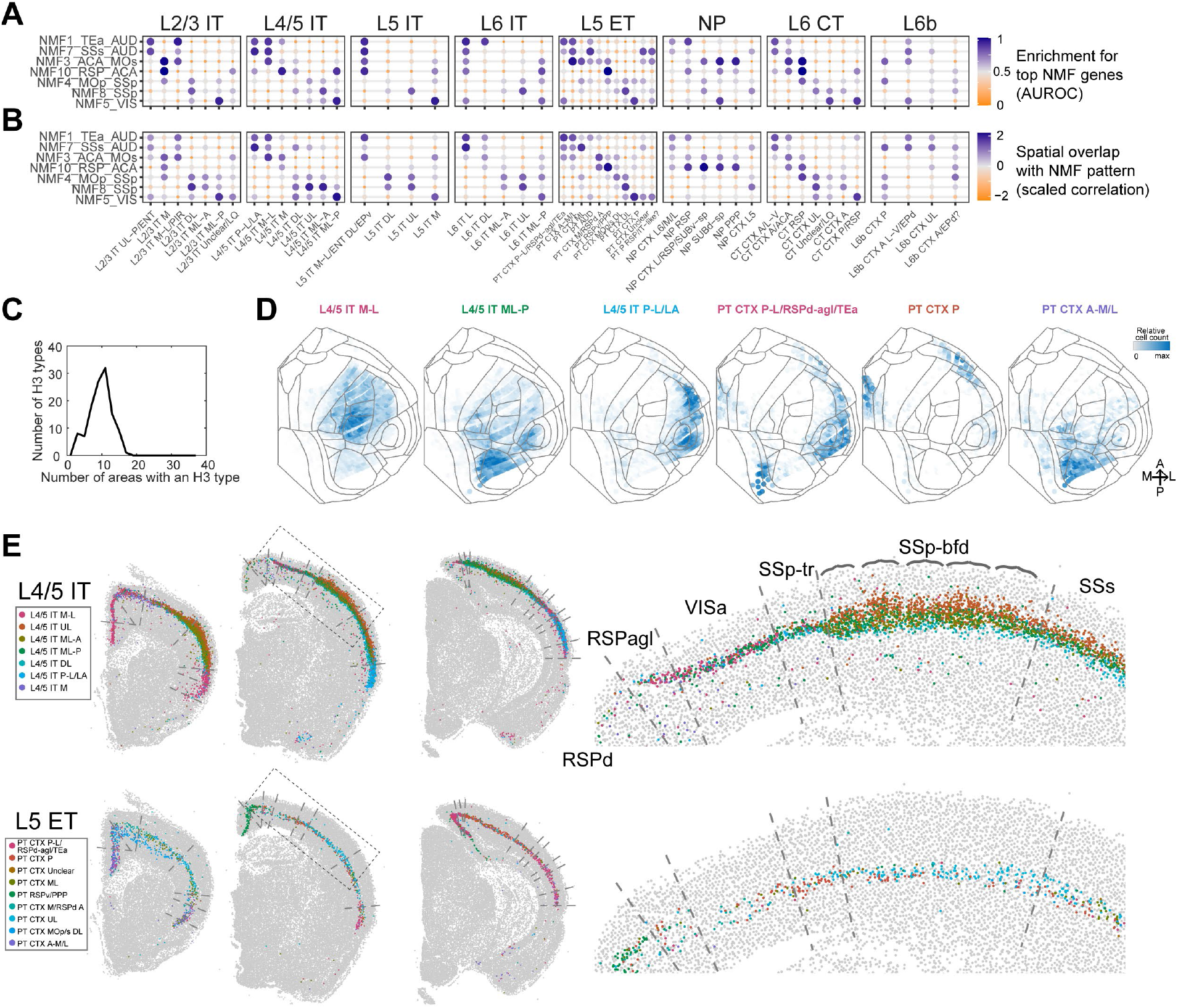
The composition of cell types is distinct across cortical areas. (**A**) AUROC of the enrichment for top NMF genes in each H3 type (see **Methods**). (**B**) Overlap between the spatial patterns of NMF module expression and the spatial distribution of H3 types. (**C**) Histogram showing the number of areas (out of 37 areas) that an H3 type was in. An H3 type was considered present in an area if that area contained at least 3% of that H3 type. (**D**) The spatial distribution of example L4/5 IT and L5 ET H3 types across the cortex plotted on cortical flatmaps. Colors indicate relative cell counts in each cubelet. Gray lines delineate cortical areas. (**E**) The distribution of L4/5 IT and L5 ET H3 types in example coronal sections. Dashed lines indicate area borders in CCF. Magnified views of the dashed boxes are shown on the right. Brackets in the top plot indicate barrels in the barrel cortex.

Consistent with the association in gene expression between NMF modules and H3 types, the H3 types also overlapped with their corresponding NMF modules in space (**Fig. 4B**; **ED Fig. 6**; see **Methods**). For example, L4/5 IT ML-P was associated with NMF5_VIS and was enriched most strongly in visual cortex; similarly, L4/5 IT P-L/LA was associated with NMF1_TEa_AUD and NMF7_SSs_AUD and was most highly enriched in auditory cortex and temporal association areas. Consistent with the expression patterns of NMF modules, many of these sets of areas were largely within cortical modules that were defined by inter-connectivity (Zingg et al. 2014; Harris et al. 2019). Thus, H3 types are associated with spatial gene co-expression modules and, at a coarse spatial resolution, are enriched in combinations of cortical areas that are highly interconnected.

To further assess the areal distribution of H3 types, we discretized the cortex on each coronal slice into “cubelets” with similar widths along the mediolateral axis across all slices. Each cubelet spanned all cortical layers and was about 100 μm to 200 μm wide along the curvature of the cortex, and 20 μm thick along the A-P axis (i.e. the thickness of each coronal section; see **Methods**; **ED Fig. 4A**). These cubelets were of similar physical sizes and were narrower on the mediolateral axis than the spatial bins used in the previous analysis; this higher lateral resolution makes it easier to assign cubelets to individual cortical areas. We found that H3 types were shared by multiple cortical areas and not specific to any single area (each H3 type was found in 9 ± 3 areas, median ± standard deviation; **Fig. 4C, D**; **Supp. Fig. 1**). Thus, the distinctness of neighboring cortical areas cannot be explained simply by the presence or absence of an area-specific H3 type. However, we noticed that the compositional profiles of H3 types often changed abruptly near area borders defined in CCF (**Fig. 4E; ED Fig. 7A**). Most salient changes occurred at the lateral and medial areas, which are consistent with single-cell RNAseq data (Yao et al. 2021b). Within the dorsolateral cortex, although neighboring cortical areas sometimes shared sets of H3 types, their compositions usually changed abruptly at or near area borders. For example, three L4/5 IT types, including L4/5 IT UL, L4/5 IT ML-P, and L4/5 IT DL, were found in three adjacent somatosensory areas (trunk area, SSp-tr; barrel cortex, SSp-bfd; secondary somatosensory cortex, SSs; **Fig. 4E, *top row***), but the numbers of neurons of each type were distinct across the three areas. L4/5 IT UL was found in only small numbers in SSp-tr but expanded in both the number of neurons and their laminar span in the barrel cortex. Furthermore, L4/5 IT UL neurons were clustered along the mediolateral axis into structures that resembled barrels. In the secondary somatosensory cortex, the distribution of L4/5 IT UL lost the barrel-like patterns but remained present in substantial numbers. In contrast, L4/5 IT DL became more dominant relative to L4/5 IT ML-P. Similarly, in L5 ET neurons, PT CTX P was more dominant in SSp-tr, whereas PT CTX ML was found mostly in SSs (**Fig. 4E, *bottom row***). To quantify how well abrupt changes in H3 type composition corresponded to borders between all cortical areas, we identified positions where H3 type composition changed abruptly by identifying peaks in the absolute value of the first derivatives of H3 type composition along the ML axis within each slice. Consistent with the impression from images of coronal sections, 53% of peaks in the first derivatives were within 150 µm from the closest CCF border (we could not assess matching using a more stringent distance because 150 µm is already comparable to the cubelet size in this dataset). The fraction of peaks that were close to CCF borders was higher than 99% of shuffled controls (**ED Fig. 7A**). These results reconcile two seemingly contradictory observations in previous single-cell RNAseq studies: distant cortical areas (e.g. visual cortex and anterolateral motor cortex) have distinct excitatory cell types (Tasic et al. 2018), but individual cell types are usually found across large areas of the cortex (Yao et al. 2021b). Thus, cortical areas are largely distinct in their compositional profiles of H3 types, but individual H3 types are rarely specific to a single cortical area.

### H3 type composition reveals modular organization of the cortex

Because variation in gene expression and H3 types both followed spatial patterns that were reminiscent of highly interconnected cortical areas, we hypothesized that, like the modular organization of corticocortical connectivity, cortical areas were also organized into modules based on H3 types: cortical areas within a module would be composed of similar sets of H3 types, whereas areas in different modules would be more distinct in their compositions of H3 types. To test this hypothesis, we first asked how well cortical areas could be distinguished using the compositions of H3 types. We then identified cortical modules using the differences in H3 type compositions.

To test how well the compositions of H3 types could predict cortical areas, we first used random forest classifiers to predict the AP and ML coordinates of each cubelet given either the total gene expression in that cubelet (**Fig. 5A**) or its H3 type composition (i.e. the fraction of each H3 type within a cubelet; **Fig. 5B**). We found that cubelet gene expression was highly predictive of locations in the cortex, capturing 94% of variance on both the AP and ML axes. The distance between the predicted and true location of a cubelet across the whole cortex was 235 ± 270 μm (median ± std) along the AP axis (spanning 5,900 μm) and 245 ± 364 μm along the ML axis (spanning 8,400 μm) (**Fig. 5A, C**). These prediction errors were close to the sampling frequency imposed by cubelet size (200 μm between adjacent slices on the AP axis, and 100 μm to 200 μm cubelet width on the ML axis). Strikingly, the H3 type compositions performed similarly well, capturing 89% variance on the AP axis and 92% variance on the ML axis (the prediction errors were 312 ± 360 μm on the AP axis and 269 ± 402 μm on the ML axis, median ± std; **Fig. 5B, C**). Consistent with the high precision in predicting the absolute locations in the cortex, both gene expression and the composition of H3 types were highly predictive of the area labels in CCF (75% correct using gene expression and 69% correct using H3 type composition, compared to 8% in shuffled control; **Fig. 5D, E**). H3 types within a single parent H2 type were also somewhat predictive of cubelet locations, but those of H2 types in superficial layers (e.g. L2/3 IT and L4/5 IT) were generally more predictive of cubelet locations along the ML axis than those of H2 types in the deep layers (e.g. L6 IT and L6 CT) (**ED Fig. 7B, C**). The predicted maps correctly captured the locations of most cortical areas, and most of the incorrect predictions occurred along the borders of areas (**Fig. 5D**). Thus, both cubelet gene expression and H3 type compositions are highly predictive of the locations along the tangential plane of the cortex and the identity of the cortical areas.

We next assessed the similarity and modularity of cortical areas based on how well cortical areas could be distinguished by their H3 type compositions (**Fig. 5F**; see **Methods**). Briefly, we built a distance matrix between cortical areas based on how well they can be distinguished pairwise using H3 type composition, then performed Louvain clustering on the distance matrix. We identified six clusters, each of which consisted of more than one area (**Fig. 5F**, gray boxes); these included two clusters that corresponded to the visio-auditory areas and one cluster each for the association areas, somatosensory cortex, motor cortex, and lateral areas. This modular organization is robust to small errors in CCF registration (**ED Fig. 7D**; see **Methods**). We further combined these clusters with areas that did not cluster with other areas (PL, RSPd, RSPv) into cortical modules based on similarity in H3 type composition. These modules largely included the visio-auditory areas, somatomotor areas, the association areas, medial areas, lateral areas, respectively (**Fig. 5F**). Strikingly, these modules were largely consistent with cortical modules that are highly connected (connectivity-based modules) (Harris et al. 2019) (**Fig. 5G**). Thus, highly interconnected cortical areas also share similar H3 types and, consequently, characteristic transcriptomic signatures.

### The H3 type compositional profiles reveal neonatal refinement by peripheral inputs

Transcriptomic types, areas, and modules reflect cortical organization at different scales, which suggest that they may be generated through different developmental mechanisms. As a first step in understanding the developmental processes that contribute to cortical organization at different scales, we applied BARseq to examine how postnatal removal of peripheral sensory inputs altered the organization of cortical transcriptomic types. Thalamocortical projections play a central role in shaping the identities and borders of cortical areas (Chou et al. 2013; Pouchelon et al. 2014; Cadwell et al. 2019), and loss of postnatal visual inputs affects gene expression in VISp and other areas (Dye et al. 2012; Chou et al. 2013; Cheng et al. 2022). How peripheral inputs shape cortical neuronal types and the characteristic cell type compositional profiles of cortical areas, however, is unclear. For example, this altered gene expression could result in new cell types that are not seen in a normal brain; alternatively, it could enrich or deplete existing cell types (**Fig. 6A**). Because BARseq is cost-effective and has high throughput, it is uniquely suited for interrogating changes in neuronal gene expression and cell type compositional profiles on a brain-wide scale across many animals, with or without developmental perturbation.

**Figure 6.**
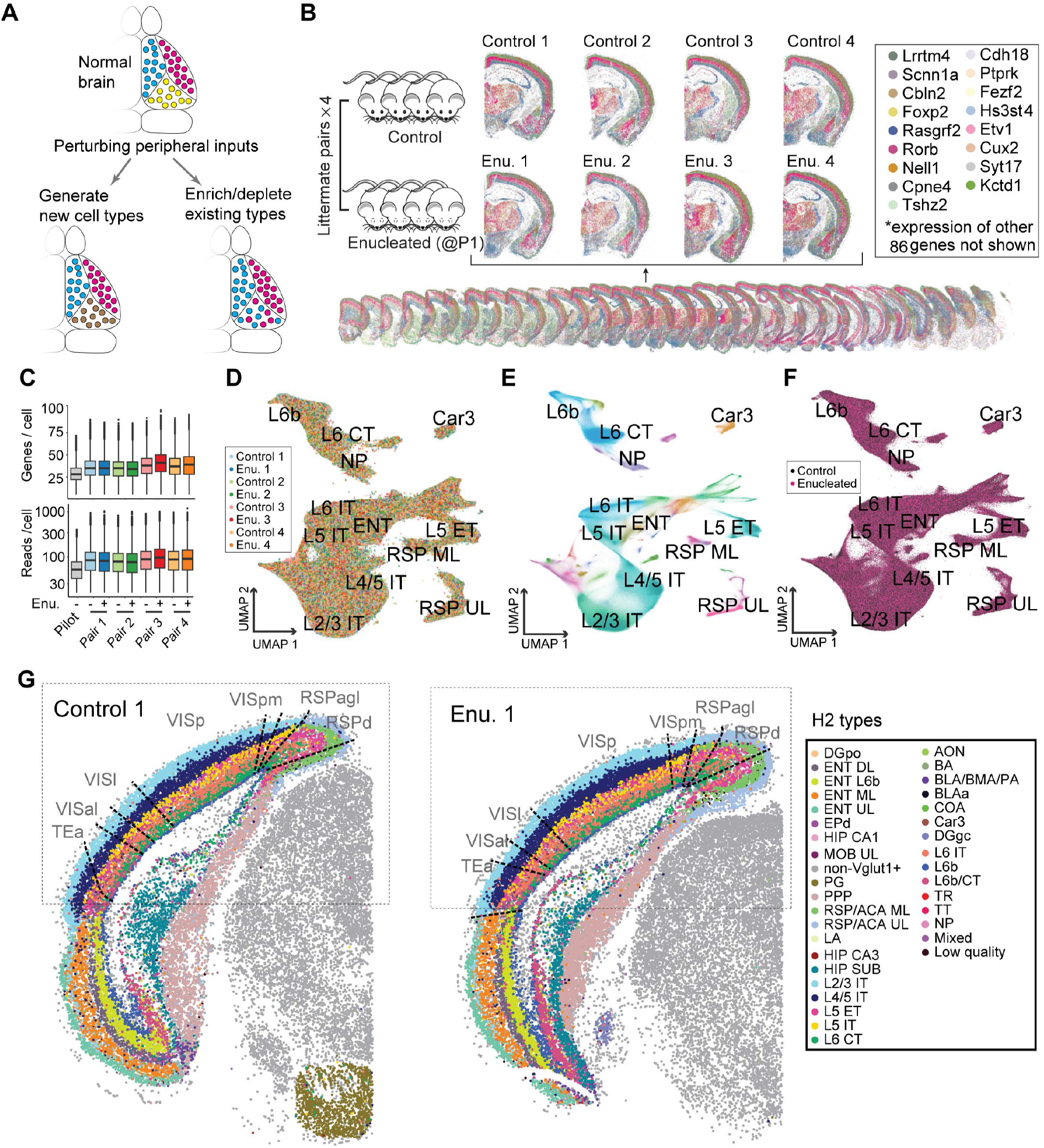
BARseq consistently detects cell types across eight enucleated and control animals. (**A**) Models of possible effects of removing peripheral sensory inputs postnatally, including generating new cell types (left), and/or enrichment and depletion of existing cell types (right). (**B**) We collected brain-wide transcriptomic data from four littermate pairs: within each pair one mouse was enucleated at P1 and the other was a sham control (left, N=8 total animals). A representative stack of 32 slices from one brain (bottom) and close-up images of matching coronal slices from all eight brains (top) are shown. For clarity, only 17 out of 104 genes (indicated on the right) are plotted. (**C**) Genes per cell and read counts per cell for the pilot dataset and the enucleation and control littermates. (**D**)(**E**)(**F**) UMAP plots of gene expression of excitatory neurons from all eight animals. Neurons are color coded by animals (D), by H2 types (E), or by conditions (enucleated or control, F). Labels show only H2 types in cortex. (**G**) Representative slices from a littermate pair color coded by H2 types. Enu, enucleated.

We performed binocular enucleation on four mice at postnatal day 1 and collected their brains at postnatal day 28 along with four matched littermate controls (*N* = 8 total animals) (**Fig. 6B**). We then performed BARseq using a high-throughput *in situ* sequencing system with improved lasers and cameras and larger field-of-views (see **Methods**), which allowed us to sequence the 104-gene panel in 32 hemi-brain coronal sections per mouse, covering most of the forebrain of all eight animals (**Fig. 6B**) in 18 days (i.e., 2.3 days per brain). The *in situ* sequencing data were processed, segmented, registered to Allen CCF v3 (Wang et al. 2020), and quality controlled using the same approach as the pilot brain. The full dataset contained 9.1 million cells after quality control with a median of 87 reads per cell and 37 genes per cell (**Fig. 6C**). These quality control measures were significantly improved from the pilot dataset, likely due to improvements in the microscope and in data processing (see **Supplementary Note 5** for discussion). Cells from individual brains were interdigitated with cells from other brains in UMAP space, suggesting that there were minimal batch effects (**Fig. 6D**; **ED Fig. 8A**). Therefore, we performed *de novo* clustering hierarchically on the concatenated data into 3,957,252 excitatory neurons, 1,526,182 inhibitory neurons, and 3,635,402 other cells at the H1 level. The fraction of other cells was significantly higher than that in the pilot dataset, likely because the improved data quality allowed more cells with lower read counts to pass QC and be included. We then re-clustered the excitatory neurons into 35 H2 types (**Fig. 6E**) and 154 H3 types, including 12 H2 types and 70 H3 types that were predominantly found in the cortex. These H3 types in the new dataset closely matched those in the pilot dataset (**ED Fig. 8B, C**; see **Supplementary Note 5** for mapping to the pilot dataset).

We first examined whether the effect of enucleation was predominantly reflected in enrichment and/or depletion of H3 types that were already present in the control brains, or distinct H3 types that were absent in the control brains. Consistent with the lack of batch effect seen in the UMAP plots (**Fig. 6D**), all four pairs of littermates had similar fractions of H1, H2, and cortical H3 types (**ED Fig 8D-F**). Strikingly, no H3 type was strongly enriched in either the control brains or the enucleated brains (**ED Fig 8F**). The lack of condition-specific H3 types can be visualized in UMAP plots, in which neurons color-coded by their conditions were smoothly intermixed together (**Fig. 6F**). This lack of new cell types is also reflected in the spatial distribution of H2 types, which were visually similar across the two conditions (**Fig. 6G**; but see below for the distribution of H3 types). To further test the hypothesis that H3 types were shared by both the control and the enucleated brains, we trained a nearest neighbor classifier to predict whether a neuron is from a control or an enucleated brain. Briefly, for each neuron from a held-out litter, we found the 100 nearest neighbor neurons in transcriptomic space across the other three litters and predicted whether the neuron was from a control or an enucleated brain based on the number of control neighbors (see **Methods**). If an H3 type or a subpopulation of an H3 type is seen only in the enucleated brain, then we would expect that neurons in that neighborhood to be surrounded by mostly neurons from the enucleated brains, thus resulting in a high AUROC score. In contrast, the classifier largely failed to predict whether a neuron was from a control or an enucleated brain (mean AUROC was 0.49), although we saw subtle secondary shifts in gene expression within some H3 types in visual areas (see **Supplementary Note 6**; **ED Fig. 8G, H**). Although we cannot fully rule out the possibility that minor changes in gene expression were missed at our transcriptomic resolution, these results suggest that enucleation did not lead to the creation of new cell types at the H3 level; rather, the main effect of enucleation was likely reflected in changes in the compositional profiles of H3 types.

### Enucleation enriches medial and lateral cell types in the visual areas

Having established that enucleation did not create new H3 types, we sought to characterize enucleation-induced changes in area-specific H3-type composition. We divided the cortex into cubelets using a similar approach as we used for the pilot data (see **Methods**). This discretization resulted in about 270 neurons per cubelet, with a mean distance between adjacent cubelets on a section of 181 µm. To visualize the H3 type composition, we plotted UMAP plots based on the fraction of H3 types in each cubelet (**Fig. 7A-C**). Consistent with the lack of batch effect seen in the single-neuron gene expression (**Fig. 6D**), cubelets from all eight animals mixed smoothly in most areas (**Fig. 7A**). Color-coding cubelets by conditions (**Fig. 7B**), however, revealed an “island” (*left*) within which cubelets from the two populations (enucleated vs. control) were largely segregated. This island contained mainly cubelets from VISp and other visual areas (**Fig. 7B, C, insets**). To quantify the differences in the compositional profiles of H3 types between the control and the enucleated brains, we trained a classifier to assess how distinct cubelets from each cortical area were between the two conditions. If enucleation consistently altered the compositional profile of H3 types in a cortical area, then we would expect that the classifier should be able to predict whether a cubelet was from a control animal or an enucleated animal based on its H3 type composition above chance level. For each cubelet, we identified the 10 nearest neighbor cubelets from the same cortical area in the six animals that came from different litters, then trained a classifier to predict the condition of the cubelet based on the conditions of the neighbors; we then compared the accuracy of prediction to a classifier trained on data with the brain conditions shuffled (see **Methods**; **Fig. 7D**). In most cortical areas, the classifier performed at chance level, but VISp cubelets were highly predictive of condition (**Fig. 7D**; AUROC 0.90 ± 0.06 compared to shuffled AUROC 0.56 ± 0.27, median ± standard deviation; p = 2 × 10^-33^ using rank sum test and Bonferroni correction). Two higher visual areas (VISpm and VISl; AUROC medians ± standard deviations are 0.70 ± 0.12 and 0.66 ± 0.15, respectively and the shuffled AUROC medians ± standard deviations are 0.50 ± 0.25 and 0.44 ± 0.25; p = 3 × 10^-9^ and 7 × 10^-9^, respectively, comparing each area to shuffled control using rank sum test and Bonferroni correction) and a non-visual area (SSp-ll; AUROC 0.57 ± 0.09 and shuffled AUROC 0.42 ± 0.18, median ± standard deviation; p = 2 × 10^-8^ comparing to shuffled control using rank sum test and Bonferroni correction) were also predictive above chance level, although the predictive powers were much lower. Thus, enucleation largely affected the relative composition of H3 types within visual areas.

**Figure 7.**
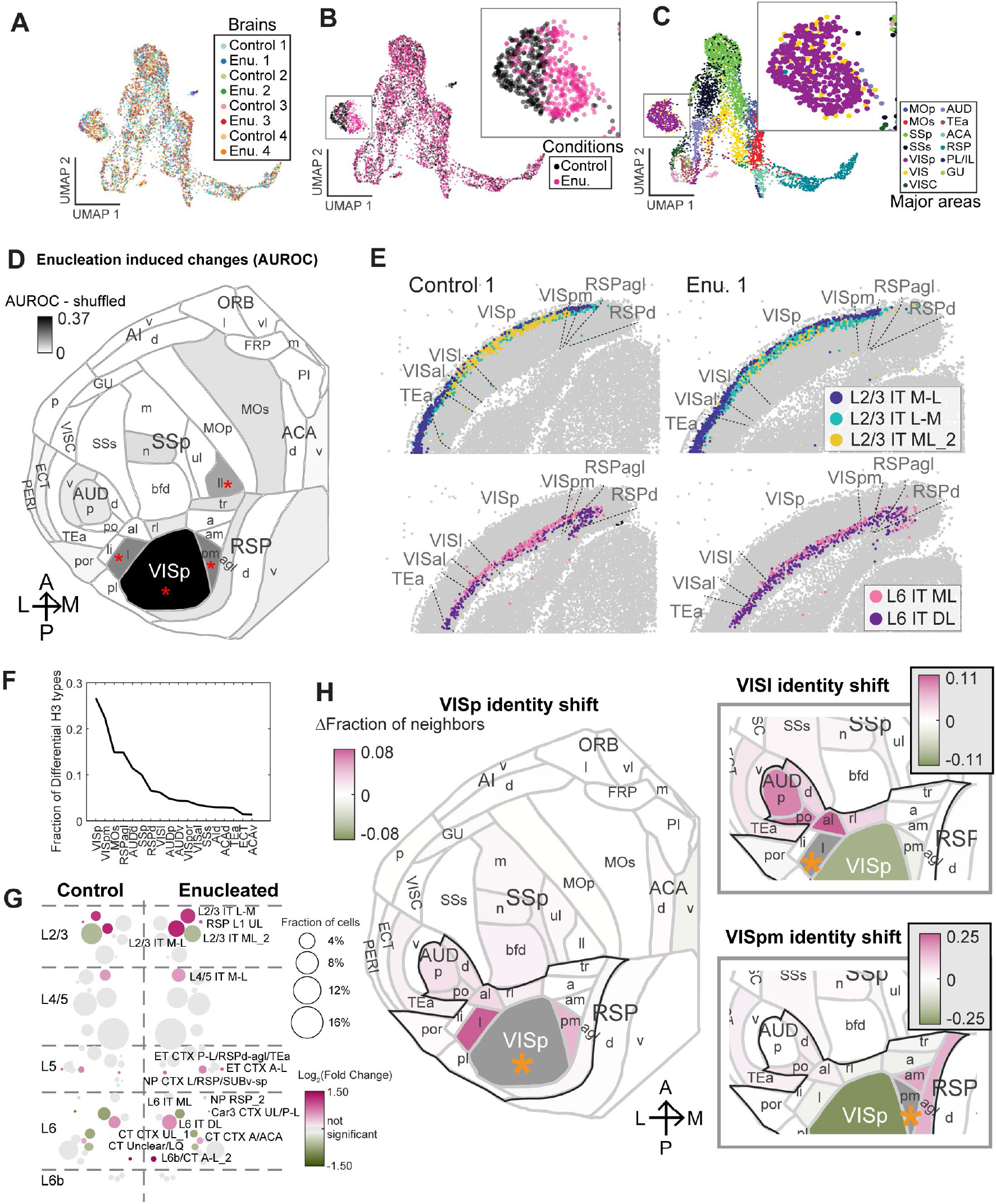
Enucleation shifts visual area identities within the visio-auditory module. (**A**)-(**C**) UMAP plots of the H3 type compositions of cubelets from all eight brains, color coded by animals (A), experimental conditions (B), or cortical areas (C). Insets show amplified views of the area corresponding to VISp cubelets. (**D**)Flatmap showing how distinct an area was between the control brains and the enucleated brains, as measured by AUROC. Colors indicate the differences in AUROC scores for the data and the AUROC scores after shuffling the conditions of the brains. Red asterisks indicate FDR < 0.05. (**E**) Same representative slices as in Fig. 6H, showing select L2/3 IT and L6 IT H3 types that were enriched or depleted in VISp after enucleation. The areas shown correspond to the dashed boxes in Fig. 6H. (**F**) The fractions of enriched or depleted H3 types in each area. (**G**) The fractions of H3 types in VISp and their fold changes after enucleation. Colors indicate log fold change and circle size indicates the fraction of VISp neurons that belonged to each H3 type. The names of H3 types that were enriched or depleted are labeled next to the dots. (**H**) Shifts in areal identity for VISp (left), VISl (top right), and VISpm (bottom right). Colors indicate fraction of enriched or depleted neighbors in enucleated brains compared to the control brains. The area of interest in each plot is shaded in gray and indicated by an orange asterisk. Areas outlined in black indicate the visio-auditory module as defined in Fig. 5. Enu, enucleated.

The effect of enucleation can be directly observed in the distribution of H3 types in the primary visual area (**Fig. 7E; ED Fig. 9A**). For example, many L2/3 IT ML_2 neurons (**Fig. 7E**, yellow dots) were found in VISp in control animals, but L2/3 IT L-M neurons (**Fig. 7E**, green dots) became enriched in VISp in the enucleated animals. Similarly, L6 IT DL neurons were seen in higher numbers in the VISp of the enucleated animals compared to that of the control animals. To systematically examine how enucleation affected the compositional profiles of cortical excitatory cell types in each area, we looked for H3 types that were enriched or depleted in enucleated brains using an ANOVA model, adjusting for litter and area effects (see **Methods**). We found that 46 H3 types in 18 areas across the whole cortex (**Fig. 7F**) were over- or under-represented in the enucleated animals compared to control animals. VISp had the most H3 types (16 types) whose compositions were altered by enucleation. The affected H3 types were found across most H2 types, with the strongest enrichment or depletion of H3 types of L2/3 IT, L4/5 IT, and L6 IT (**Fig. 7G**). Intriguingly, L6b/CT A-L_2, a transitional type between L6 CT and L6b H2 types that was usually found only in lateral areas, was also highly enriched in VISp after enucleation. The affected H3 types remained in their characteristic sublaminar positions (**ED Fig. 9B)**, and the overall changes were consistent with, but broader than those observed after dark rearing during the critical period (Cheng et al. 2022) (**ED Fig. 9C**; see **Supplementary Note 7**). The top enriched types, including L2/3 IT M-L, L2/3 IT L-M, L4/5 IT M-L, L6 IT DL, and L6b/CT A-L_2, were all enriched in medial and lateral areas in the control brains, including areas that were immediately medial and lateral to the visual areas (**ED Fig. 9D**). Thus, enucleation broadly shifted neurons in VISp towards H3 types that were usually enriched in the medial and lateral areas in control brains.

### Peripheral inputs sculpt characteristic cell type compositions of the visual areas within the visio-auditory module

Because the enriched H3 types were consistently found in medial and/or lateral areas, we wondered if these changes also shifted the overall area identity – as defined by the H3 type compositional profiles – of the visual cortex towards other areas. To examine how area identities changed after enucleation, we used a nearest-neighbor based approach that was inspired by MetaNeighbor (Crow et al. 2018) to assess how similar cubelets in the control and the enucleated brains were to other cubelets in control brains (see **Methods**). If enucleation shifted the compositional profile of an area towards a target area, then cubelets from that area in the enucleated brain would have more neighbors in the target area than cubelets from the same area in the control brain. For each cubelet in a littermate pair, we found the 20 nearest neighbor cubelets in H3 type composition in the control brains of the other three pairs of littermates. We then calculated the similarity, quantified by the AUROC for assigning cubelets from each area to areas in the control brains based on the nearest neighbors (**ED Fig. 9E**). All three visual areas (VISp, VISl, and VISpm, circled in **ED Fig. 9E**) remained highly similar to the same areas in control brains (AUROC 0.97 and 0.98 for control and enucleated VISp, 0.90 and 0.92 for control and enucleated VISl, and 0.88 and 0.94 for control and enucleated VISpm), indicating that their H3 type compositions remained highly distinct to other areas despite the changes induced by enucleation. However, each affected area also shifted towards the identities of neighboring areas as judged from the fraction of neighbors from an area (**Fig. 7H**). For example, VISp cubelets from the enucleated brains had higher AUROC scores with both VISl and VISpm than VISp cubelets from the control brains (0.85 and 0.89, for the enucleated cubelets, and 0.76 and 0.83 for the control cubelets; **ED Fig. 9E**). Consistent with the high AUROC scores, VISp cubelets from the enucleated brains also had more neighbors in VISl and VISpm (**Fig. 7H**). Similarly, VISl cubelets from the enucleated brains had more neighbors in auditory areas, and VISpm cubelets from the enucleated brains had more neighbors in VISam and RSPagl (**Fig. 7H**, insets). Strikingly, all three areas shifted towards neighboring areas that were physically further away from VISp, but within the visio-auditory module (black outlines in **Fig. 7H**). To examine whether these changes reflected a shift in area borders or a change in composition across an area, we plotted each cubelet from the enucleated brains and colored them by the differences in the number of neighbor cubelets in VISl (**ED Fig. 9F, *top***) and VISpm (**ED Fig. 9F, *top***). In VISp, the enrichment of neighbors in VISl and VISpm were found in cubelets across the whole area. In particular, cubelets that had more neighbors in VISpm following enucleation (red dots in VISp in **ED Fig. 9F, *top***) appeared to be concentrated at the center of VISp rather than at the borders, suggesting that the changes in the similarity among these areas reflected an overall change in cell type composition, not a shift in area borders. Thus, enucleation shifted the H3 type composition-defined area identities of the visual areas towards neighboring areas within the visio-auditory module.

## Discussion

Using BARseq, we have generated cortex-wide maps of transcriptomic types of excitatory neurons with high transcriptomic and spatial resolution across nine animals. These maps revealed that the spatial patterns of gene expression and the compositions of neuronal types delineate the divisions of cortical areas. This area-based specialization in gene expression further revealed similarities across subsets of cortical areas, which we call modules, that are also highly interconnected. By comparing binocular enucleated animals to control littermates, we further showed that the precise cell type compositional profiles of cortical areas are refined by activity-dependent postnatal development. Together, these results reveal that peripheral inputs play a key role in shaping the distinct cell type compositional profiles of cortical areas, especially within a cortical module.

### BARseq reveals cortex-wide molecular architecture across brains

BARseq was originally developed for mapping long-range projections and associating these projections with gene expression, with high-throughput at single-cell resolution (Chen et al. 2019; Sun et al. 2021). Here we demonstrate that BARseq can also be used as a standalone, high-throughput, cost-effective, and reproducible spatial transcriptomic approach to build a cortex-wide map of transcriptomically defined neuronal types. This map not only elaborates upon the distribution of cortical excitatory neuron types previously revealed by single-cell studies (Tasic et al. 2018; Yao et al. 2021b), but can also serve as an “anchor” to associate other neuronal properties and activity with neuronal types. Our cortex-wide map coincides with the emergence of cortex-wide functional mapping, which are largely driven by advances in large-scale recording techniques such as neuropixel recordings and mesoscale calcium imaging (Sofroniew et al. 2016; Jun et al. 2017; Findling et al. 2023). Associating transcriptomic types and neuronal activity even in single cortical areas has already generated valuable information on how neuronal types contribute to cortical processing (Bugeon et al. 2022; Condylis et al. 2022). Thus, our spatial cell-type map provides a foundational resource for understanding the structural and functional specialization of cortical areas.

The present study is focused on understanding the organization of cell types in the cortex, but the same approach can be flexibly applied to any other brain region. Application to other brain regions will likely require optimizing gene panels for the target regions and/or optimizing cell segmentation. Because BARseq uses combinatorial coding in the GII, the number of genes that can be resolved increases exponentially with the number of sequencing cycles (Sun et al. 2021). Thus, BARseq can potentially interrogate more genes to distinguish more transcriptomic types in other brain regions. When examining large numbers of genes, optical crowding may eventually limit the number of reads that can be resolved in each cell, but computationally demixing overlapping signals (Chen et al. 2021a) and/or using multiple sets of sequencing primers can potentially overcome this limitation. Cell segmentation may also be challenging in brain regions with densely packed cell bodies (e.g., dentate gyrus or piriform cortex), and new approaches, such as Cellpose2 (Pachitariu and Stringer 2022), pci-seq (Qian et al. 2020), and ClusterMap (He et al. 2021) may further improve segmentation accuracy in these regions.

In addition to generating a cell type map of the cortex, our study further leverages the high throughput, low cost, and reproducibility of BARseq to establish causal relationships between developmental perturbations and cortex-wide cell type organization by comparing across multiple animals. In this study, we collected eight hemispheres from enucleated and control littermates in eighteen days of sequencing using a single microscope and spent $2,000 in reagents per brain. Furthermore, data across the eight animals had no apparent evidence of batch effects. This ability to compare molecular architecture across animals at such a large scale would be difficult to achieve using other spatial transcriptomic techniques.

### Fine-grained neuronal types reflect cortical area-based specialization

By leveraging a high-resolution map of gene expression across the cortex, our data reveal the transcriptomically defined neuronal type and gene expression basis of cortical areas at a higher spatial resolution than has previously been possible (Tasic et al. 2018; Yao et al. 2021b). Our results suggest that the cell type compositional profiles of cortical areas reflect their modular organization seen in connectivity studies: cortical areas that are highly interconnected also have similar H3 types (**Fig. 8, *top***). This relationship between transcriptomically defined neuronal types and connectivity, which we summarize as “wire-by-similarity,” is distinct from conventional wiring rules that are commonly observed at individual neuronal type level. In most circuits, similar cell types usually share similar inputs and/or outputs. For example, the connections from cortical neurons in a given layer to another layer are similar across areas (Shepherd et al. 2003; Barbour and Callaway 2008; Weiler et al. 2008; Oviedo et al. 2010; Meng et al. 2017), albeit with small differences; we speculate that these differences in connectivity may be associated with distinct area-specific composition of H3 types. Cortical neurons of the same type, however, are not necessarily highly connected [e.g., *Sst* neurons are sparsely connected with each other (Campagnola et al. 2022)]. Wire-by-similarity is thus not a trivial consequence of cell type-specific connectivity observed at a cortex-wide scale, but it is also *not in conflict* with conventional cell type wiring rules: wire-by-similarity does not describe the connectivity of individual neuronal types but rather reflects how divisions within a large brain region (i.e., areas within the cortex) relate to each other in terms of cell types and connectivity. Future studies using BARseq to map the projections of neuronal types at cellular resolution, from multiple cortical areas, at multiple developmental time points, can help resolve the single-cell basis of the wire-by-similarity organization.

**Figure 8.**
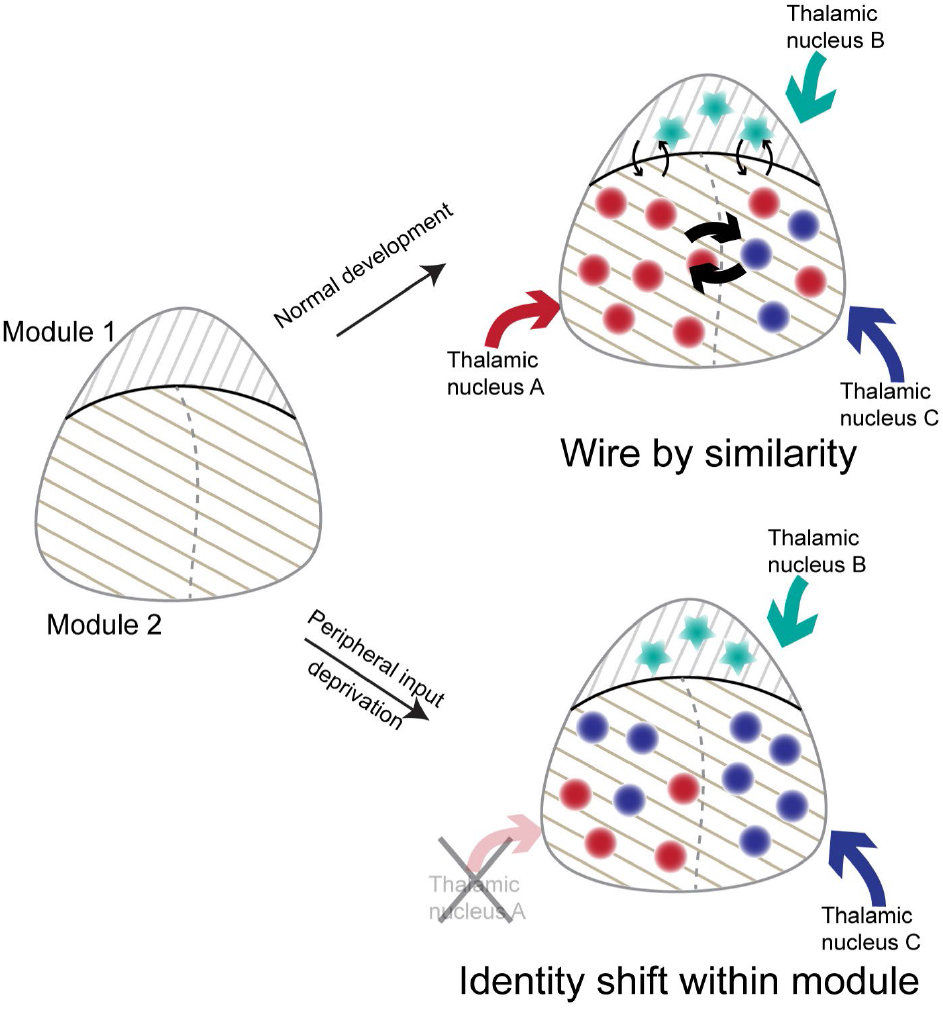
Models of cell type organization across cortical areas and its development. Cortical modules are established independent of peripheral inputs from the thalamus (left), possibly due to earlier developmental mechanisms. Under normal conditions, cortical areas follow the wire-by-similarity rule (top). Areas within a module have similar cell types and are more interconnected. When peripheral inputs are removed (bottom), cell type compositions of the affected area shift towards other areas within the same module. In all three plots, modules are shown by patterned backgrounds. Colored dots and stars indicate cell types in the two cortical modules, respectively. Colored arrows indicate thalamic inputs. Black arrows within the cortex indicate connectivity across areas.

Transcriptomically defined neuronal types are frequently represented hierarchically (Zeng and Sanes 2017), but the biological basis for such hierarchy remains unclear. For example, cortical excitatory neurons and inhibitory neurons are segregated at a coarse granularity in the transcriptomic taxonomy regardless of the cortical area they are from (Tasic et al. 2018; Yao et al. 2021b). In contrast, cortical areas have a strong influence on the activity of both excitatory and inhibitory neurons in response to sensory stimuli. Thus, the hierarchy of transcriptomic taxonomy does not necessarily reflect similarity in stimulus-evoked activity or other functional measures. What, then, generates the hierarchy that is often imposed on transcriptomically defined neuronal types across numerous transcriptomic studies? Our results provide an interesting clue to this question. We found that the medium-grained H2 types represent general cell types that are shared across all cortical areas. In contrast, fine-grained H3 types are restricted in distribution and reflect area specialization. We speculate that different developmental processes established divisions at these two hierarchies: Divisions of H2 types are driven by developmental timing-dependent cell type specification (Shen et al. 2006; Custo Greig et al. 2013), whereas divisions across H3 types are likely driven by mechanisms that contribute to cortical arealization, such as an early diffusive signaling “protomap” and later refinement by thalamocortical axons (Rakic 1988; O’Leary et al. 2007; Cadwell et al. 2019). These developmental processes that give rise to H2 types and H3 types do not arise from a lineage tree, which is necessarily hierarchical. However, because these processes likely affect different sets of genes at different stages of development, the resulting overall gene expression observed in adult neurons differ in magnitude and thus *appear* hierarchical. Consistent with this hypothesis, the observation that spatial gradients in the expression of some genes were shared across H2 types also supports a non-hierarchical origin of the imposed hierarchy of transcriptomically defined neuronal types. Indeed, our experiments showed that post-natal visual deprivation affected the distribution of H3 types across cortical areas, suggesting that the area specialization of H3 type is at least partially caused by thalamic inputs that reflect different sensory modalities. Peripheral input-driven specialization is distinct from earlier developmental timing-dependent specification of H2 types (Shen et al. 2006), and are not inherently hierarchical. Comprehensive interrogation and analysis of gene expression during cortical development combined with lineage tracing will provide crucial evidence to support or refute our hypothesis.

### Peripheral inputs sharpen cell type compositional profiles within a cortical module

Thalamocortical axons play an important role in defining and refining cortical areas and their characteristic cellular and molecular features (Chou et al. 2013; Li et al. 2013; Pouchelon et al. 2014; Cadwell et al. 2019; Cheng et al. 2022). In our experiments, the combination of single cell resolution, high transcriptomic resolution, and broad interrogation across many cortical areas allowed us to reveal unprecedented details of how gene expression and cell type compositional profiles change after enucleation. Overall, the effects of enucleation on the visual areas suggest that peripheral activity refines the cell type compositional profiles of cortical areas, so that they acquire the compositional profiles that are characteristic to each area. Interestingly, many H3 types that were enriched in VISp following enucleation were often found in the control brains in lateral and medial areas, such as the anterior cingulate areas and the agranular insular areas, which are far outside of the visio-auditory module. This broad distribution of enriched H3 type is expected from our results in the pilot dataset: Whereas most individual H3 types are broadly distributed across many cortical areas, the compositional profiles of H3 types are distinct across cortical areas, and thus reflect area identity. This distinction between how area identities and neuronal identities change highlights the utility of comprehensively interrogating cortex-wide neuronal gene expression following developmental perturbation.

The changes associated with binocular enucleation in our experiments are different from, but consistent with changes in gene expression observed in previous studies that performed related perturbations (see **Supplementary Note 7** for details) (Chou et al. 2013; Cheng et al. 2022). Enucleation affected IT neurons in all layers and L6b/CT neurons, but dark rearing during the critical period only affected IT neurons in layer 2/3 (Cheng et al. 2022), suggesting that the deep layer neurons are sensitive to peripheral sensory deprivation only during earlier developmental time points. This sensitivity for sensory deprivation during early postnatal development is consistent with the early innervation of deep subplate/future L6b neurons by thalamic afferents and their susceptibility to early sensory manipulations (Friauf and Shatz 1991; Molnar et al. 2020; Meng et al. 2021). Because enucleation also affected L6b/CT neurons, which are related to the subplate/L6b neurons but usually enriched in lateral areas, our results are consistent with a role of thalamic innervation of the subplate/L6b neurons in determining and refining areal identity. The cell type changes we observed, however, did not completely abolish the distinction between primary and secondary visual areas, as observed by Chou et al. (2013) after genetically ablating the thalamocortical axons. Genetic ablation of thalamic projections postnatally also has strong effects on L4 neuron identity and function, suggesting an ongoing role for thalamocortical projections in maintaining area specificity (Pouchelon et al. 2014). Thus, our results, together with previous studies, suggest a consistent model: the physical connections established by thalamocortical axons are needed to define the primary visual cortex, and the peripheral activity sharpens the cell type composition across both the primary visual cortex and neighboring higher visual areas (**Fig. 8, *bottom***). Our study does not provide information on the mechanism through which peripheral activity refines cell type composition: These changes could be driven by thalamic inputs regardless of activity pattern, by patterned activity that is characteristic of visual inputs, by inducing cell death in select cell types, and/or by rewiring of thalamocortical axons and/or cortical connectivity due to lack of activity. Distinguishing these possibilities and determining whether the refinement role of peripheral inputs is restricted to within modules for other sensory modalities will require future experiments exploring changes in projections with finer manipulation of thalamic and cortical activity.

### BARseq enables brain-wide comparison of cellular gene expression across individuals

Animals in a natural population, like human populations, are diverse in their genetics and anatomy; even mice within the same inbred strain can display differences in brain morphology and development (Valiquette et al. 2023). Thus, ideally, a reference transcriptomic atlas of the brain should encompass thousands of individual animals to generate consensus and to estimate individual variability. This approach mirrors progress made in human genomics in the past two decades, starting from the first human genome draft (Lander et al. 2001; Venter et al. 2001) to recent drafts of the human pangenome (Gao et al. 2023; Liao et al. 2023). Similar thoughts also guided the creation of the current version of the Allen Common Coordinate Framework (Wang et al. 2020), which involved averaging fluorescent images across thousands of animals. Such diversity, however, is largely obscured in current mouse brain atlasing efforts (Langlieb et al. 2023; Yao et al. 2023; Zhang et al. 2023), which applied spatial transcriptomic approaches to one to three animals. The number of animals that can be interrogated is largely constrained by the low throughput and high cost of many spatial transcriptomic approaches. Furthermore, technical variability across samples (i.e., batch effects) also confounds integrating data across multiple animals, making it difficult to interpret any differences found across animals. These factors raise questions of how generalizable these atlases are across individual animals and across strains of mice, and limit how they can be used to help interpret other experiments. BARseq overcomes all three barriers (throughput, cost, batch effect) to enable brain-wide interrogation across many animals. Because it is easier to associate neuronal morphology, electrophysiology, connectivity, and neuronal activity with gene expression than with cytoarchitecture, redefining a mouse brain common coordinate framework using transcriptomic type information could be more informative than the existing one based on cytoarchitecture. Our current throughput (two days per hemi-brain) already allows hundreds of animals to be interrogated in a microscope year, and moderate scaling up can easily achieve thousands of animals per year. BARseq thus provides a path to go beyond a reference mouse brain transcriptomic atlas based on representative individuals (Langlieb et al. 2023; Yao et al. 2023; Zhang et al. 2023) towards a “pan-transcriptomic” atlas that builds consensus across individuals and captures population diversity.

In addition to creating a pan-transcriptomic atlas of the mouse brain, the ability to interrogate and compare brain-wide molecular architecture across multiple individuals can facilitate discovery broadly across the neuroscience field. This study provides a proof-of-principle by identifying how cell types change across the whole cortex in response to a developmental perturbation. The same approach can also be used to generate information on cell type differences associated with neuropsychiatric disease models, age, species, and other experimental perturbations. The ability to identify differences brain-wide is especially useful to understanding neuropsychiatric conditions, because many neuropsychiatric conditions and/or vulnerability to mental disorders are associated with differences across many brain regions. Our approach based on BARseq can thus be generalized to other brain regions and questions to bridge brain-wide network-level dynamics with detailed changes in gene expression in single neurons and to establish causal relationship between developmental processes and brain-wide cell type organization.

## Methods

### Animals and tissue processing

All animal and surgical procedures were carried out in accordance with the Institutional Animal Care and Use Committee at Cold Spring Harbor Laboratory and Johns Hopkins University. The animals were housed at maximum of five in a cage on a 12-h on/12-h off light cycle. The temperature in the facility was kept at 22 °C with a range not exceeding 20.5 °C to 26 °C. Humidity was maintained at around 45–55%, not exceeding a range of 30–70%. The animal used to generate the pilot dataset was 7–8-week-old C57BL/6J animals purchased from Jackson Laboratory.

Bilateral enucleation surgeries were performed on C57BL/6J (JAX strain no. 000664) mouse pups of both sexes on postnatal day (P) 1 using previously published methods (Dye et al. 2012; Deng et al. 2021; Mukherjee et al. 2023). Pups were anesthetized briefly with 1-2% inhaled isoflurane (Fluriso, VetOne, Boise, ID). The eyelids were opened with a sterile surgical scalpel blade. Using fine forceps, eyeballs were lifted away from the orbit and freed from the optic nerve and surrounding musculature. Following eye removal, eyelids were closed and sealed with surgical glue (Vetbond, 3M, Maplewood, MN). After surgery, pups were placed in a plastic box in a warm water bath maintained at 37°C for ∼1 h for recovery before returning to their mothers. Only one-half of the entire litter underwent enucleation surgery to ensure maternal care and the thriving of the pups. Sham surgery was performed as the control on the other half of the litter at the same age, where pups were subjected to anesthesia and revival procedures as described above. All pups were housed with their mothers until they were used for experiments at P28 and were weighed routinely to ensure normal thriving.

To collect brains for BARseq, we euthanized the animals with isoflurane overdose, decapitated the animals, embedded the brain in OCT and snap-froze in an ethanol dry-ice bath. In experiments in which only one hemisphere was used, we bisected the brain along the midline before OCT embedding. The brains were then sliced to 20 μm hemi-coronal sections and mounted onto Superfrost Plus Gold slides. Eight sections were mounted onto each slide.

### BARseq library preparation

BARseq samples were prepared as previously described (Sun et al. 2021). Briefly, slides were fixed in PFA, dehydrated in ethanol, rehydrated in PBST (PBS with 0.5% Tween-20), then reverse transcribed using random primers with a N-terminal amine group and Revert-aid H-minus reverse transcriptase (Thermo Fisher). On the second day, we crosslinked the cDNA products, then performed padlock probe hybridization and ligation, followed by rolling circle amplification with ϕ29 DNA polymerase (Thermo Fisher). On the third day, we crosslinked rolonies. To sequence the samples, we hybridized sequencing primers and manually performed sequencing using either the Illumina HiSeq Rapid SBS kit v2 (for the pilot dataset) or the Illumina Miseq nano v2 kit (for the eight littermates) following previous protocols (Sun et al. 2021). After seven sequencing cycles were completed, we then striped the sequencing primers using three incubations at 60℃ for 5 mins each in 60% formamide, 2× SSC, 0.1% Tween 20. We then hybridized fluorescent probes in 10% formamide, 2× SSC for 10 mins, followed by incubation in 2 µg/mL DAPI in PBST for 5 mins. We then imaged the final hybridization cycle. Detailed protocols are available at protocols.io: dx.doi.org/10.17504/protocols.io.81wgbp4j3vpk/v2. Primers and padlock probes used are listed in **Supplementary Table 1**.

### BARseq data collection

BARseq data from the pilot brain was collected as described previously (Sun et al. 2021) on a Olympus IX-81 microscope with PI P-736 piezo z-stage, Olympus UCPLFLN20X 20× 0.7NA air objective, a Crest xlight v2 spinning disk confocal, 89North LDI-7 laser bank, and Photometrics BSI-prime camera. Because the microscope stage could only fit 3 slides at a time, the posterior 22 sections and the anterior 18 sections were collected in two separate batches. BARseq data from the eight littermate brains were collected on a Nikon Ti-2E microscope with PI P-736 piezo z-stage, Nikon CFI S Plan Fluor LWD 20XC 0.7NA objective, a Crest xlight v3 spinning disk confocal, Lumencor Celesta 7-line laser, and a Photometrics Kinetix camera. Four slides with eight slices per slide, all from the same animal, were imaged in each batch. The filters and lasers used are listed in **Supplementary Table 2**.

### BARseq data processing

BARseq data were processed as described previously(Sun et al. 2021) with slight modifications. Briefly, we created max-projections from the image stacks, applied noise reduction using Noise2Void (Krull et al. 2018), applied background subtraction, corrected for channel shift and bleed-through, and registered images through all sequencing cycles. We segmented cells using Cellpose (Stringer et al. 2020), decoded rolonies using BarDensr (Chen et al. 2021a), and assigned rolonies to cells. During BarDensr decoding, we used negative control GIIs (i.e., GIIs that were not carried by any padlock probe) to estimate false discovery rate (FDR), and automatically adjusted decoding threshold to target about 5% FDR. Finally, the images were stitched to generate whole-slice images. All processing steps were performed on each imaging field of view (FOV) separately, not on the stitched images, to avoid stitching errors, to minimize alignment artifacts due to imperfect optics, and to facilitate parallel processing. The stitched images were only used to generate a transformation, which we applied to each rolony and cell after all other steps were finished. Overlapping cells from neighboring FOVs were detected using a custom implementation of sort and sweep (Baraff and Witkin 1992), an algorithm that is commonly used in detecting object collision in video games. When two or more cells were detected as overlapping, cells that had more read counts (which we assumed to have higher read quality) were kept, and all other cells were removed. See **Data and Code Availability** for processing script and intermediate processed data.

We then registered each stitched slice to Allen CCFv3. For each slice, we first generated an image that color coded each cell by its H2 type identity. We then used QuickNii (v.3 2017) to manually select the CCF plane for each slice and Visualign (v. 0.9) to align area borders within each slice. After we obtained CCF coordinates for each neuron, we registered all cortical cells onto a CCF flat map using python ccf_streamlines package (https://pypi.org/project/ccf-streamlines/). Streamlines represent the paths that most directly connect the pia of the isocortex to the white matter while following the curvature of those surfaces. Because the streamlines were defined in CCF space and are thus consistent across all animals, this approach allowed us to more reliably obtain relative cortical depth and relative cortical location of each neuron across the eight brains compared to the manual approach used in the pilot brain.

### Quality control

After segmentation, we obtained a total of 13,886,988 segmented cells. We kept all cells expressing at least 5 unique genes and at least 20 total counts, resulting in a count matrix with 104 genes and 10,378,092 cells.

### Iterative clustering and annotation of BARseq transcriptomic data

To obtain H1, H2, and H3 types in the pilot brain, we adopted an iterative clustering pipeline adapted from single-cell RNA sequencing (scRNAseq) studies (Amezquita et al. 2020). We performed 3 rounds of clustering with the following steps: normalization, dimension reduction (PCA), computation of a shared nearest neighbor (SNN) network, Louvain clustering. We normalized counts to CP10 (counts per 10) values, then applied the log1p transformation (log with a pseudo-count of 1). Because our panel consists of marker genes, we skipped highly variable gene selection and ran PCA on all genes. We computed PCA using the scater::runPCA function (McCarthy et al. 2017) and kept the first 30 PCs. We built the SNN network using the scran::buildSNNGraph function with 15 nearest neighbors and the “rank” metric. We ran the Louvain clustering as implemented in the igraph::cluster_louvain function with default parameters. All UMAP visualizations were obtained using the scater::runUMAP function starting from the PCA dimension reduction and using the 15 nearest neighbors.

In the first round of clustering, we separated cells in three classes reflecting neurotransmitter expression: excitatory (expressing *Slc17a7*), inhibitory (expressing *Gad1*), and others (expressing neither *Slc17a7* nor *Gad1*). Because marker genes are frequently undetected at the single-cell level, we ran the clustering pipeline on all cells, obtaining 24 clusters, then assessed marker expression at the cluster level. From the UMAP visualization, we distinguished 3 groups of clusters. The first group contained excitatory clusters expressing *Slc17a7*, the second group contained inhibitory clusters expressing *Gad1*, the third group expressed neither of these markers. Based on these observations, we manually annotated the clusters as “excitatory”, “inhibitory”, and “other”, respectively (H1 types, 642,340, 427,939, and 188,977 cells, respectively). In the second round of clustering, we extracted all cells labeled as “excitatory” and “inhibitory”, then ran our pipeline again on each H1 type separately. By re-running the pipeline, the dimension reduction and clustering are better targeted at finding variability specific to either excitatory or inhibitory cells. We obtained 18 excitatory clusters and 11 inhibitory clusters, which formed the basis for our H2 types. In the third round of clustering, we re-ran the pipeline on each excitatory H2 type, obtaining roughly 5 to 6 clusters by type (117 total H3 types).

To annotate H2 and H3 types in the pilot brain, we examined the brain-wide distribution and the marker expression of H2 and H3 types. Almost all H3 types showed brain-area specificity, suggesting that the data were clustered at a biologically meaningful granularity. We annotated isocortical H2 types based on aggregate marker expression (see **Supplementary Note 1** for marker selection). We annotated non-isocortical H2 types based on the localization of cells (e.g., hippocampal areas, thalamus, entorhinal cortex). When H2 types contained a mix of isocortical and other cells (e.g., “L6 IT-like”), we split the H2 type into multiple H2 types, one containing the isocortical cells (e.g., “L6 IT”), the others containing the other cells (e.g., “PIR L6 IT-like”, “AON DL”). In the end, we obtained a list of 11 cortical H2 types and 23 non-cortical H2 types. After mapping to scRNAseq reference types, we noticed that four H3 types (“PT AUD”, “PT P RSP/IT-like?”, “RSP DL”, “CT CTX A/L-V”) were assigned to incorrect H2 types (“L5 IT”, “RSP DL”, “L5 IT”, “L6b”); we manually corrected their H2 annotation (to “PT”, “PT”, “RSP DL”, “CT”).

The eight littermate brains were processed as a single batch using the same overall procedure as the pilot brain, with the following adjustments at the H1 and H2 levels to minimize computational load. We used scater’s calculatePCA and calculateUMAP functions (with external_neighbors=TRUE and BNPARAM=AnnoyParam()). We computed clusters using Seurat’s FindNeighbors function(Stuart et al. 2019) followed by Leiden clustering(Traag et al. 2019) using igraph::cluster_leiden (with objective_function=”modularity”, resolution=0.2 at the H1 level and 0.8 at the H2 level). For cell type annotation, we used a combination of the strategy described above and top hits to pilot brain annotations using MetaNeighbor.

### Mapping of BARseq types to scRNAseq reference types

To map BARseq types to reference types, we used a k-nearest neighbor (kNN) approach to label each cell according to its closest neighbors in a whole-cortex and hippocampus reference compendium (Yao et al. 2021b). First, we evaluated the accuracy of the kNN approach on a subset of the reference compendium using leave-one-cell-out cross-validation (10X MOp dataset, “Glutamatergic” cells, cortical cells labeled as “CTX” or “Car3”, clusters with ≥ 50 cells, CP10K and log1p normalization). To transfer labels, we picked each cell’s closest 15 neighbors using the BiocNeighbors::queryKNN function, then predicted the cell’s type by taking a majority vote across the neighbors. We compared accuracies across 4 gene panels: highly variable genes (HVGs, 2000 genes selected using scran::modelGeneVar and scran::getTopHVGs), HVG selection followed by PCA (30 components, scater::runPCA), the BARseq panel (104 genes, after excluding *Gad1* and *Slc17a7*), the BARseq panel with reads down-sampled to match BARseq sensitivity (104 genes, binomial sampling of reads, re-normalization through sample-wise ranking and scaling). For the latter gene set, reads were downsampled for each gene according to the sensitivity ratio (BARseq average counts divided by reference average counts). For genes that had a sensitivity ratio *r* > 1, we oversampled reads to match BARseq sensitivity (reads multiplied by ⌊*r*⌋ + binomial sampling with probability ⌊*r*⌋ − *r*).

Having validated that the kNN mapping procedure was able to assign cell types with high accuracy, we applied the same procedure (sensitivity adjustment of reference datasets, sample-wise ranking and scaling of reference and target datasets, BiocNeighbors::queryKNN with 15 neighbors) to assign a reference label to each BARseq cell. In contrast with the previous evaluation, we used all excitatory cells from the reference compendium (40 individual datasets, all “Glutamatergic” cells) and adjusted reference reads using a simplified downsampling procedure (reads multiplied by the sensitivity ratio for each gene). To compute the overlap between BARseq and reference cell types, we used the Jaccard coefficient (number of cells labeled as BARseq type 𝑏𝑏 and predicted to be reference type *r* divided by the number of cells of type 𝑏𝑏 or predicted to be type *r*).

We mapped both H3 types in the cortex (**Fig. 2E**) and those in the hippocampal formation (**ED Fig. 2F**) to single-cell RNAseq data. However, we do not expect perfect matching for clusters outside of the cortex, because our dataset sampled additional brain regions that were not sampled in the single-cell RNAseq data. In addition, our gene panel was optimized for cortical excitatory neurons and could miss highly differentially expressed genes in other brain regions.

We mapped H3 types from the eight littermate brains against the pilot brain and scRNAseq data from (Cheng et al. 2022) using the procedure outlined above.

### Variance of expression explained by H2 types, H3 types, and space

To evaluate how well H2 types, H3 types, and spatial information recapitulate the variability of expression, we performed one-way ANOVA on pseudo-bulk data. This analysis was run on a subset of data containing the 8 isocortical H2 types with isocortex-wide localization (“L2/3 IT”, “L4/5 IT”, “L5 IT”, “L6 IT”, “PT”, “NP”, “CT”, “L6b”).

We started by computing the pseudo-bulk expression matrix *B_gts_*, providing expression of gene g for H3 type t in spatial bin s. We defined 540 spatial bins containing an average of 14 cells per H3 type as follows: 27 slices along the A-P axis (corresponding to slices 7 to 33), 20 bins along the un-warped M-L axis for each slice (chosen to balance the number of cells in each bin, computed independently for each slice). Slices at the anterior and posterior end of the brain were excluded because coronal sections were not perpendicular to the cortical surface at these extreme positions and would thus bias gene expression. Starting from the gene by cell count matrix *C_gc_*, we have *B_gts_* = *mean_cϵt,cϵs_* (*C_gc_*).

Next, we computed the variance explained by the 8 H2 types, 51 H3 types and 540 spatial bins by applying the one-way ANOVA formula for each gene and factor independently. Let *M* = *mean_tϵtH3,sϵbin_* (*B_gts_*) be the overall average expression and *T* = Σ*_tϵH3,sϵbin_* (*B_gts_* − *M*)^2^ be the total variance. For gene g, we computed the variability explained as follows:

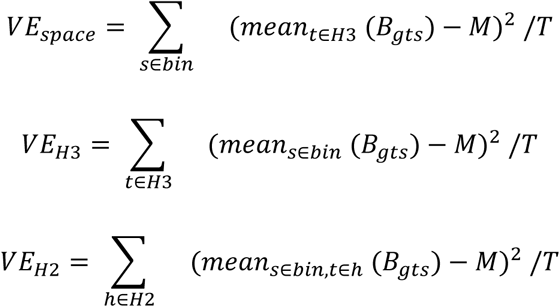

Because H3 types are nested factors of H2 types, the variability explained by H3 types is necessarily higher; the additional variability explained by H3 types is given by 𝛥𝑉𝐸 = 𝑉𝐸_𝐻3_ − 𝑉𝐸_𝐻2_.

### Extraction of recurrent spatial patterns using non-negative matrix factorization

The ANOVA analysis revealed the presence of recurrent spatial patterning across genes and H2 types. We used non-negative matrix factorization to extract these patterns. First, we defined a pseudo-bulk matrix using the same procedure as the ANOVA analysis (see above), except that we computed the pseudo-bulk matrix at the level of H2 types. Starting from the count matrix 𝐶_𝑔c_, we have *B_gts_* = *mean_cϵt,cϵs_* (*C_gc_*), where 𝑡𝑡 is one of the 8 H2 types. Here, we consider the spatial bins as features (rows), genes and types as variables (columns), resulting in a matrix with 540 rows and 848 columns. Because spatial patterns had different scales (average level of expression) across genes and H2 types, we rescaled each column using L2-normalization (squared columns sum to 1). This rescaling ensures that factors reflect recurrent patterns (seen in multiple genes and H2 types) rather than a single instance (highly expressing gene in one H2 type). This procedure (pseudo-bulking at H2 level and rescaling) can also be seen as a correction for H2 type composition (overall expression patterns are dominated by the most common or the highest expressing H2 type). We extracted 10 NMF factors using the NMF::nmf function using default parameters (Brunet algorithm), obtaining a nonnegative basis matrix W (540 bins by 10 factors) containing spatial patterns and a nonnegative coefficient matrix H (10 factors by 848 columns) such that B≈W.H. For later analysis, we removed 3 NMF factors that reflected obvious slice-specific batch effects (see **Supplementary Note 4**).

To identify genes associated with each NMF factor, we computed the average fraction of expression variance explained by each NMF factor. Given an NMF factor f, we computed the predicted gene expression for gene g in type t as 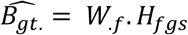, where 𝑊_.𝑓_ is the column in W representing factor f, and *H_fgs_* is the coefficient associated with factor f, gene g and type t. The variance explained is then computed as 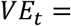, where 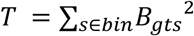 is the total uncentered variance. We then took the average variance explained across the 8 H2 types. To estimate the null fraction of variance explained, we permuted coefficients associated with each factor (permutation of rows in H), then recomputed the average variance explained across all genes and factors. We labeled gene-NMF associations as “significant” if the average variance explained exceeded the 99th percentile of the null distribution.

To identify H3 types associated with each NMF factor, we computed two measures of association: enrichment of NMF-associated genes and correlation of spatial distribution with NMF patterns. Using the MetaMarkers package, we computed differentially expression (DE) statistics for each H3 type (1-vs-all differential expression against other H3 types from the same H2 type). We then asked whether top DE genes were enriched for NMF-associated genes (genes with significant association with a single NMF). We report the enrichment as an AUROC, asking how well expression fold changes predict NMF-associated genes. Independently, we computed the overlap between NMF patterns (columns in W) with the spatial distribution of H3 types (count of cells in bin s divided by total count) using the Spearman correlation. Because the dynamic range of the correlation coefficient depends on the sparsity of the pattern (sparser patterns tend to have a smaller range of correlation values), we report the association as the Z-scored Spearman correlation (Z-scoring across H3 types within a given H2 type and factor).

### Predicting cortical areas and locations using gene expression and H3 type composition

We first binned cells in the cortex into cubelets, which were drawn separately on each coronal section, spanned all cortical layers, and were about 100 μm - 200 μm on the M-L axis along the curvature of the cortex. To draw the cubelets so that their medial and lateral borders were perpendicular to the layers, we manually drew matching points along the top and bottom surface of the cortex, especially at locations where the curvature of the cortex is extreme (e.g., the medial part of the cortex). This step thus separates both the bottom and top surfaces of the cortex into several segments. Within each segment, we then cut the two surfaces into the same number of smaller segments of roughly equal distance. Each cubelet was then defined by connecting the ends of the small segments. Slices at the anterior and posterior end of the brain were excluded because coronal sections were not perpendicular to the cortical surface at these extreme positions and would thus bias composition of neuronal populations. Unlike the spatial bins used in the NMF analysis, which had equal cell numbers within each slice but unequal widths, the cubelets had similar widths across all slices, but not necessarily similar cell numbers (**ED Fig. 4A**).

Because the cubelets are generated within each of the large segments, they may be slightly different in size across different segments. We thus normalized both H3 type counts and gene read counts within each cubelet by the total number of cells and/or gene reads for downstream analyses. For the A-P locations of each cubelet, we used the CCF coordinates directly. For the M-L locations, we calculated an un-warped coordinate along the cortex within each coronal section as follows. We connected the centroids of adjacent cubelets and defined the un-warped distance between two cubelets as the sum of all connected lines across all cubelets between the two. We then defined the zero position along this un-warped M-L axis as the point that was closest to the point in CCF where the midline of the brain intersects the top surface of the brain. Thus, cubelets on the medial side have negative M-L coordinates, whereas those on the dorsolateral side have positive M-L coordinates.

To predict the coordinates of each cubelet, we trained random forest regression models, each with 50 trees and an in-bag-fraction of 0.5. We used 100-fold cross-validation to evaluate the performance of the models. To evaluate the prediction performance using the composition of H3 types within each H2 type, we trained similar regression models using only the relevant H3 types and evaluated performance using 5-fold cross-validation. To predict cortical area labels in CCF, we first assigned CCF area labels to each cubelet using its centroid location. We then trained random forest classifiers with 500 trees and an in-bag-fraction of 0.5 and evaluated the performance with 10-fold cross validation. All models were built in MATLAB using the TreeBagger function.

### Correspondence between abrupt changes in the composition of H3 types and area borders defined by CCF

To find positions of abrupt changes in the composition of H3 types within each coronal section, we performed principal component analysis on the composition of H3 types and calculated the two-norm of the 1st derivative of the first five principal components along the un-warped M-L axis. We then convoluted the two-norm of the derivatives with a smoothing window of the shape [0.25, 0.5, 0.25]. We then looked for local peaks with prominence that was larger than half a standard deviation, and with the value at the peak higher than the median of the smoothed norm of the derivatives. Peaks were considered close to a CCF-defined border if it was within 150 μm from that border, calculated based on the CCF coordinates of the centroids of cubelets. All relevant analyses were performed in MATLAB.

### Inferring cortical modules from H3 type composition

We only clustered areas with at least 7 cubelets. For each pair of areas, we built a support vector machine classifier with Lasso regularization to predict the area identity given H3 type composition. We calculated the area under the ROC curves following 5-fold cross validation. The AUROC values were used to build a distance matrix among cortical areas. We then performed Louvain community detection using this distance matrix, which generated six clusters and three areas that did not cluster with any other areas. We then built a dendrogram of the six clusters and three areas using the medians of all pairwise distances between areas from each pair of clusters and ward linkage. We then manually cut the dendrogram to generate the five modules. The cortical flatmap of modules was drawn based on that in (Harris et al. 2019), and color-coded manually based on this analysis.

To test how the strength of cortical modularity, we calculated the modularity of clusters with the following perturbations: (1) Randomly shuffle cortical area labels across all cubelets. (2) Randomly assign each cubelet with an area label within 1-cubelet distance with same probabilities. That is, for each cubelet, there is an equal chance of assigning an area label of the cubelet itself, or the labels of the two cubelets adjacent to it. (3) Randomly assign each cubelet with an area label within 2-cubelet distance with the same probabilities. In addition, for the random shuffling control, we calculated modularity for both the clusters identified by Louvain clustering on the original data and the clusters identified by Louvain clustering on the shuffled data.

All relevant analyses were performed in MATLAB. The linear classifier was generated using the fitclinear function, and Louvain community detection was performed using a MATLAB implementation by Antoine Scherrer.

### Gene expression changes in the enucleated animals

To assess the nature of the expression shift from control to enucleated animals, we first investigated the neighborhood of individual neurons. If types appeared or disappeared in enucleated animals, we would expect that the neighborhood of groups of neurons would be entirely composed of neurons of either control or enucleated animals. To avoid donor and litter effects, we only considered neighborhoods across litters (e.g., neurons from donors 3 to 8 for donor 1). For each neuron, we picked the k = 100 closest neighbors (using the same procedure as kNN dataset mapping, sample-wise ranking and scaling followed by queryKNN), then computed the fraction of neurons from control animals.

We then used the same procedure to better understand the nature of the shift. Instead of looking at the “condition” label of each neighbor (control or enucleated), we collected the brain area labels of the top k = 20 neighbors. We aggregated these counts at the H2 level for each animal, then performed a two-sided hypergeometric test within each litter to identify brain region neighborhoods that were significantly depleted and enriched in enucleated animals. We combined p-values from individual litters into an overall p-value using Fisher’s method, then corrected p-values using the FDR procedure.

### Cell type composition changes in the enucleated animals

To assess how cell type composition is related to cortical areas in control and enucleated animals, we analyzed the differences in H3 type composition for each cubelet in all control and enucleated littermate brains. For the eight littermate brains, we used the flatmap mapping to define cubelets. For each slice, we fitted a polynomial curve line using flatmap coordinates of the middle layer neurons (relative depth between 80 to 120 on a scale of 0 to 200) and used this fit to represent the midline of the cortex. We then projected neurons at all depths onto this midline based on their flatmap coordinates. After projecting all neurons, we segmented the fitted line into 50 roughly equal segments. In this way, we assigned neurons into cubelets with consistent number of neurons for the control and enucleated littermate brains.

For all cubelet-based analyses, we filtered out cubelets with damaged or warped tissue, as well as cubelets on the posterior edge of cortex that did not contain all cortical layers. Areas were assigned to each cubelet based the area assigned to the majority of neurons within that cubelet. UMAP plots were generated using the fraction of H3 types in each cubelet.

To test how distinguishable an area is between the enucleated and the control brains, for each cubelet we identified the 10 nearest neighbor cubelets from the same cortical area in the six animals that came from different litters. We randomly selected 20% of all available cubelets for each iteration and performed 50 iterations. For shuffled controls, we shuffled the conditions of the six brains, but cubelets within each brain retained the same condition label as other cubelets from the same brain. We then calculated AUROC based on the condition labels for the nearest neighbors. We performed t-test and used the Benjamini-Hochberg procedure to find the false discovery rate (FDR).

To test how well each area from the enucleated or the control brains maps onto areas in the control brains, for each cubelet we identified the 20 nearest neighbor cubelets from any cortical area in the three control animals that came from different litters. Then for each pair of cortical areas A and B, we then calculated AUROC for classifying cubelets from A or non-A areas in the test brain to B or non-B areas in the control brains. To test the shift of cell types within each H2 type, we performed the same test using only the compositions of H3 types that belonged to a H2 type.

### Data and code availability

Raw sequencing images are available from the Brain Image Library. Cell-level and rolony-level data and scripts used for both data processing and data analysis are provided at Mendeley data (doi: 10.17632/8bhhk7c5n9.1 and 10.17632/5xfzcb4kn8.1) (https://data.mendeley.com/datasets/8bhhk7c5n9/draft?a=f37296fe-dc00-46a6-9e69-46f4c3d0deec and https://data.mendeley.com/datasets/5xfzcb4kn8/draft?a=45f018da-d0cf-4bde-b7a4-e3a567e9edaf for preview) and on Github (https://github.com/gillislab/barseq_analysis).

## Acknowledgement

The authors thank W. Wadolowski for technical support; A. Bhandiwad for support in converting CCF to flatmap coordinates; K. Brouner for histology support; B. Tasic, Z. Yao, B. Long, D-W. Kim, and other members of the Allen Institute for discussion. This work was supported by the National Institutes of Health (5RO1NS073129, 5RO1DA036913, RF1MH114132 and U01MH109113 to A.M.Z; R01MH113005 and R01LM012736 to J.G.; U19MH114821 to A.M.Z. and J.G.; 1DP2MH132940 to X.C.; R01DC009607 to P.O.K), the Brain Research Foundation (BRF-SIA-2014-03 to A.M.Z.), IARPA MICrONS (D16PC0008 to A.M.Z.), and Robert Lourie award (to A.M.Z.). A.M.Z. was supported by an Allen Distinguished Investigator Award, a Paul G. Allen Frontiers Group advised grant of the Paul G. Allen Family Foundation. The content is solely the responsibility of the authors and does not necessarily represent the official views of the National Institutes of Health. X.C., M.R., and A.Z. wish to thank the Allen Institute founder, Paul G. Allen, for his vision, encouragement, and support.

## Author contribution

X.C. conceived the study, S.F. selected the gene panel, D.M. and P.O.K. designed and performed binocular enucleation, X.C., M.R., A.Z. collected data, X.C., S.F., M.R., J.G., A.M.Z., and A.Z. analyzed the data, X.C., M.R., S.F., A.M.Z. drafted the manuscript, all authors were involved in the interpretation of results and finalizing the manuscript.

## Materials and correspondence

Correspondence and requests for materials should be addressed to X.C.

---

**ED Figure 1.**
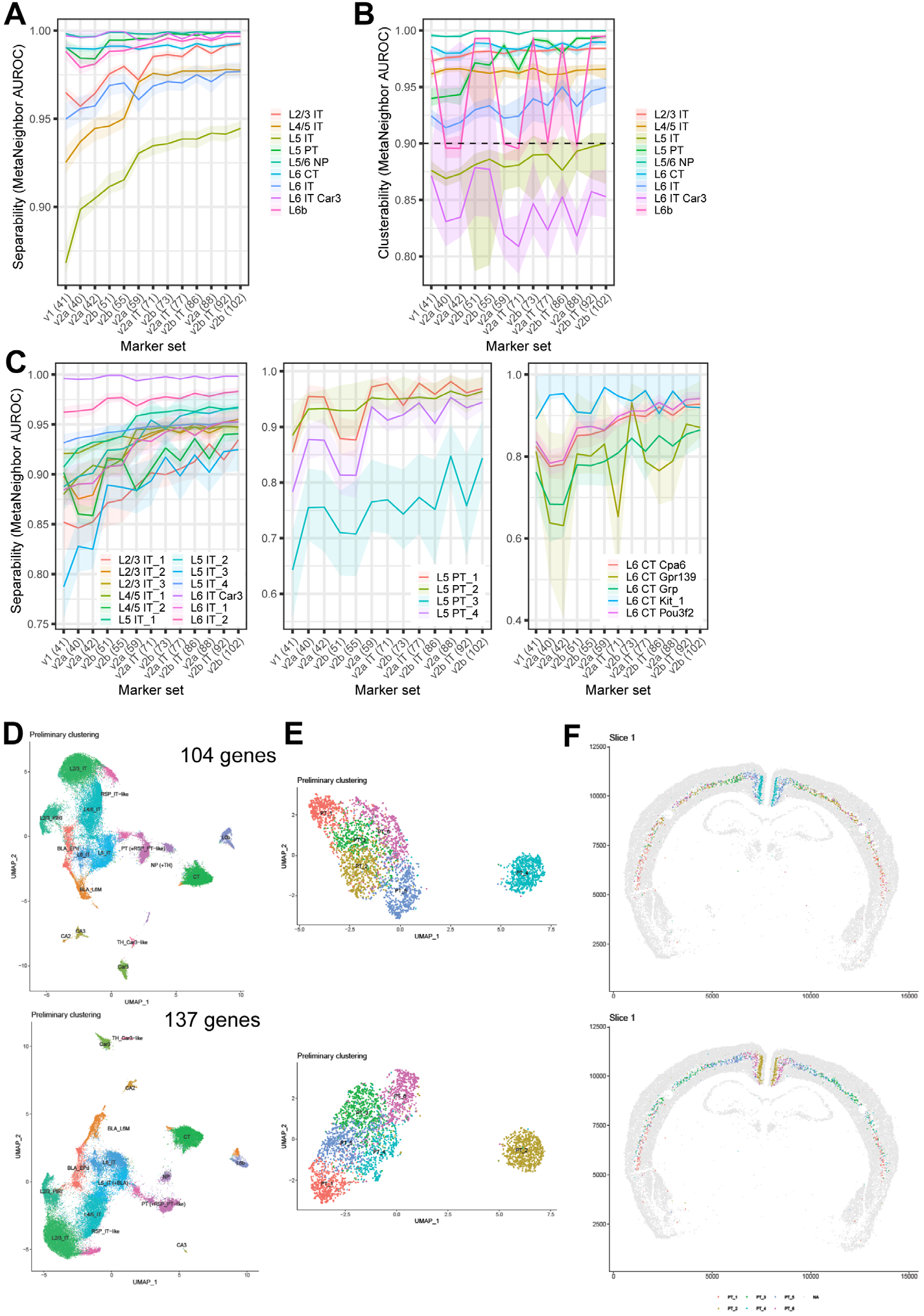
In silico design and evaluation of the gene panel. (A) *In silico* assessment of H2 type separability (supervised analysis, ability to distinguish cell types given reference labels) in a surrogate snRNAseq dataset across marker sets ranging from 40 to 102 genes. The number of genes in each panel is shown in parentheses. The final gene panel include the 102-gene panel, plus two manually added genes (*Fezf2* and *Hgf*). **(B)** In silico assessment of H2 type clusterability (unsupervised analysis, ability to recover reference labels through standard clustering) across the marker sets. **(C)** In silico assessment of H3 type separability for IT, L5 ET (PT), and L6 CT cells across the marker sets. **(D)(E)** UMAP plots of gene expression of cortical excitatory neurons (D) and L5 ET neurons (E) calculated from the 104-gene panel with or without an additional 33 genes. Colors indicate H2 types in (D) and H3 types in (E). **(F)** Images of a coronal section showing the distribution of L5 ET types clustered using the gene panels.

**ED Figure 2.**
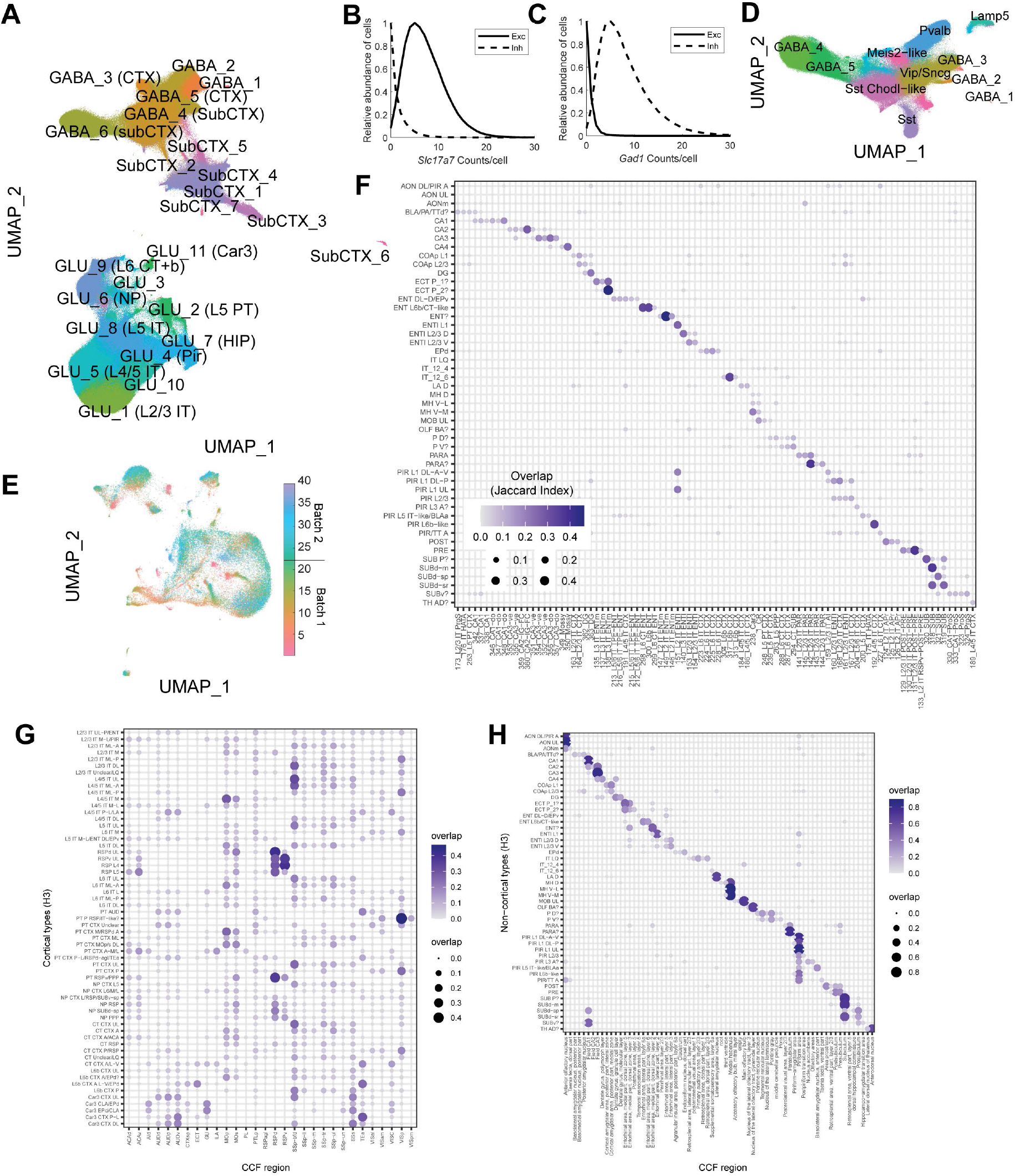
Hierarchical clustering of BARseq data. (**A**) UMAP plot of the gene expression of all cells. Colors and labels indicate H1 clusters. (**B**)(**C**) Histograms of *Slc17a7* and *Gad1* counts per cell in excitatory and inhibitory neurons. (**D**) UMAP plot of the gene expression of inhibitory neurons. Colors and labels indicate H2 types of inhibitory neurons. (**E**) UMAP plot of the gene expression of excitatory neurons. Colors indicate slice numbers. The coordinates of dots in the UMAP plot are the same as those in Fig. 2C. (**F**) Cluster correspondence between non-isocortical H3 types in BARseq (rows) and single-cell RNAseq (columns) (Yao et al. 2021b). (**G**)(**H**) Overlap between isocortical (G) and non-isocortical (H) H3 types and CCF-defined areas.

**ED Figure 3.**
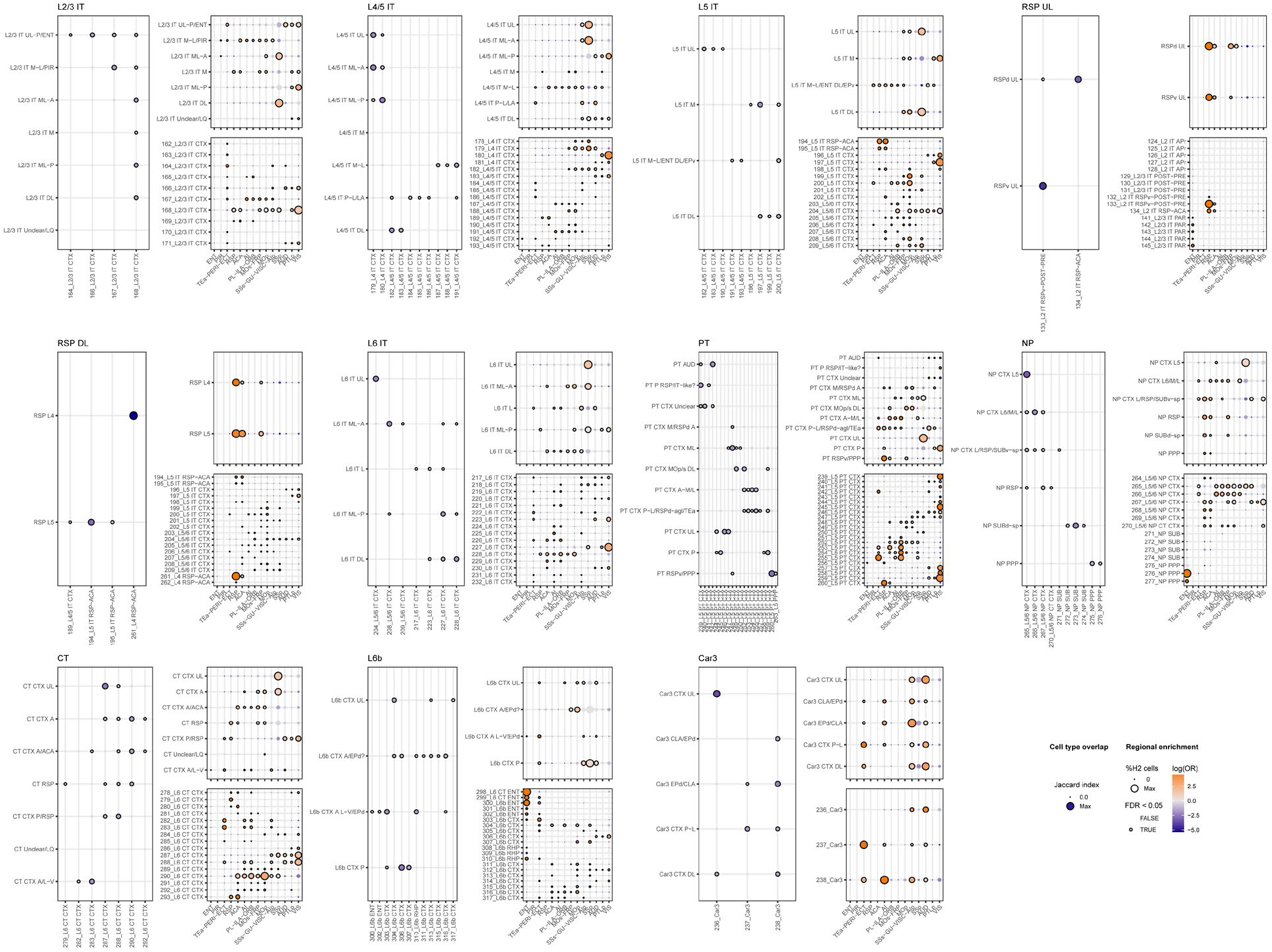
Mapping and comparative regional enrichment of BARseq and scRNAseq types. For each BARseq H2 type, we show the mapping of BARseq H3 types with reference scRNAseq type (*left*), the CCF enrichment of H3 types (*top right*), and the CCF enrichment of scRNAseq types (*bottom left*). The mapping between BARseq and scRNAseq types is quantified as the Jaccard Index, significant associations (permutation test) are shown by outlining dots with black circles. The regional enrichment is quantified as odds ratios, significant deviations (hypergeometric test) are shown by outlining dots with black circles. In the mapping panel, all BARseq H3 types are shown but, for readability, only scRNAseq types with significant associations are plotted. In contrast, the CCF enrichment is shown for all scRNAseq types belonging to subclasses that are equivalent to the BARseq H2 type (e.g., the BARseq L4/5 IT type corresponds to the L4 IT and L4/5 IT subclasses in the scRNAseq dataset). Colors indicate log odds ratios and circle size indicates the fraction of cells among all cells of that H2 type.

**ED Figure 4.**
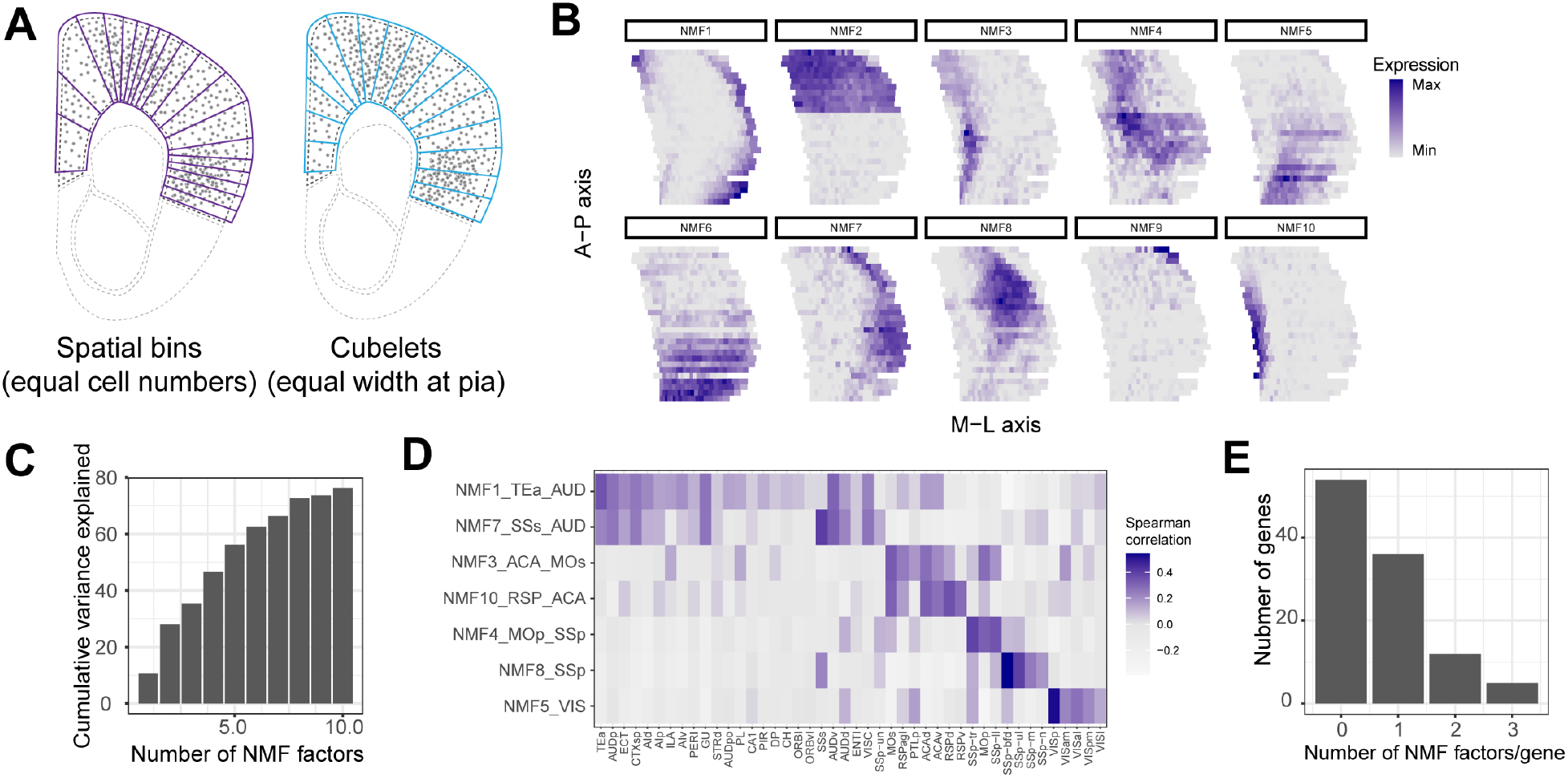
Identifying shared spatial patterns of gene expression using non-negative matrix factorization (A) Illustrations of the definition of “spatial bins” used in gene expression analyses (purple outlines) and “cubelets” used in the analyses of H3 type distributions (blue outlines) in the pilot brain. The definition of spatial bins aimed for equal cell numbers across bins within a slice, whereas the definition of cubelets aimed for equal width on the surface of the cortex. Dots indicate cells. (**B**) Spatial patterns of all 10 NMF factors. (**C**) Cumulative variance explained by the indicated number of NMF factors. (**D**) Spearman correlation between NMF factors and cortical areas. (**E**) Histogram of the number of NMF factors that each gene is associated with.

**ED Figure 5.**
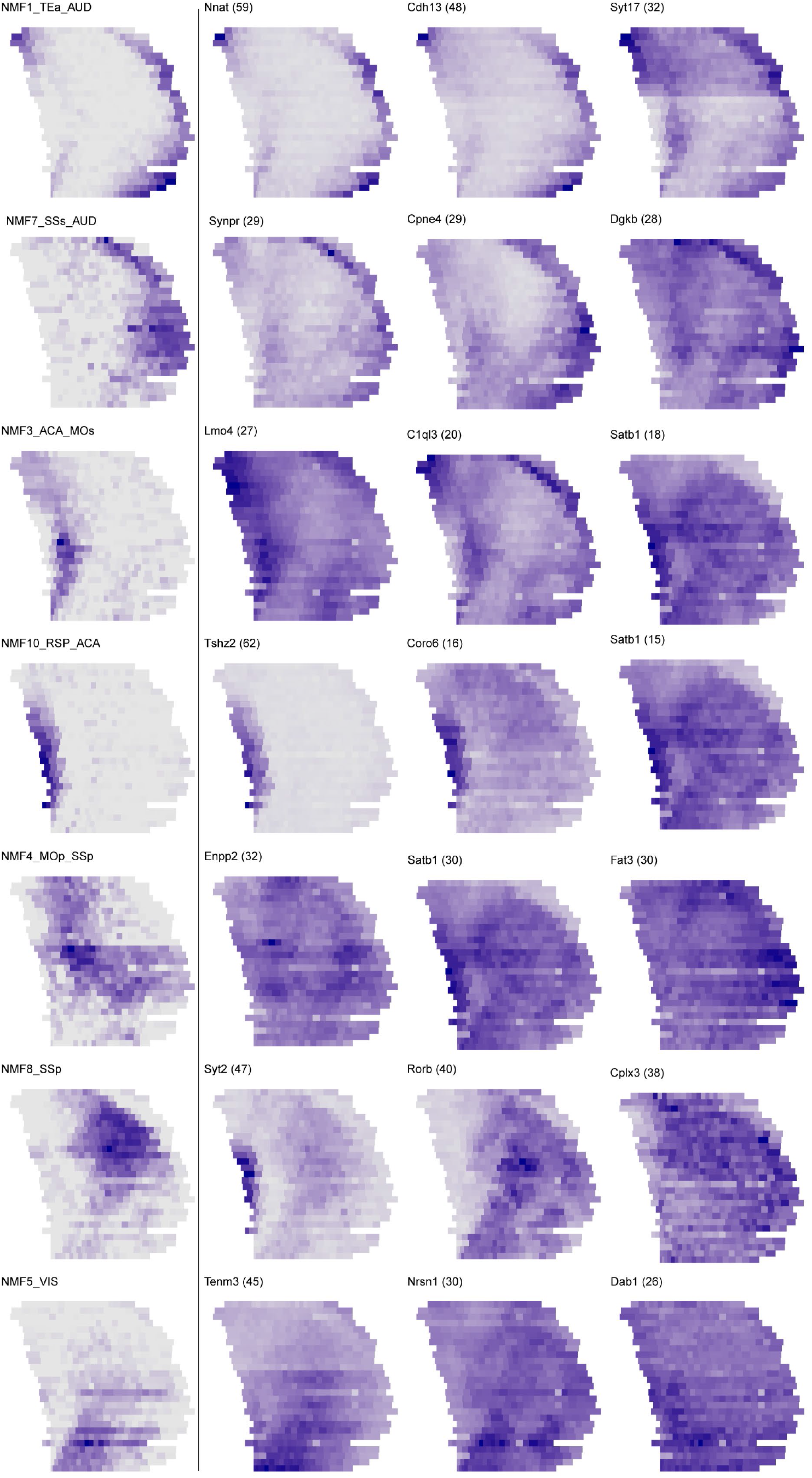
Expression patterns of top genes associated with each NMF spatial pattern. The left column shows the pattern associated with each NMF component; the right column shows the overall expression patterns (total expression counts across all cortical cells) of the top 3 genes associated with each NMF component. Expression patterns were min-max standardized (max expression = blue). Numbers in parentheses next to gene names show the average percentage of gene expression variance explained by the NMF pattern across cortical H2 types.

**ED Figure 6.**
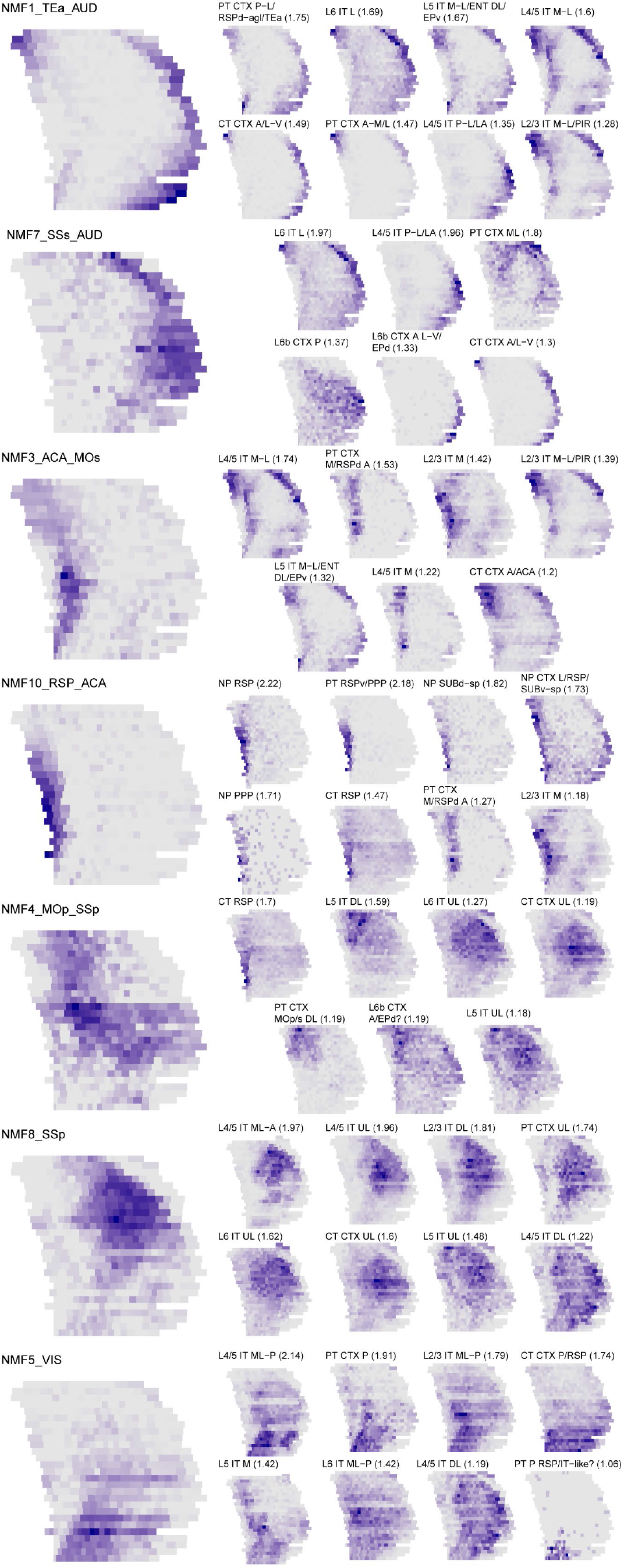
Distribution patterns of top H3 types associated with each NMF spatial pattern. The left column shows the pattern associated with each NMF component; the right column shows the distribution pattern (fraction of cells from the H3 type found in each bin) of H3 types showing above-null association with the NMF pattern. Numbers in parentheses next to H3 type names show the scaled Spearman correlation between the NMF pattern and the distribution pattern.

**ED Figure 7.**
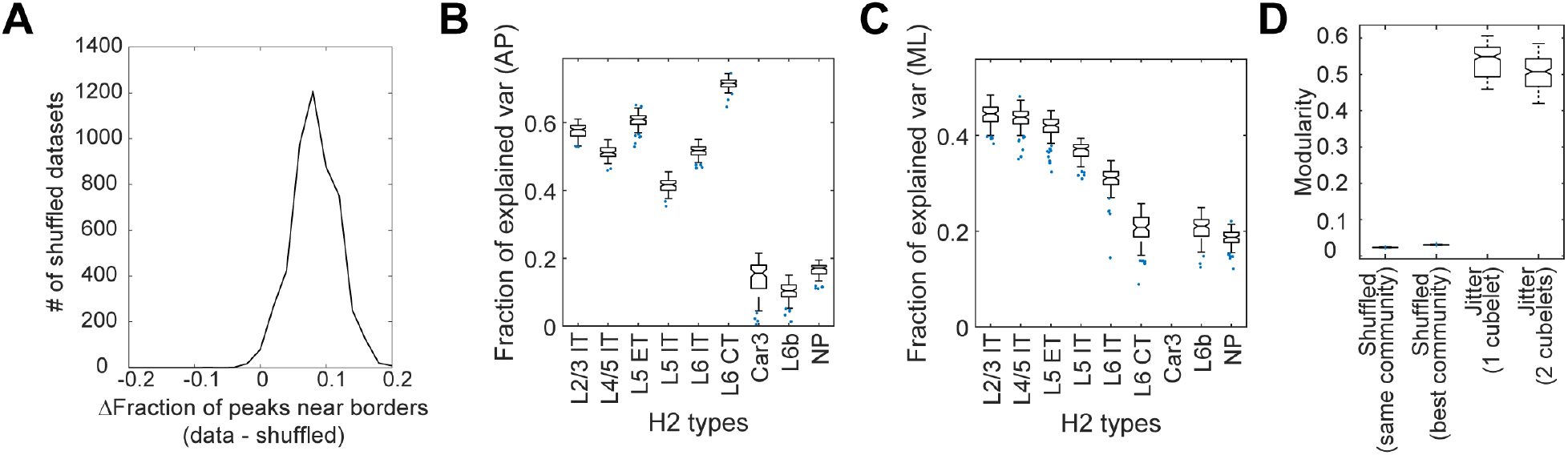
Cortical areas are distinct in H3 type composition. (**A**) The positions of abrupt changes in the composition of H3 types were shuffled randomly within each slice, and the difference in the fractions of positions that were close to a CCF area border between the real data and shuffled data was calculated (see **Methods**). Positive values indicate that abrupt changes in the composition of H3 types were more likely to be associated with area borders in real data than in shuffled control. This shuffling was repeated 5,000 times, and the distribution of this difference is plotted in a histogram. (**B**)(**C**) The composition of H3 types within each indicated H2 types (x-axes) were used to predict the AP (B) and ML (C) locations of a cubelet. For each H2 type, we performed 100 trials. In each trial, we randomly held 10% of data as test set to determine the fractions of variance explained. (**D**) The distribution of modularity of shuffled data, or data with 1-2 cubelets of jitter in CCF registration. For shuffled data, we calculated modularity based on either the same clusters obtained from real data, or by the best clusters obtained by Louvain community detection on the shuffled data.

**ED Figure 8.**
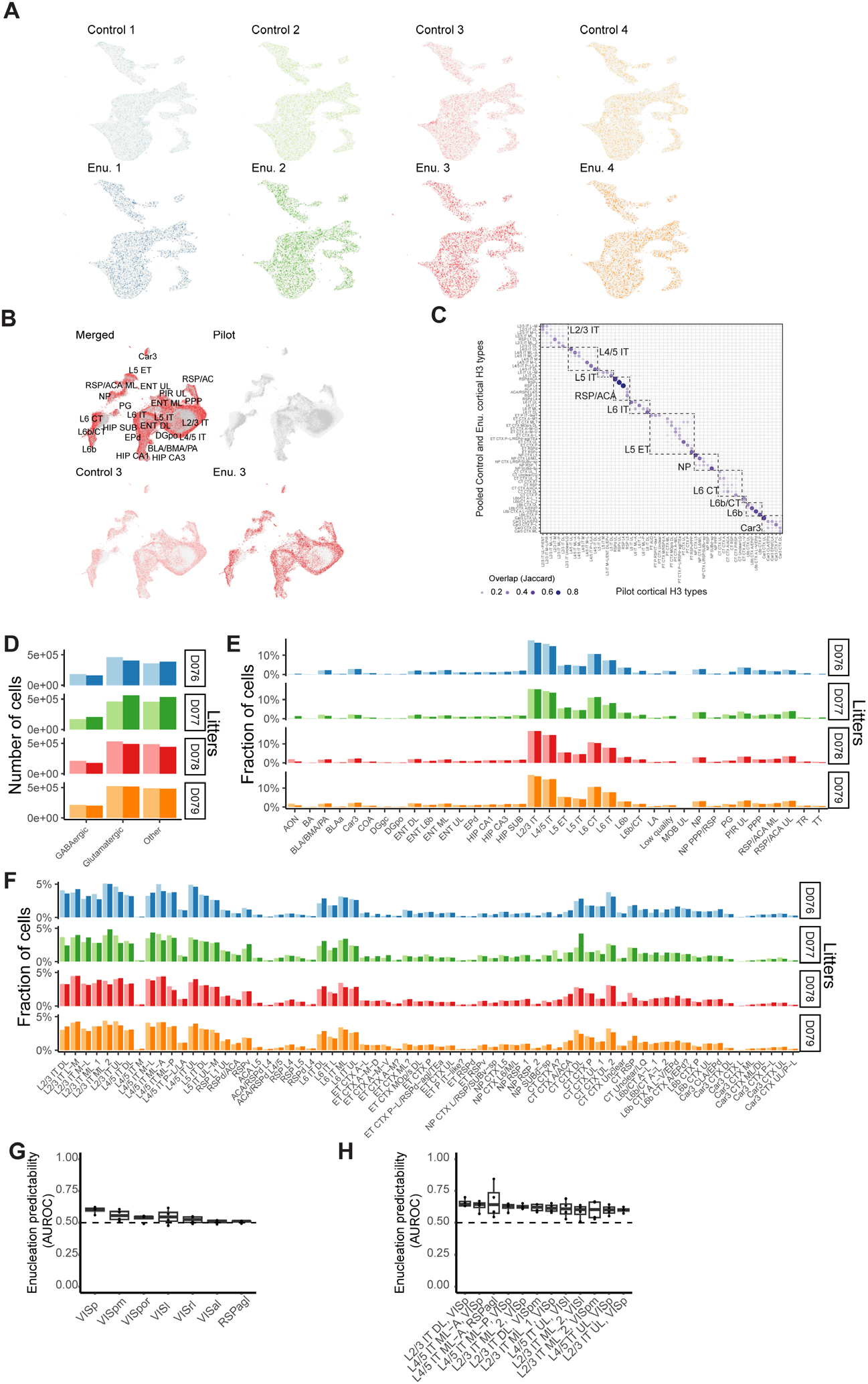
H3 types were consistent between the control brains and the enucleated brains. (**A**) UMAP plots of the gene expression of neurons from all eight littermate brains. In each plot, neurons from the indicated brain are colored and neurons from all other brains are shown in gray. (**B**) UMAP plot of all excitatory neurons from the pilot brain with excitatory neurons from the eight littermate brains projected onto the same UMAP coordinate space. Merged data on top left shows neurons from the pilot brain in gray and all eight littermate brains in red. Top right shows only neurons from the pilot brain in gray. Bottom row shows excitatory neurons from one pair of littermates. (**C**) Correspondence between H3 types from the eight animals to H3 types in the pilot brain. Dot sizes and colors indicate Jaccard index. Dashed boxes indicate the parent H2 types. (**D**)(**E**) Fractions of cells belonging to each cortical H1 type (D) and H2 type (E) in all paired littermate brains. (**F**) Fractions of cells belonging to each cortical H3 type in all paired littermate brains. In all fraction plots, enucleated animals are represented by the darker color. (**G**) The AUROC scores of a nearest neighbor classifier that predicts the condition (control or enucleated) of a neuron in the indicated cortical areas. Only areas with performance over 0.5 for at least 3 out of 4 litters were shown. Boxes indicate quartiles and medians, and whiskers indicate range. Dots indicate performance for each held-out litter. (**H**) The AUROC scores of a nearest neighbor classifier that predicts the condition (control or enucleated) of a neuron of the indicated H3 types in the indicated cortical areas. Only combinations with moderate performance (> 0.6) were shown. Boxes indicate quartiles and medians, and whiskers indicate range. Dots indicate performance for each held-out litter. Enu, enucleated.

**ED Figure 9.**
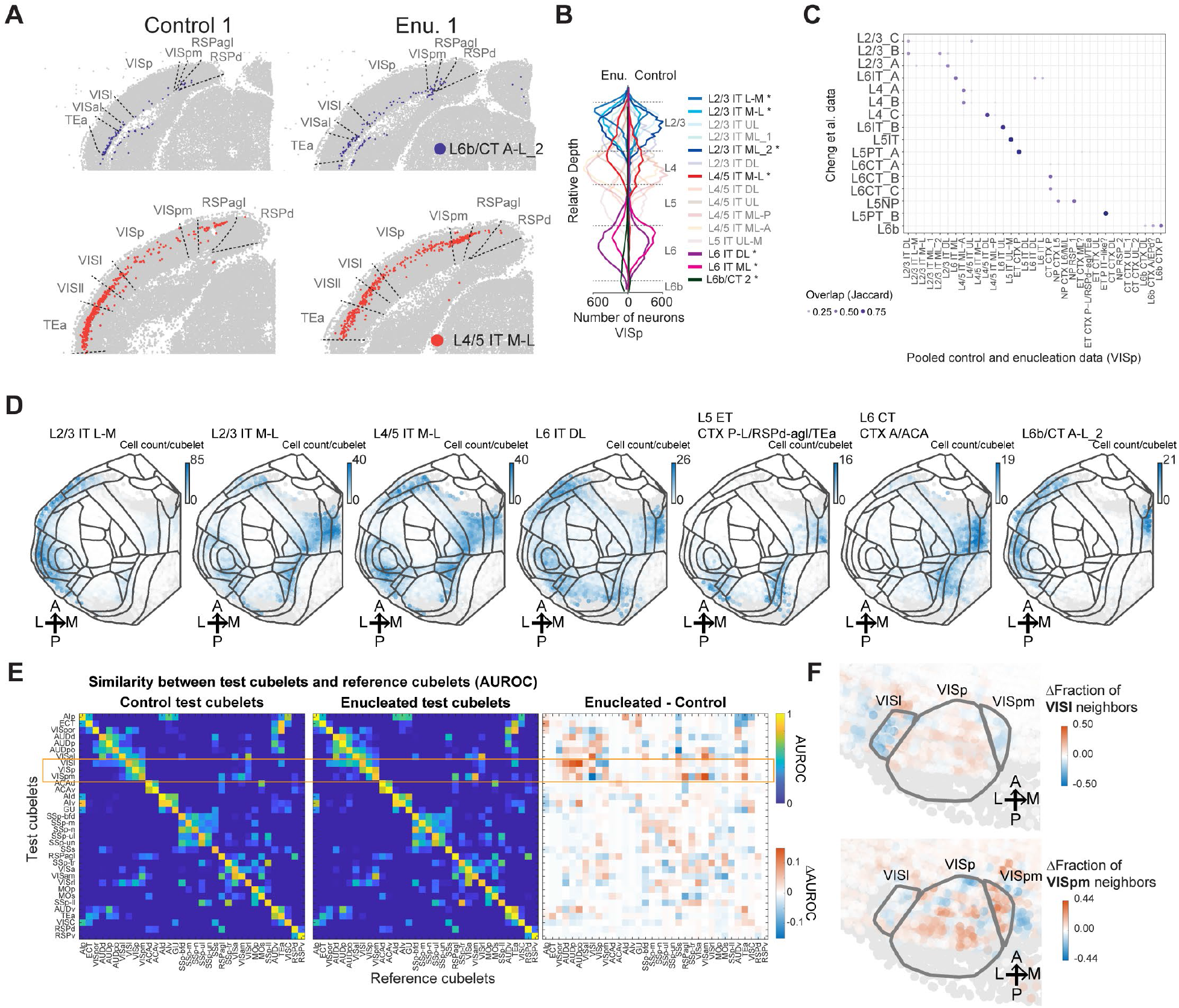
Enucleation broadly shifted visual area neurons to H3 types in medial and lateral areas. (**A**) Example slice images for L6b/CT A-L_2 and L4/5 IT M-L in a representative littermate pair. (**B**) The laminar distributions of H3 types in VISp. H3 types that were enriched or depleted are shown in dark colors. (**C**) Cell type mapping (Jaccard index) of cortical H3 types in the enucleated and control littermates to cell types in Cheng et al. (2022). (**D**) The number of cells per cubelet for the top enriched H3 types in VISp. Colors indicate cell counts in each cubelet. (**E**) AUROC of a nearest neighbor classifier assigning cubelets from control (left) or enucleated (middle) brains to cubelets in reference control brains. The difference in AUROC between the enucleated and the control brains are shown on the right. Orange box highlights the relevant VIS areas (VISp, VISpm, and VISl). (**F**) Magnified views of flatmaps showing the enriched or depleted fraction of VISl (top) or VISpm (bottom) neighbors for each cubelet. The circled areas indicate VISl, VISp, and VISpm. Enu, enucleated.

