## Supplementary Note for "Whole-cortex in situ sequencing reveals peripheral input-dependent cell type-defined area identity"

### Supplementary Information

#### *Supplementary Note 1: Gene panel selection and optimization*

We selected marker genes based on 47 sets of single-cell RNAseq datasets collected in adult mouse brains, including seven datasets in the motor cortex (MOp) using different single-cell RNAseq techniques (Yao, Liu, et al., 2021) and 40 datasets in the whole cortex and hippocampus using SmartSeq and 10x v3(Yao, van Velthoven, et al., 2021). Multiple datasets allowed us to pick genes that were differentially expressed across cell types consistently across techniques. This consistency thus increases the likelihood that it would also be differentially expressed when detected using BARseq, which has different constraints and limitations from all single-cell RNAseq techniques. We focused on picking highly expressed combinatorial marker genes with large fold change between cell types instead of binary markers. Specifically, we automatically aggregated differential expression statistics using MetaMarkers (Fischer & Gillis, 2021), then restricted panel selection to genes with an average AUROC>0.8, an average detection rate (fraction of cells expressing the gene)>0.9, and an average fold change>2. We then manually selected potential gene panels based on observed expression patterns. We iterated through several panel designs with increasing numbers of genes and assessed their cell typing power using published single-cell RNAseq data. Specifically, we applied MetaNeighbor to a surrogate dataset that had comparable sensitivity to BARseq (10x v2 single-nuclei RNAseq from the MOp) and computed our ability to distinguish existing cell types (cell type separability, MetaNeighbor with 5-fold cross-validation, **ED Fig. 1A, C**) and our ability to retrieve cell types through de novo clustering (cell type clusterability, CPM-log1p normalization, kNN-Louvain clustering, MetaNeighbor against other MOp datasets, **ED Fig. 1B**). Increasing the number of genes generally improved cell typing performance, which largely plateaus with the 104-gene panel we used.

We further experimentally assessed whether the 104-gene panel (**Supplementary Table 1**) was sufficient. To do so, we performed BARseq to interrogate a 137-gene panel on two coronal sections; this gene panel included the whole 104-gene panel and an additional 33 genes that were selected following similar criteria. We then compared clusters obtained using either all genes, or the 104-gene subset from the same dataset. We found that the two gene panels resolved similar clusters at the H2 and H3 levels, and the separation of clusters was also similar (**ED Fig. 1D-F**). We thus concluded that the 104-gene panel was sufficiently optimized for resolving cortical excitatory types in the current experiments.

#### *Supplementary Note 2: Estimation of cell segmentation errors*

Segmentation errors in *in situ* sequencing can produce artifacts in single-cell gene expression in at least two ways. First, segmentation may exclude a portion of a cell. This type of error results in subsampling of all transcripts in the cell and potentially result in lower quality of single-cell data. To exclude these low-quality cells, we use a genes-per-cell threshold and a counts-per-cell threshold to remove cells with insufficient gene expression data. Second, segmentation may combine two neighboring cells into one cell, which is reminiscent of doublets in single-cell RNAseq experiments. To estimate doublet rate, we examined the expression of *Gad1* and *Slc17a7*, two highly expressed and mutually exclusive marker genes for cortical inhibitory neurons and cortical excitatory neurons, respectively, in both inhibitory neurons and cortical excitatory neurons. Consistent with the mutual exclusivity, the excitatory and inhibitory neurons showed distinct distribution of these two genes (**ED Fig. 2B, C**). We first calculated the median expression level for each marker gene in their corresponding cell class. We then asked what fraction of cells in the two H1 types had expression that was equal to or higher than this median expression level. The doublet rate is defined as the ratio between the fractions of cells with above-median expression in the two H1 types, divided by the fraction of the H1 type in which the marker is normally expressed [i.e. $P(Gad1 >$

$median(Gad1)|Excitatory)/P(Gad1 > median(Gad1)|Inhibitory)/P(Inhibitory/All\ neurons)$ and $P(Slc17a7 > median(Slc17a7)|Inhibitory)/P(Slc17a7 > median(Slc17a7)|Excitatory)/P(Excitatory/All\ neurons)$ ]. The estimated doublet rate is 7% based on *Slc17a7*, and 5% based on *Gad1*. The value based on *Slc17a7* is likely an overestimate, because many *Slc17a7* transcripts are found outside of neuronal somata, which could inflate the estimated doublet rate.

In *in situ* sequencing experiments, the doublet rate is heavily dependent on cellular density and may vary widely across brain regions. Thus, our estimates of doublets, which were based on cortical neurons, may not reflect doublet rates in other brain regions. However, because our analyses focused on the cortex, our cortex-based estimates were appropriate.

***Supplementary Note 3: Assessing differential expression of marker genes across H2 types***

We used MetaMarkers to identify the most differentially expressed markers across H2 types (**Fig. 2D**). Reassuringly, many identified marker genes for H2 types coincided with marker genes identified in previous studies. Whenever possible, we showed previously identified markers in three studies, which we abbreviated as T18(Tasic et al., 2018), Y21a(Yao, Liu, et al., 2021), and Y21b(Yao, van Velthoven, et al., 2021). The other markers were identified as strong markers during panel selection (see **Supplementary Note 1**), and were labeled as “strong panel genes”, along with the dataset used to identify the marker. The genes we selected for **Fig. 2D** are: *Slc30a3* (known pan-IT marker, T18), *Cux2* (known L2/3-L4/5 IT marker, Y21b), *Rasgrf2* (strong L2/3 IT panel gene, based on Y21a data), *Rorb* (known L4/5 IT marker, Y21a), *Etv1* (known pan-L5 marker - L5 IT, NP, PT, Y21b), *Scnn1a* (known deep layer RSP marker, Y21b), *C1ql3* (strong L6 IT panel gene, based on Y21a data), *Fezf2* (known non-IT marker, except L5 IT, Y21a), *Rab3c* (strong L5 ET panel gene, based on Y21a data), *Tle4* (known CT-NP-L6b marker, Y21b), *Tshz2* (known NP marker, T18, Y21b), *Foxp2* (known CT marker, T18, Y21a, Y21b), *Ctgf* (known L6b marker, Y21b), *Synpr* (strong Car3 panel gene, based on Y21a data). We also noted that RSP cells have known *Tshz2* and *Slc30a3* co-expression (Y21b)

***Supplementary Note 4: NMF curation.***

To extract spatial expression patterns that recurred across genes and H2 types, we applied non-negative matrix factorization (NMF) to pseudobulk expression data (one pseudobulk vector per spatial bin and H2 type), treating spatial bins as features (see **Methods**). To select a suitable number of NMF components, we progressively increased the number of components until we saw the appearance of components with negligible contributions to the variance explained (covering a small number of spatial bins, unlikely to represent robust biological signal). We identified 10 NMF components (**ED Fig. 4B**), which explained 76% of the variance in gene expression across space. Component NMF2 is found only in the anterior sections, which was sequenced in a different batch from the posterior sections. NMF2 thus reflects the batch effect between the two batches of samples. NMF6 is found mostly in posterior sections and shows clear strip-like patterns. Because each strip corresponds to a coronal section, and we expect gene expression to vary smoothly across adjacent coronal sections, NMF6 appears to capture differences in gene expression associated with cryo-sectioning differences across sections. Because these two components largely reflect technical variations in gene expression, we excluded them from further analyses. In addition, NMF9 is found only in the lateral edge of the most anterior sections. This area corresponds to the transition among the orbitofrontal cortex, the piriform cortex, and the endopiriform nucleus. Because of the area specificity and the continuity of its spatial distribution, this component likely reflects gene expression that is specific to this transition area. Nonetheless, because this component was found in only a small subset of spatial bins, we also excluded NMF9 from subsequent analyses.

#### *Supplementary Note 5: Data quality comparison across brains*

To assess the consistency between the pilot brain and the four littermate pairs, we co-embedded the two datasets in the same UMAP space (**ED Fig. 8B**). The two datasets largely overlapped, but data for the four littermates were systematically "shifted" so that different clusters were more distinguishable compared to the pilot brain. This pattern of the shift between the two datasets is consistent with an overall change in sensitivity and data quality. To formally assess clusters between the two datasets, we mapped H3 cortical types between the pilot brain and the combined new dataset of eight brains (**ED Fig. 8C**) and found 1:1 or 1:2 cell type mapping between them for all clusters. Additionally, the H3 types from the eight brains were evenly distributed across all brains, indicating that H3 types were highly consistent across the eight animals (**ED Fig. 8D-F**). Thus, these results indicate that cell typing was consistent between the pilot brain and the eight brains, but that the eight brains had better data quality.

The better data quality achieved on the eight brains was not surprising and could reflect either biological differences or improvement in instrumentation. The data from the pilot brain were collected at Cold Spring Harbor Laboratory (Zador lab), and the data from the four littermate pairs were collected on an improved system at the Allen Institute (Chen lab). The improved system has an improved spinning disk confocal with larger field-of-view, large field-of-view cameras, higher-power lasers, and optimized filter sets compared to the original system at CSHL. Furthermore, we also improved the data registration pipeline and imaging settings between collection of these two datasets. Thus, the differences in instrumentation and data processing alone could account for the improved data quality of the four littermate pairs. In addition to the differences in instrumentation, the pilot brain was collected at P56 and the four littermate pairs were collected at P28. Based on the current data, we could not distinguish whether the improved data quality was due to improvements in instrumentation or due to differences in age.

#### *Supplementary Note 6: The effect of enucleation on gene expression*

H3 types were shared by neurons from both the control brains and the enucleated brains: for all H3 types, between 25% and 80% of neurons were from the control brains. To test whether there are changes at a finer granularity than the H3 types, we identified the 100 nearest neighbors of each neuron, and asked what fraction of those neurons were from the control brains. For all neurons, 14 – 90% of neighboring neurons were from the control brains, which was consistent with the composition of H3 types. These results suggest that enucleation did not produce neighborhoods of neurons that were absent in the control animals.

We next tested if subpopulations of neurons within H3 types were preferentially enriched in neurons from the enucleated brains or the control brains by treating the kNN structure as a classifier: For each neuron, we predicted its condition of origin (control or enucleated) using its 100 closest neighbors. At a global level, the two conditions were uniformly mixed on average (AUROC=0.49). The classifier, however, was weakly predictive in seven areas (including visual areas and RSPagl) in at least 3 out of 4 litters (visual areas and RSPagl, **ED Fig. 8G**), with VISp showing the clearest difference (AUROC=0.61). Within H3 types, 12 combinations showed moderately positive performance using the neighbor-based classifier (AUROC > 0.6,

**ED Fig. 8H**), all involving L2/3 or L4/5 IT types. This moderate performance suggests that either heterogenous subpopulations within these H3 types were non-uniformly depleted or enriched, or that enucleation created new cell states within the existing H3 types. Whether such heterogeneity reflects the creation of new cell types relies on distinguishing transcriptomic signatures of cell types versus cell states, which is hotly debated in the field and cannot be settled with our current dataset. Thus, we did not further analyze the within-H3 type shifts in gene expression.

***Supplementary Note 7: Comparison to existing studies***

Several studies have previously examined the effect of removing visual stimuli on gene expression and cell types in the mouse visual cortex. In these studies, the effect of the stimulus removal varied according to how visual inputs were removed. Removal of sensory input generally falls into two categories: blocking thalamic axons from reaching cortical targets, or removing sensory input from the thalamic projections via sensory deprivation after the axons have reached the cortex. This study (bilateral enucleation at P1) falls into the second category.

Two previous studies (Cheng et al., 2022; Chou et al., 2013) perturbed visual thalamic inputs in ways that were most relevant to the perturbation we performed. Cheng et al. (2022) examined the effects of dark rearing during the visual critical period. This perturbation represented a similar but less severe sensory deprivation compared to bilateral enucleation performed in this study. Cheng et al. (2022) reported changes in gene expression in L2/3 IT neurons, corresponding to a change in cell type composition from three identified L2/3 IT cell types in visual cortex. To compare our results to their observations, we mapped H3 types in our dataset to cortical excitatory types in Cheng et al. (2022) (**ED Fig. 9C**) and found a strong correspondence across all types. For L2/3 IT neurons, three out of six H3 types in our data corresponded strongly to types L2/3 IT A, B, and C in Cheng et al. (2022). What is more, the sub-laminar distribution of matched cell types (**ED Fig. 9B**) also matched across our study and Cheng et al. (2022): Type L2/3 IT UL maps closely to type A in Cheng et al. (2022) and is in the superficial layer 2/3; L2/3 IT ML-2 corresponds to type B and is located in mid layer 2/3; L2/3 IT DL matches to both type B and C and is located in deep layer 2/3. Consistent with the findings of Cheng et al. (2022), we found that L2/3 IT ML-2 were reduced in VISp (**Fig. 7E**). In addition, we also observed broader changes in other IT neurons (L4/5 IT and L6 IT), which were not seen in Cheng et al. (2022). We speculate that the broader changes seen in our data reflected the earlier developmental timepoint at which enucleation was performed. Because IT neurons in deeper layers differentiate and presumably mature earlier, an earlier sensory deprivation would likely have a stronger effect on neurons in deeper layers than sensory deprivation during only the critical period, as performed in Cheng et al. (2022). Thus, our data recapitulated similar changes in L2/3 IT as observed in Cheng et al. (2022), but also revealed broader changes in other cell types that likely resulted from sensory deprivation starting from an earlier age.

In a second study, Chou et al. (2013) used a conditional knockout mutant to prevent most axons from the lateral geniculate nucleus (LGN) to reach the visual cortex during development, which eliminated the difference between primary visual cortex and higher visual areas. This change contrasts with our finding that the primary visual cortex remained distinct from secondary visual areas despite broad changes in H3 types. We speculate that these differences likely reflect the different types of perturbation performed in the two studies: Whereas Chou et al. (2013) blocked thalamic axons from reaching the cortex, our study only blocked sensory activity in those axons after they reached the cortex. These differences suggest that the physical connections established by thalamocortical axons are needed to define the primary visual cortex, and that the sensory activity conveyed through these axons plays a refinement role in cell type composition across both the primary visual cortex and neighboring higher visual areas.

*See SuppTableS1.xlsx*

***Supplementary Table 1 Primers and padlock probes used for BARseq***

| | Olympus IX81 | | | | Nikon Ti-2E (no excitation filter) | | |
|---|---|---|---|---|---|---|---|
| Channel | Laser | Excitation filter | Dichroic | Emission filter | Laser | Dichroic | Emission filter |
| G/YFP | 520 | zet443-518x | zt443-518rpc | FF01-565/24 | 514 | Zt405/514/635rpc | FF01-565/24 |
| T | 555 | zet402/468/555/640x | zt402/468/555/640rpc-ufs | ff01-585/11 | 561 | FF421/491/567/659/776-Di01 | FF01-441/511/593/684/817 |
| A | 640 | zet402/468/555/640x | FF652-Di01 | FF01-676/29 | 640 | Zt405/514/635rpc | FF01-676/29 |
| C | 640 | zet402/468/555/640x | FF652-Di01 | FF01-725/40 | 640 | Zt405/514/635rpc | FF01-775/140 |
| GFP | 470 | zet402/468/555/640x | zt402/468/555/640rpc-ufs | FF01-525/30 | 488 | FF421/491/572-Di01 | 69401m |
| DAPI | 405 | zet402/468/555/640x | zt402/468/555/640rpc-ufs | 69401m | 405 | FF421/491/572-Di01 | 69401m |
| TxRed | 555 | zet402/468/555/640x | zt402/468/555/640rpc-ufs | 69401m | 561 | FF421/491/572-Di01 | 69401m |
| Cy5 | 640 | zet402/468/555/640x | FF652-Di01 | FF01-676/29 | 640 | Zt405/514/635rpc | ZET532/640m |

***Supplementary Table 2. Microscopy filters and lasers used for BARseq***

*See SuppFig1.pdf*

***Supplementary Figure 1. Areal distribution of all H3 types***
