## Supplementary material for "Whole-cortex in situ sequencing reveals peripheral input-dependent cell type-defined area identity": Supp. Fig. 1

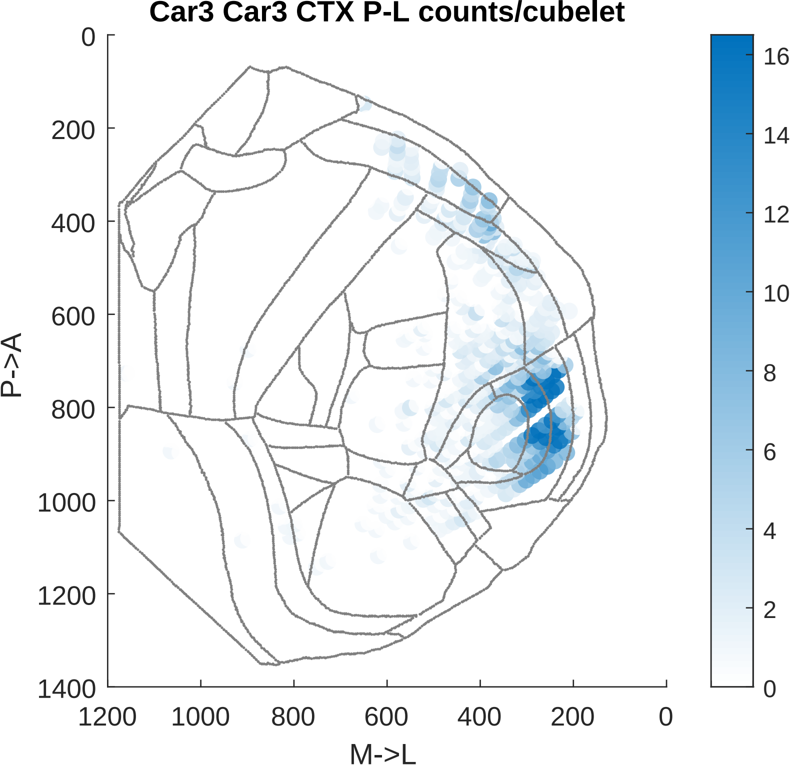

Car3 Car3 CTX P-L counts/cubelet
P->A
M->L
0
200
400
600
800
1000
1200
1400
1200
1000
800
600
400
200
0
16
14
12
10
8
6
4
2
0

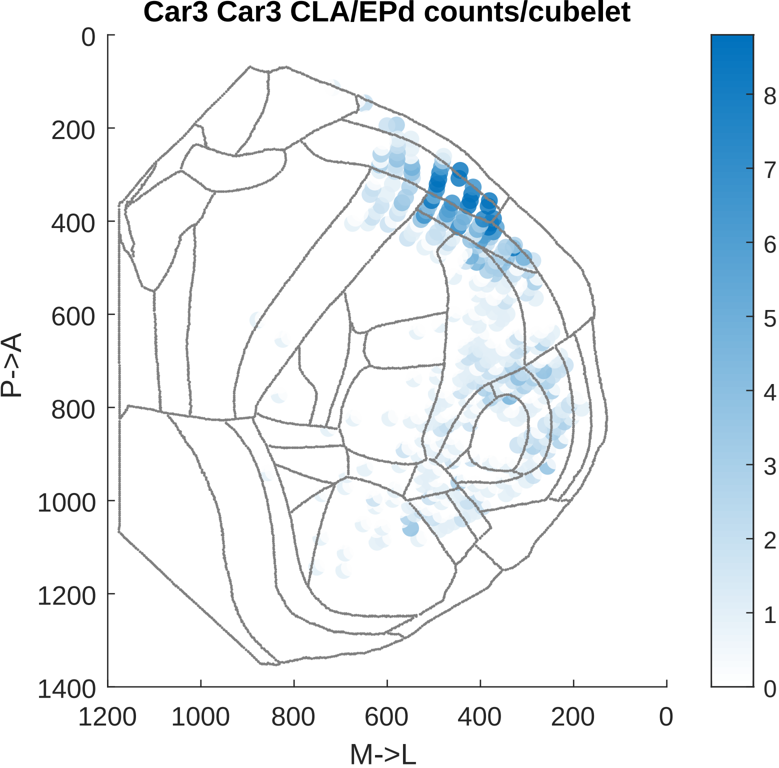

Car3 Car3 CLA/EPd counts/cubelet
0
200
400
600
800
1000
1200
1400
P->A
1200
1000
800
600
400
200
0
M->L
8
7
6
5
4
3
2
1
0

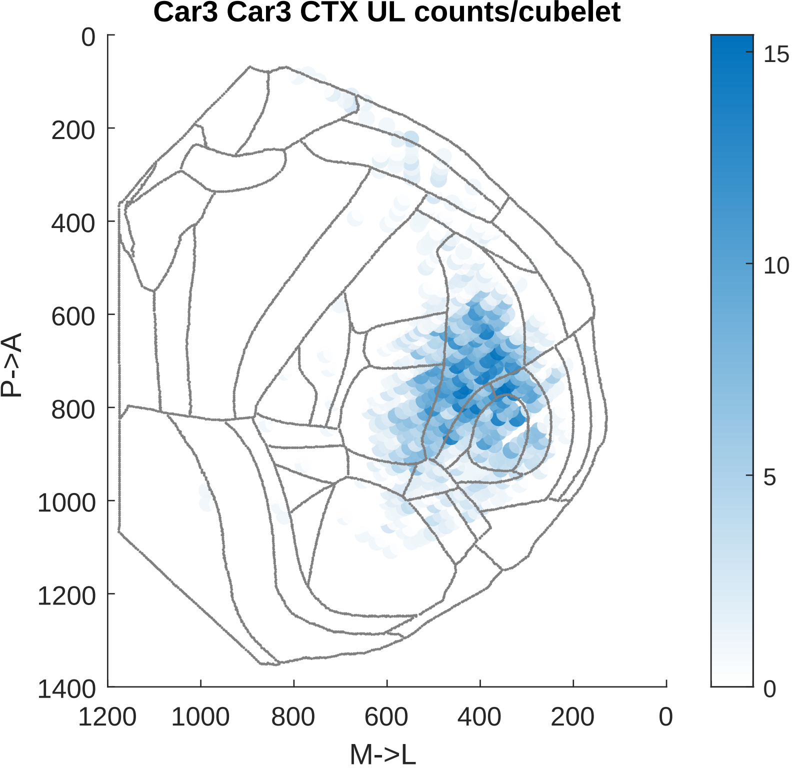

Car3 Car3 CTX UL counts/cubelet
P->A
M->L
0
200
400
600
800
1000
1200
1400
1200
1000
800
600
400
200
0
15
10
5
0

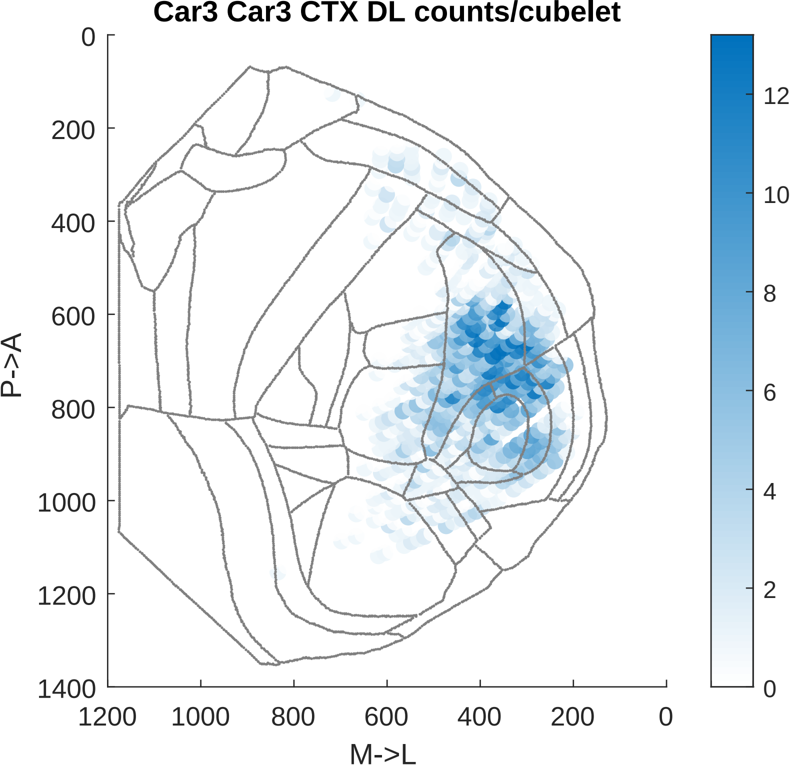
Car3 Car3 CTX DL counts/cubelet
P->A
M->L
0
200
400
600
800
1000
1200
1400
1200
1000
800
600
400
200
0
12
10
8
6
4
2
0

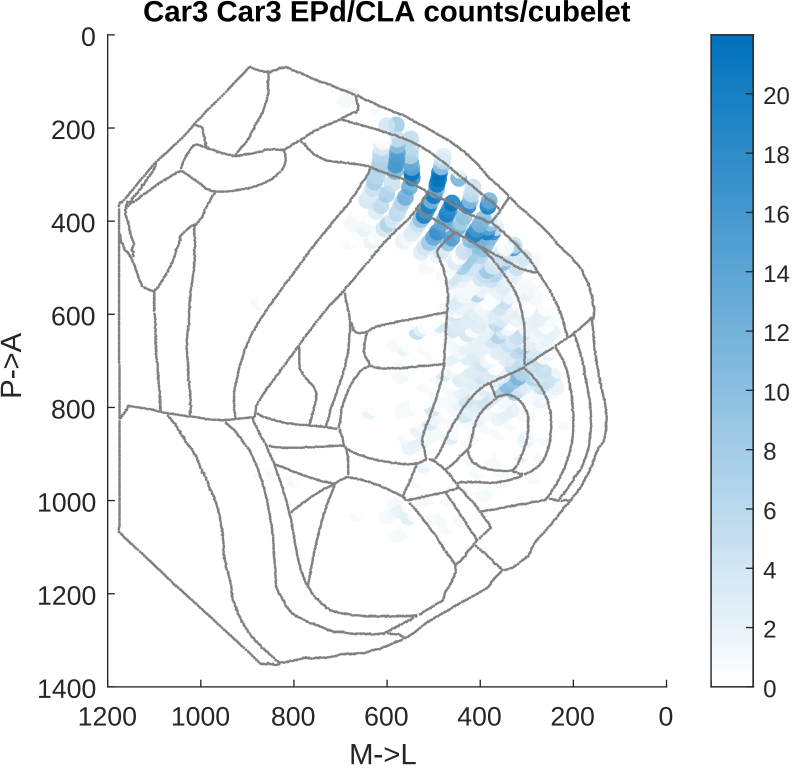

Car3 Car3 EPd/CLA counts/cubelet
0
200
400
600
800
1000
1200
1400
P->A
1200
1000
800
600
400
200
0
M->L
0
2
4
6
8
10
12
14
16
18
20

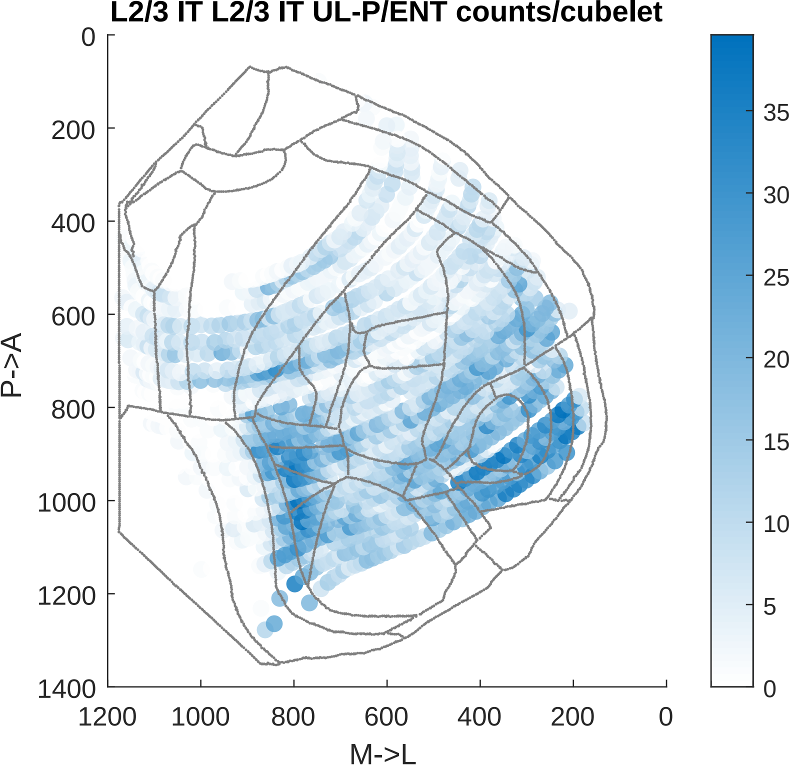

L2/3 IT L2/3 IT UL-P/ENT counts/cubelet
0
200
400
600
800
1000
1200
1400
P->A
1200
1000
800
600
400
200
0
M->L
35
30
25
20
15
10
5
0

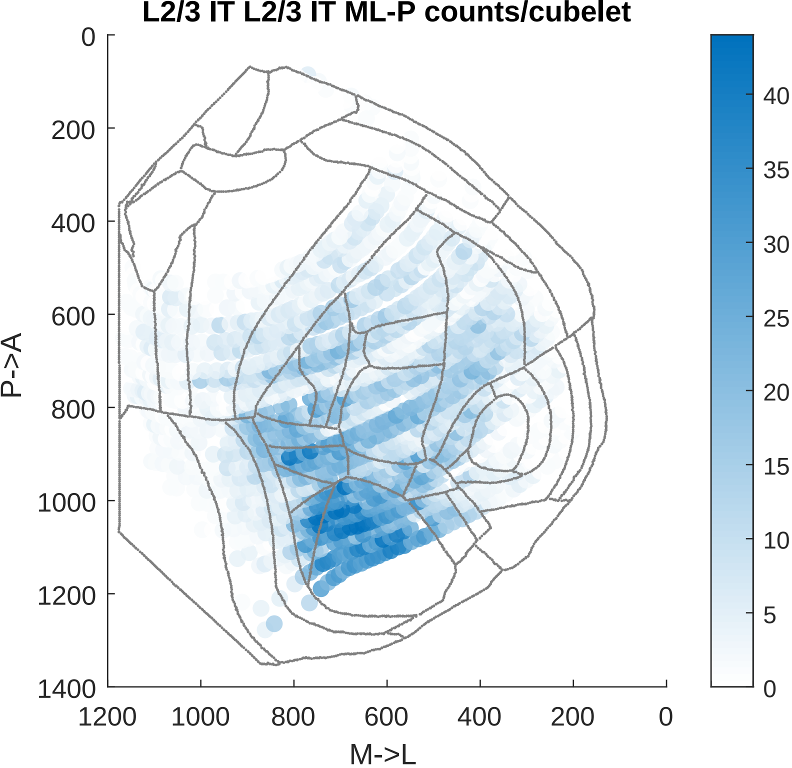
L2/3 IT L2/3 IT ML-P counts/cubelet
P->A
M->L
0
200
400
600
800
1000
1200
1400
1200
1000
800
600
400
200
0
40
35
30
25
20
15
10
5
0

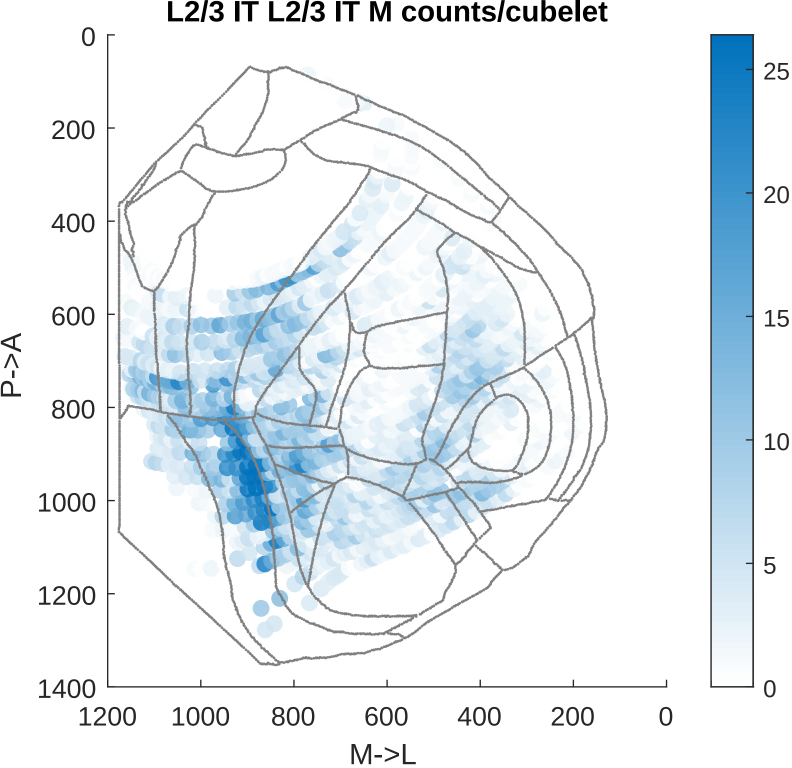

L2/3 IT L2/3 IT M counts/cubelet
P->A
M->L
0
200
400
600
800
1000
1200
1400
1200
1000
800
600
400
200
0
25
20
15
10
5
0

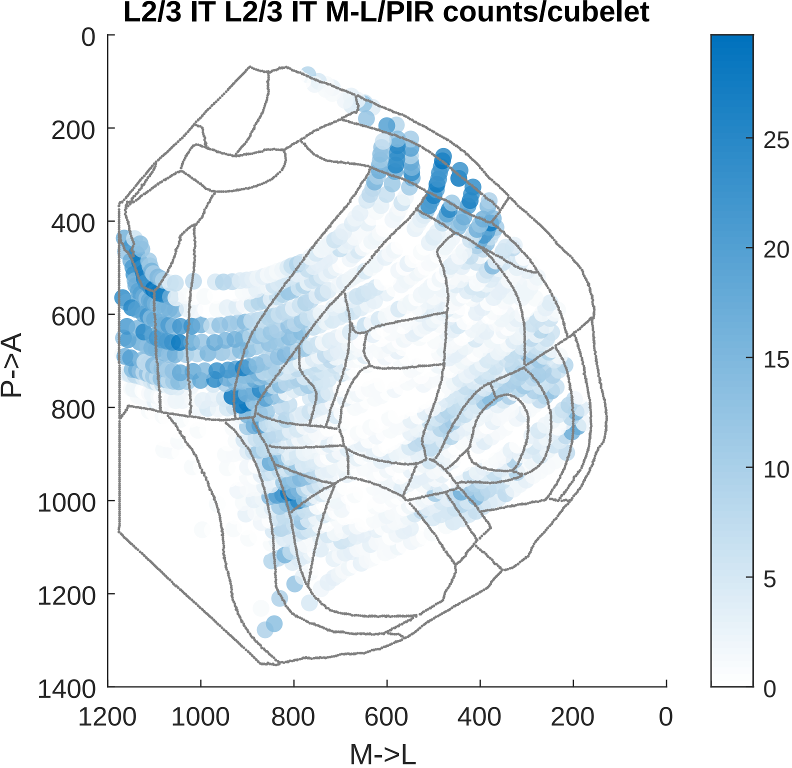

L2/3 IT L2/3 IT M-L/PIR counts/cubelet
P->A
M->L
0
200
400
600
800
1000
1200
1400
1200
1000
800
600
400
200
0
25
20
15
10
5
0

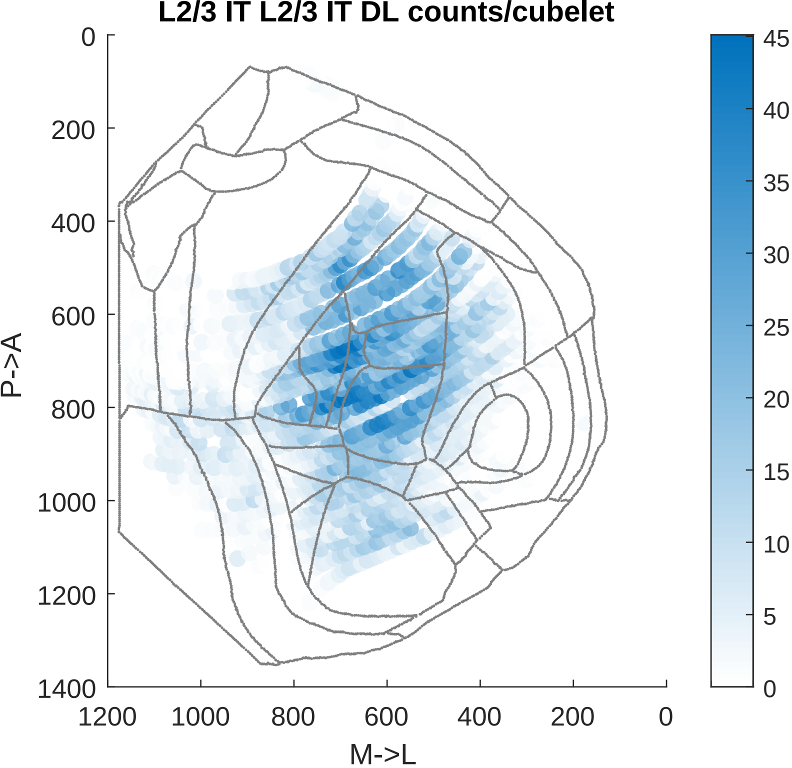

L2/3 IT L2/3 IT DL counts/cubelet
P->A
M->L
0
200
400
600
800
1000
1200
1400
1200
1000
800
600
400
200
0
45
40
35
30
25
20
15
10
5
0

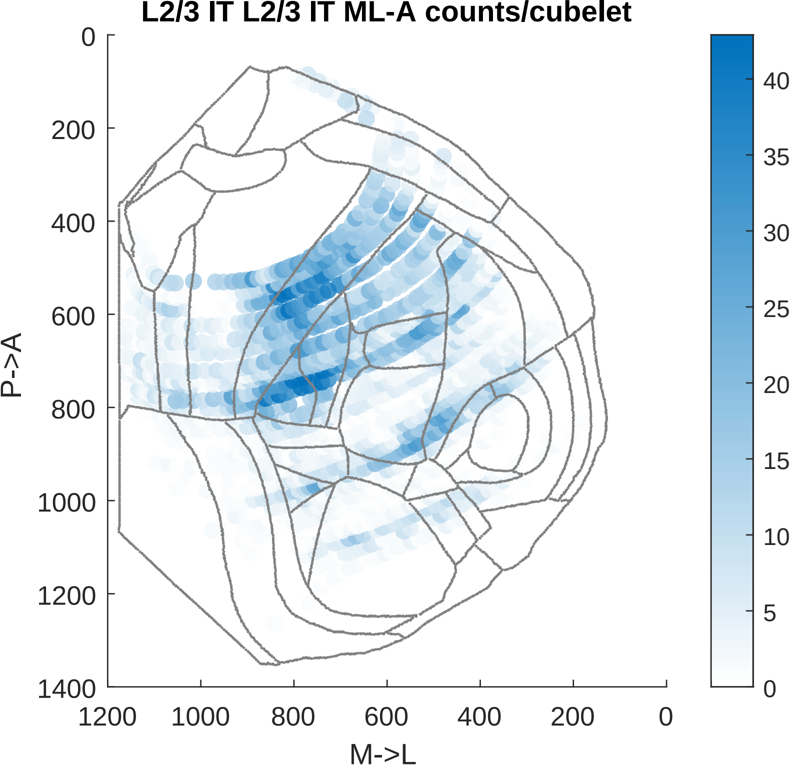

L2/3 IT L2/3 IT ML-A counts/cubelet
P->A
M->L
0
200
400
600
800
1000
1200
1400
1200
1000
800
600
400
200
0
40
35
30
25
20
15
10
5
0

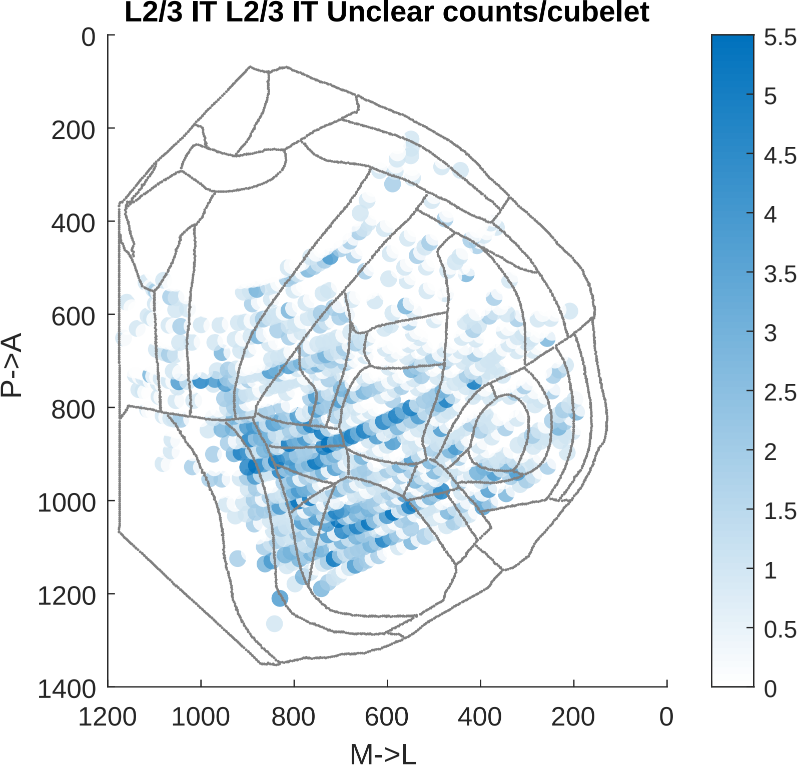

L2/3 IT L2/3 IT Unclear counts/cubelet
P->A
M->L
0
200
400
600
800
1000
1200
1400
1200
1000
800
600
400
200
0
5.5
5
4.5
4
3.5
3
2.5
2
1.5
1
0.5
0

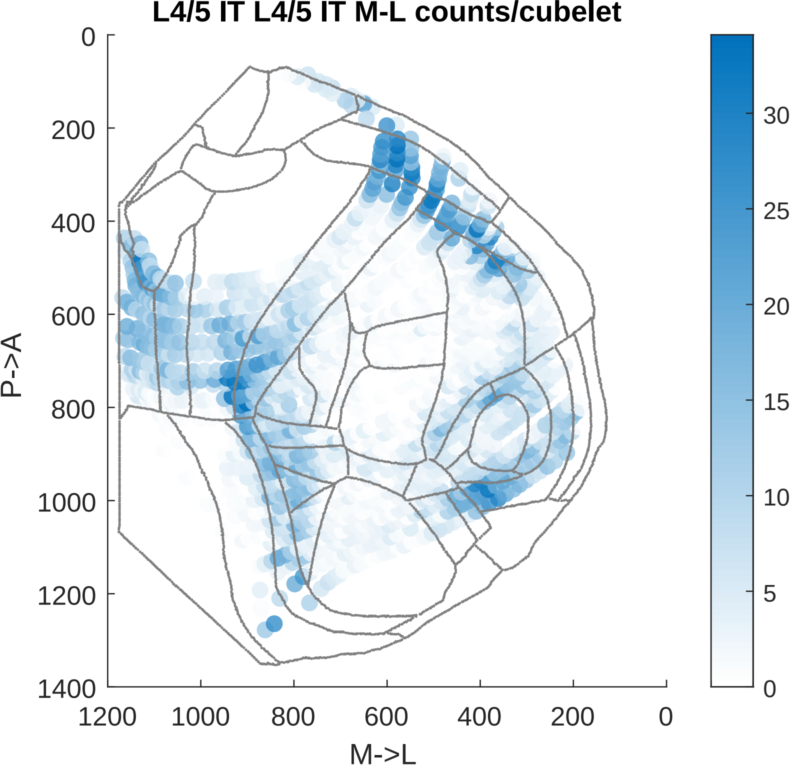
L4/5 IT L4/5 IT M-L counts/cubelet
P->A
M->L
0
200
400
600
800
1000
1200
1400
1200
1000
800
600
400
200
0
30
25
20
15
10
5
0

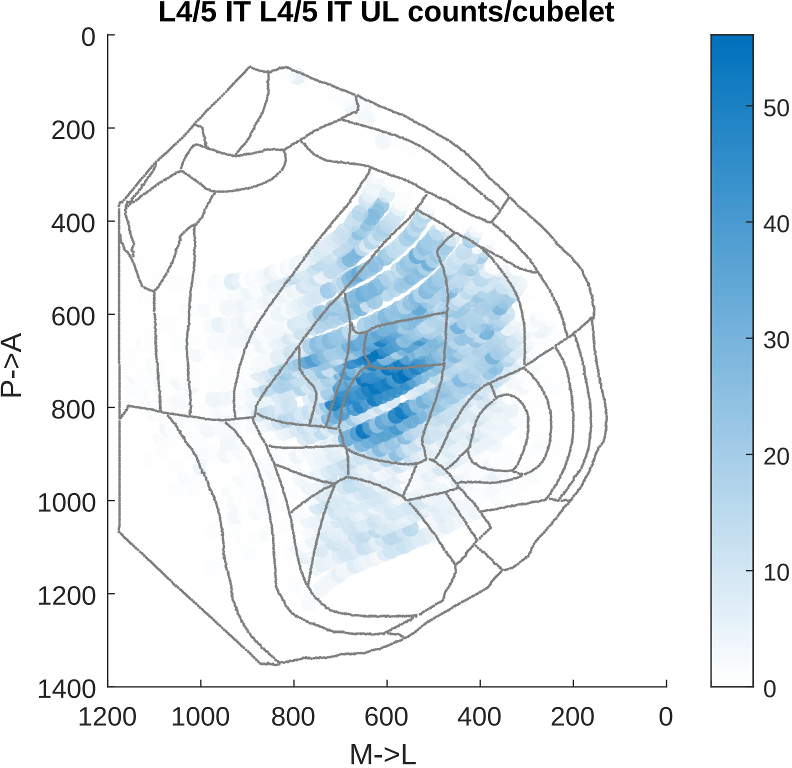
L4/5 IT L4/5 IT UL counts/cubelet
P->A
M->L
0
200
400
600
800
1000
1200
1400
1200
1000
800
600
400
200
0
50
40
30
20
10
0

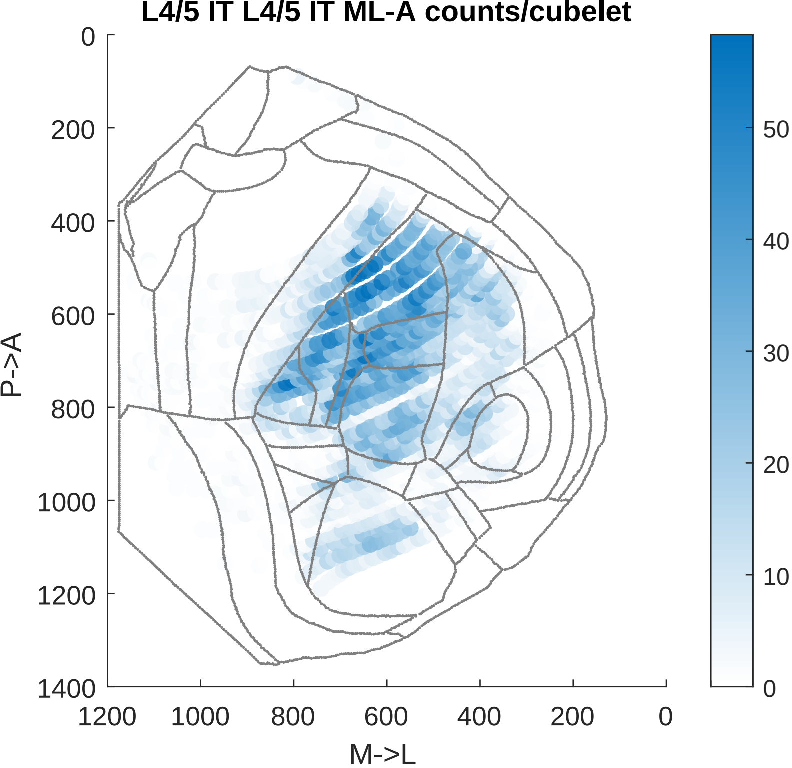

L4/5 IT L4/5 IT ML-A counts/cubelet
0
200
400
600
800
1000
1200
1400
P->A
1200
1000
800
600
400
200
0
M->L
50
40
30
20
10
0

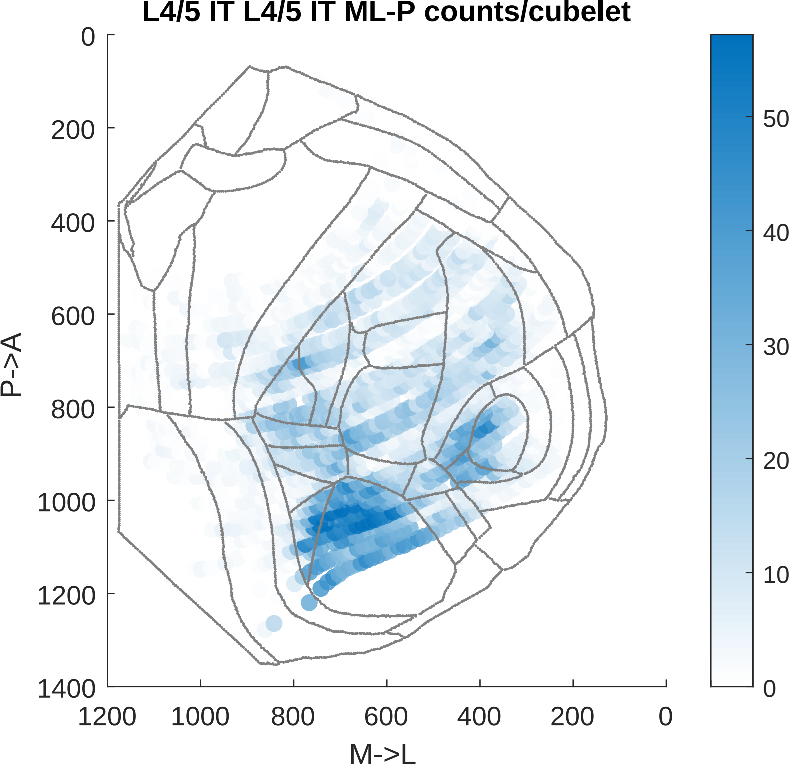

L4/5 IT L4/5 IT ML-P counts/cubelet
P->A
M->L
0
200
400
600
800
1000
1200
1400
1200
1000
800
600
400
200
0
50
40
30
20
10
0

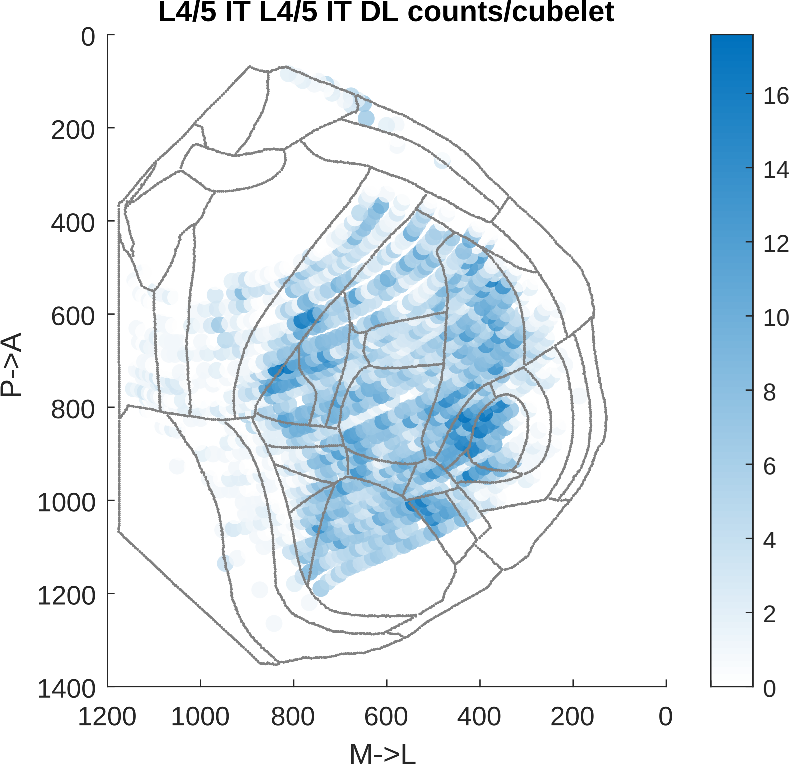

L4/5 IT L4/5 IT DL counts/cubelet
P->A
M->L
0
200
400
600
800
1000
1200
1400
1200
1000
800
600
400
200
0
16
14
12
10
8
6
4
2
0

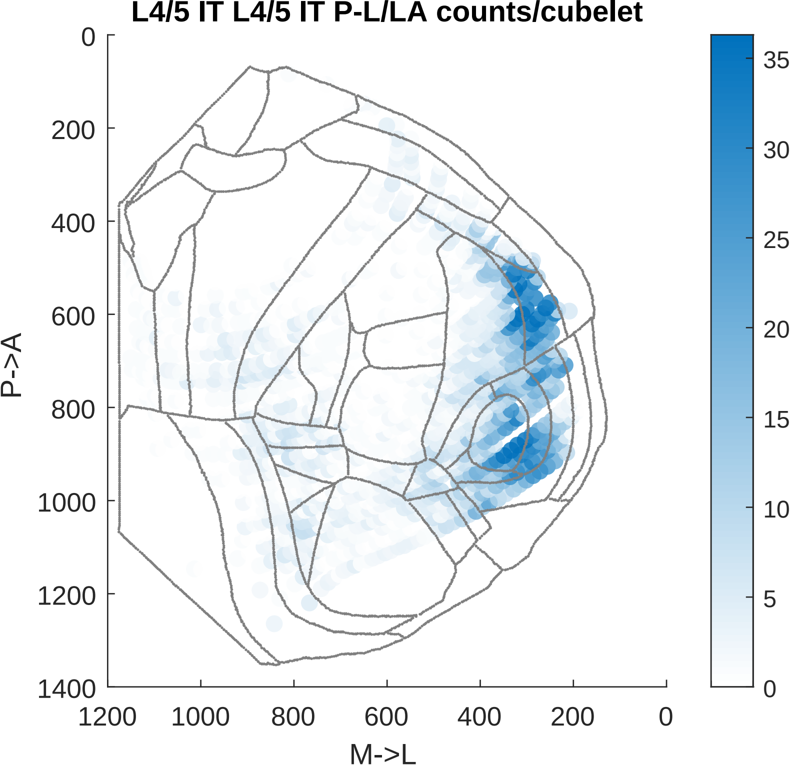

L4/5 IT L4/5 IT P-L/LA counts/cubelet
P->A
M->L
0
200
400
600
800
1000
1200
1400
1200
1000
800
600
400
200
0
35
30
25
20
15
10
5
0

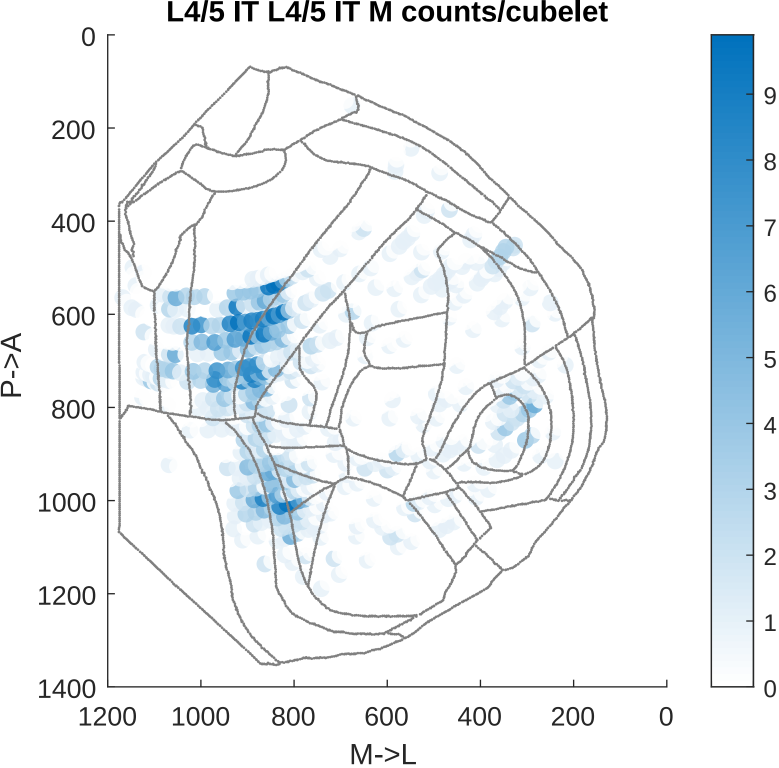

L4/5 IT L4/5 IT M counts/cubelet
0
200
400
600
800
1000
1200
1400
P->A
1200
1000
800
600
400
200
0
M->L
9
8
7
6
5
4
3
2
1
0

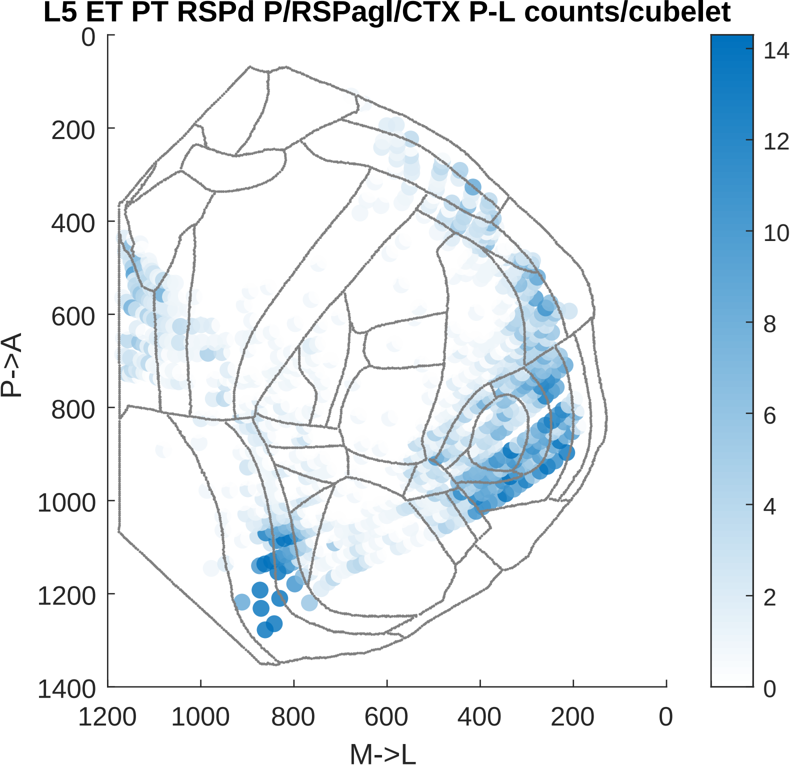

L5 ET PT RSPd P/RSPagl/CTX P-L counts/cubelet
0
200
400
600
800
1000
1200
1400
P->A
1200
1000
800
600
400
200
0
M->L
14
12
10
8
6
4
2
0

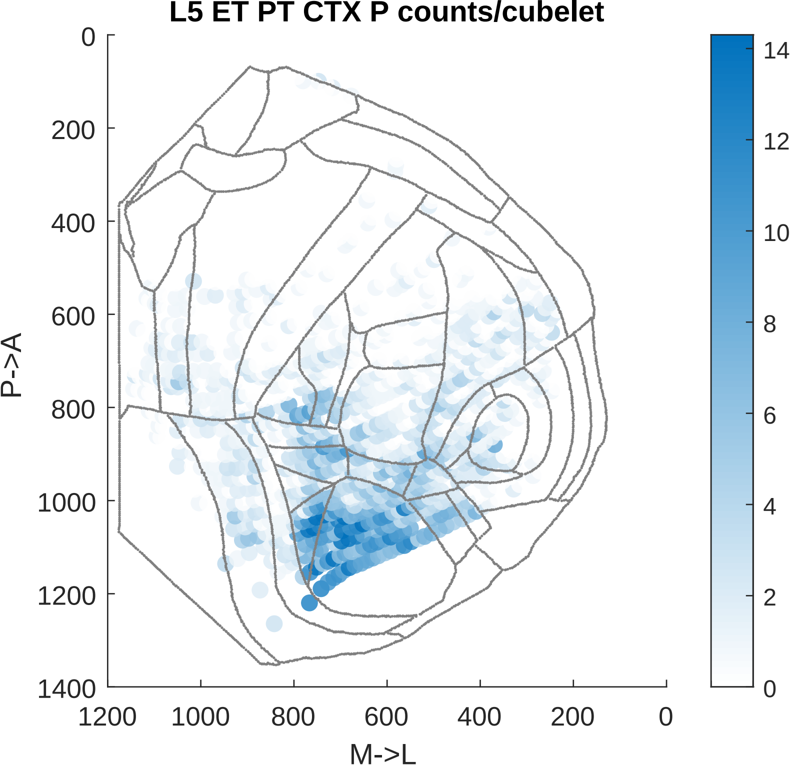

L5 ET PT CTX P counts/cubelet
P->A
M->L
0
200
400
600
800
1000
1200
1400
1200
1000
800
600
400
200
0
14
12
10
8
6
4
2
0

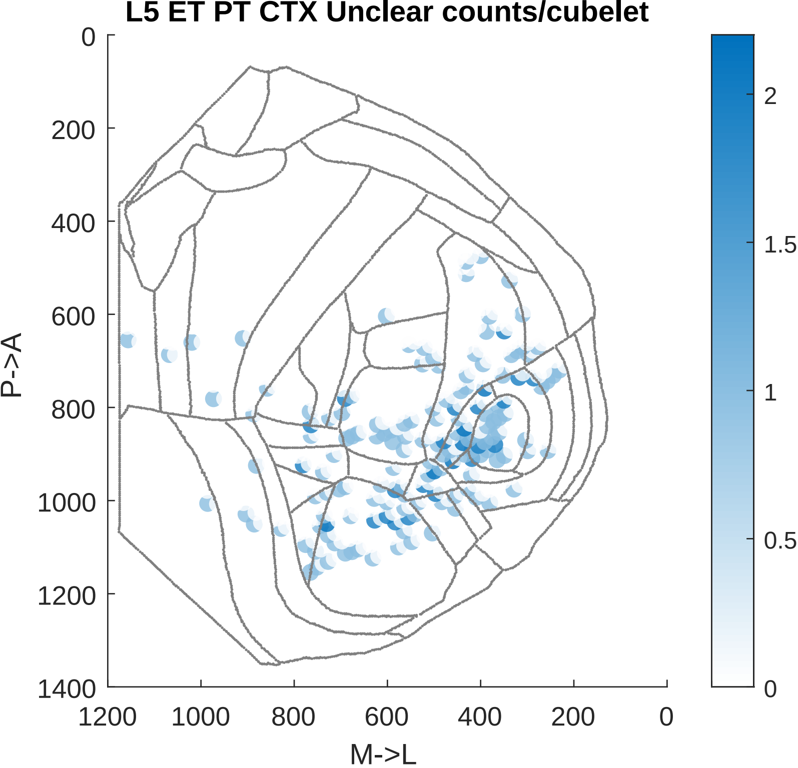

L5 ET PT CTX Unclear counts/cubelet
P->A
M->L
0
200
400
600
800
1000
1200
1400
1200
1000
800
600
400
200
0
2
1.5
1
0.5
0

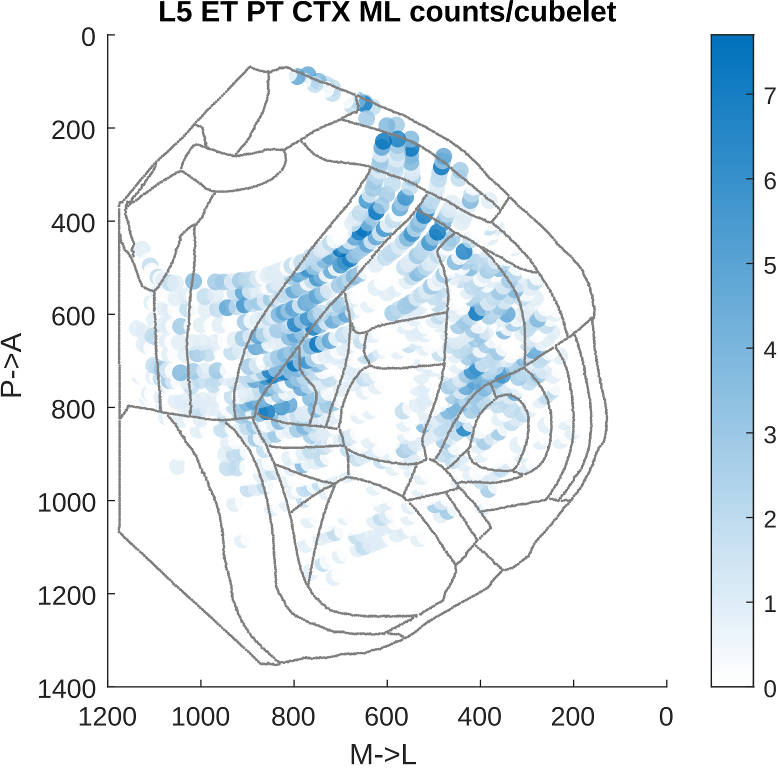

L5 ET PT CTX ML counts/cubelet
P->A
M->L
0
200
400
600
800
1000
1200
1400
1200
1000
800
600
400
200
0
7
6
5
4
3
2
1
0

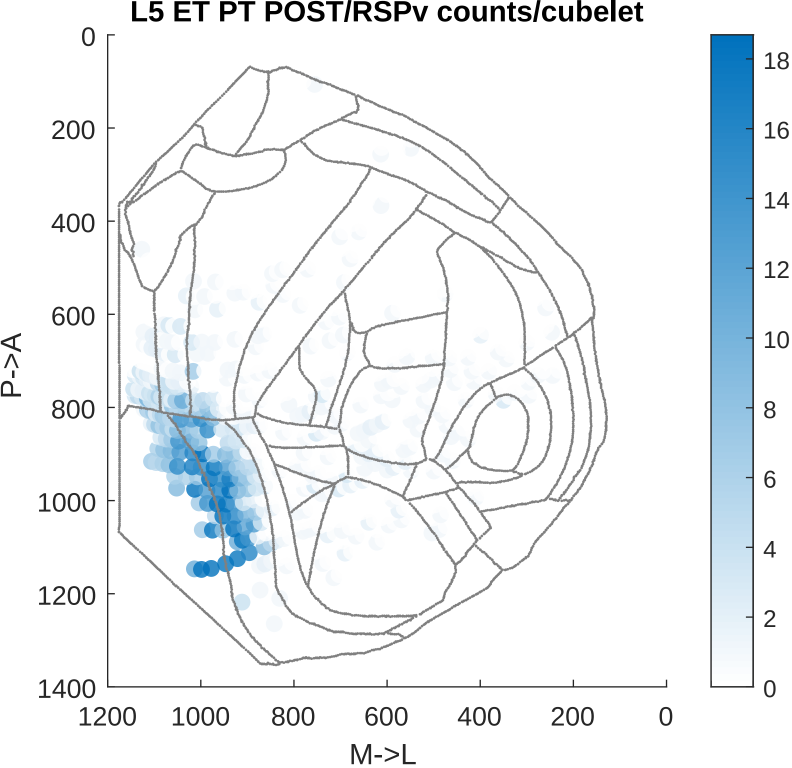

L5 ET PT POST/RSPv counts/cubelet
P->A
M->L
0
200
400
600
800
1000
1200
1400
1200
1000
800
600
400
200
0
18
16
14
12
10
8
6
4
2
0

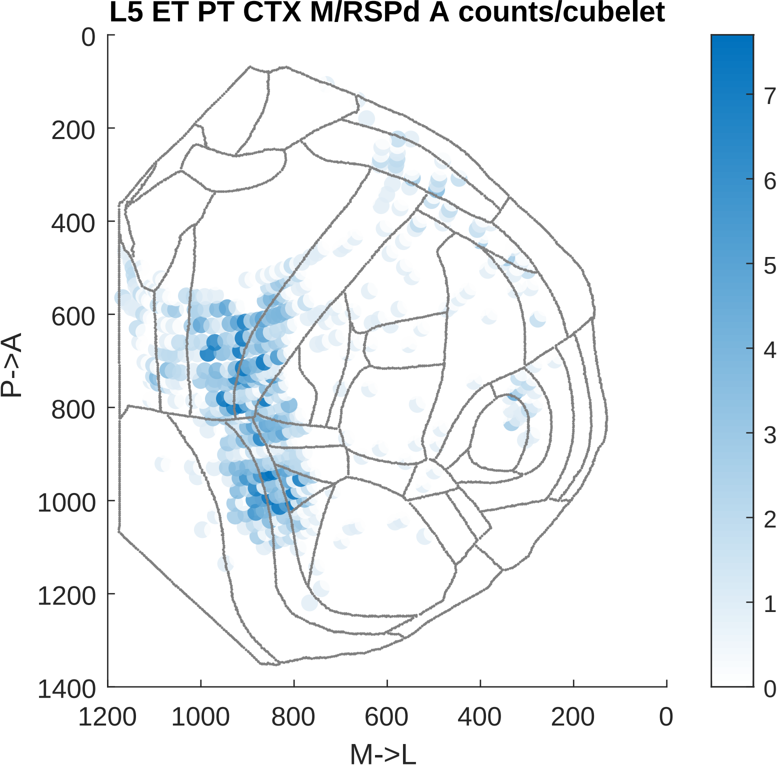
L5 ET PT CTX M/RSPd A counts/cubelet
P->A
M->L
0
200
400
600
800
1000
1200
1400
1200
1000
800
600
400
200
0
7
6
5
4
3
2
1
0

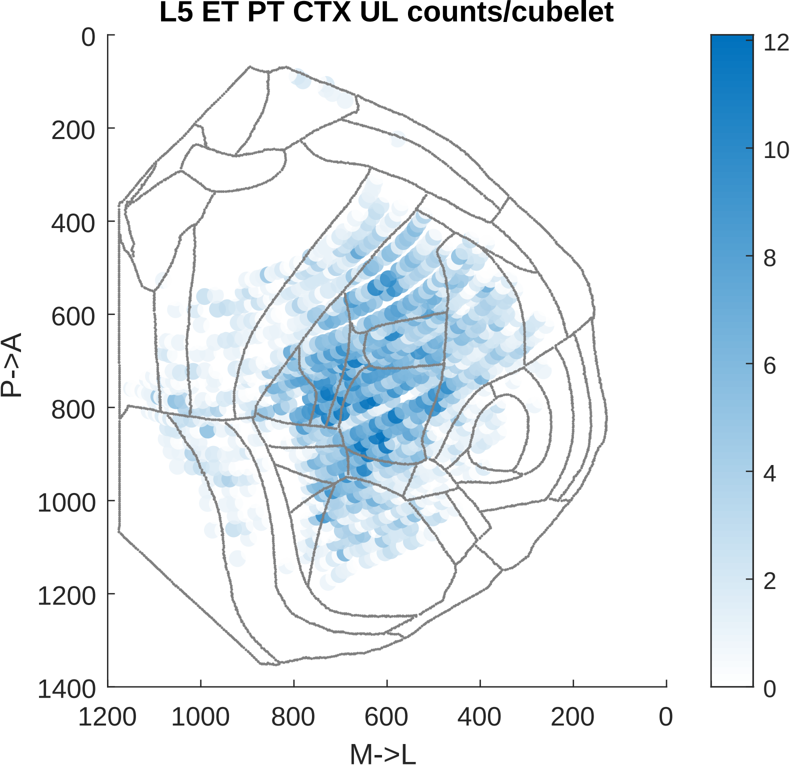
L5 ET PT CTX UL counts/cubelet
P->A
M->L
0
200
400
600
800
1000
1200
1400
1200
1000
800
600
400
200
0
12
10
8
6
4
2
0

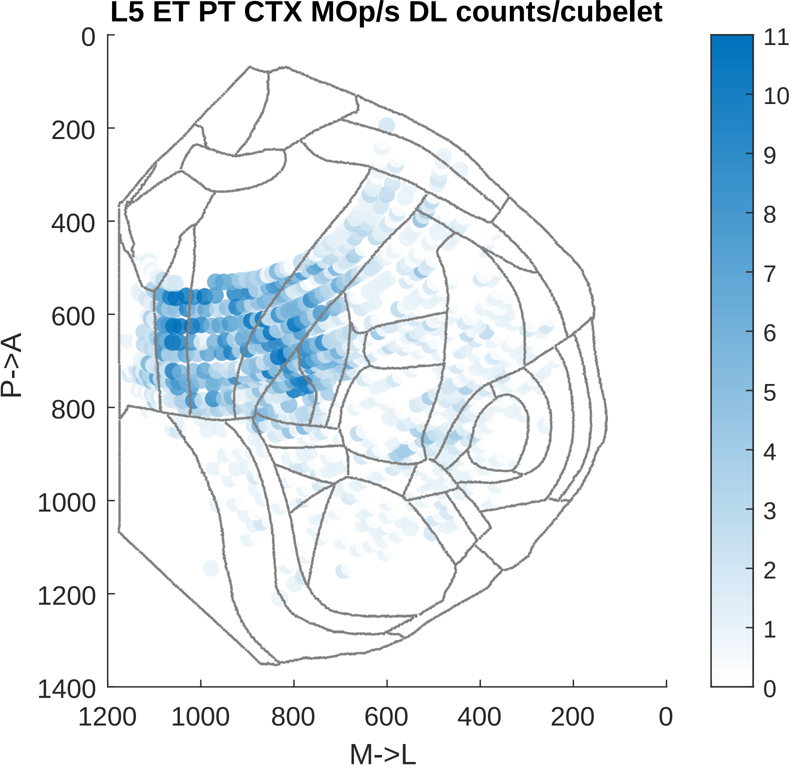
L5 ET PT CTX MOp/s DL counts/cubelet
P->A
M->L
0
200
400
600
800
1000
1200
1400
1200
1000
800
600
400
200
0
11
10
9
8
7
6
5
4
3
2
1
0

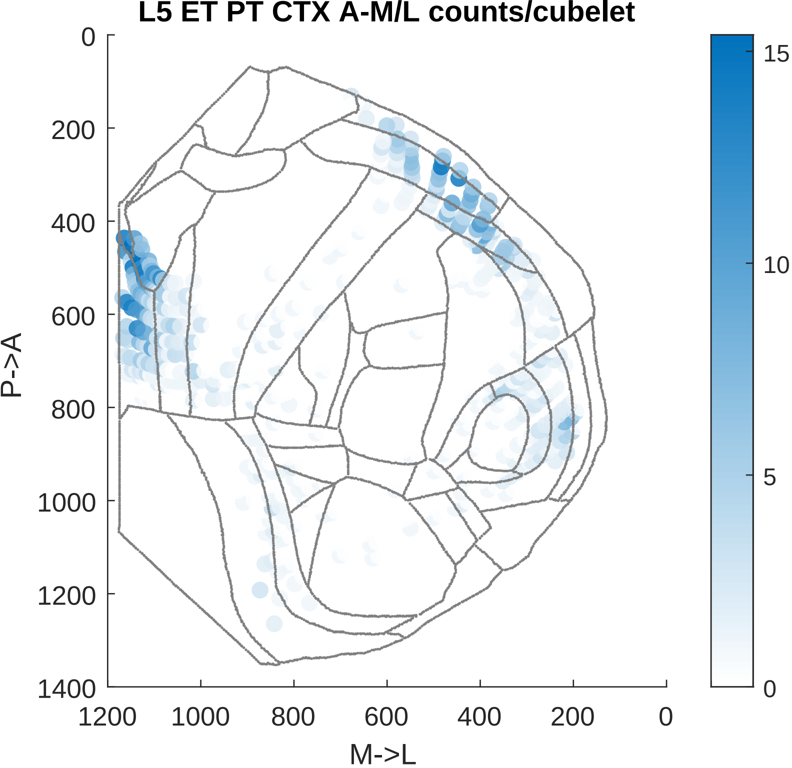

L5 ET PT CTX A-M/L counts/cubelet
0
200
400
600
800
1000
1200
1400
P->A
1200
1000
800
600
400
200
0
M->L
15
10
5
0

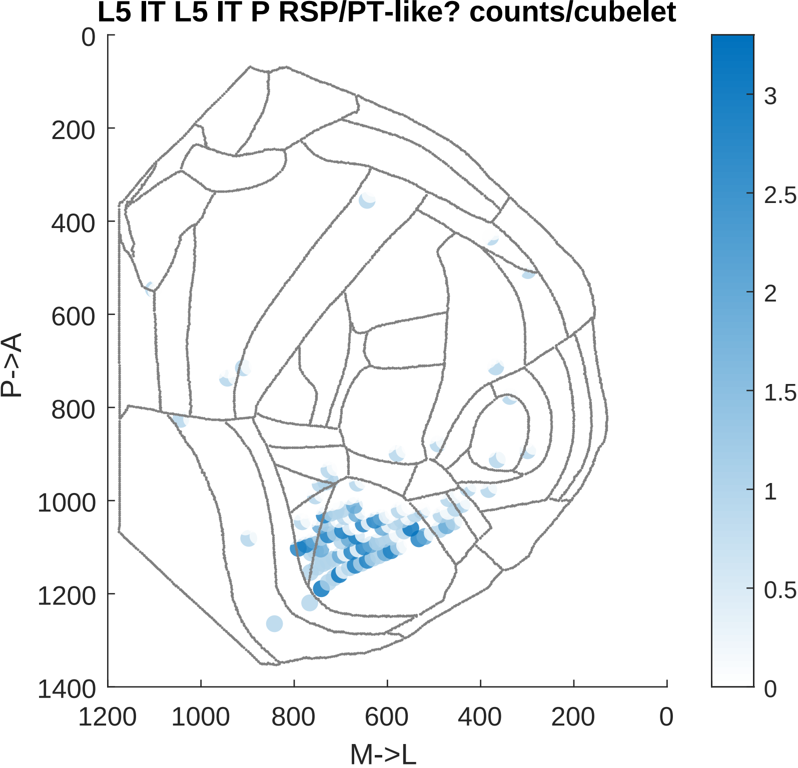

L5 IT L5 IT P RSP/PT-like? counts/cubelet
0
200
400
600
800
1000
1200
1400
P->A
1200
1000
800
600
400
200
0
M->L
3
2.5
2
1.5
1
0.5
0

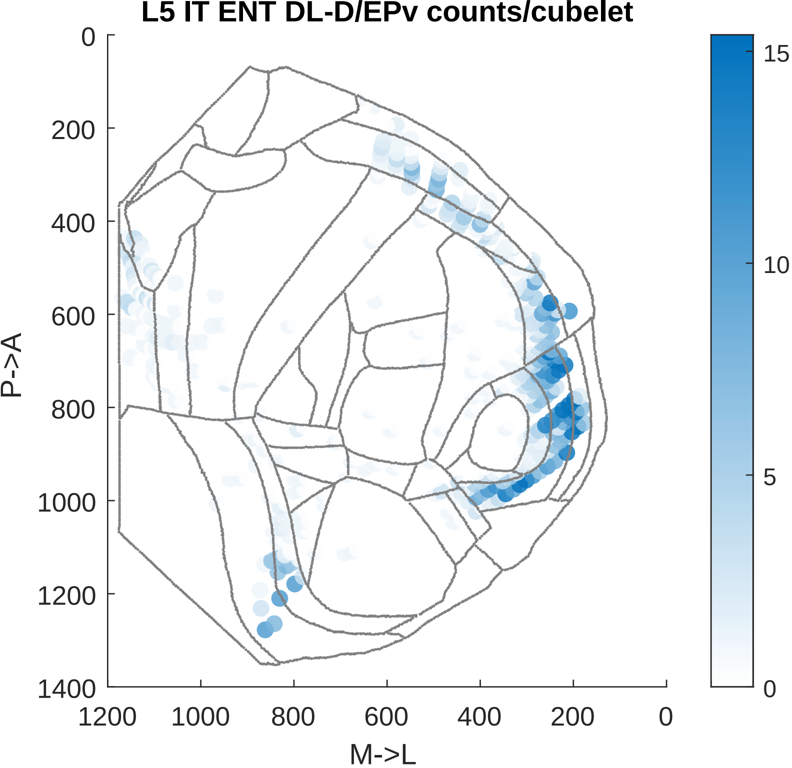

L5 IT ENT DL-D/EPv counts/cubelet
0
200
400
600
800
1000
1200
1400
P->A
1200
1000
800
600
400
200
0
M->L
15
10
5
0

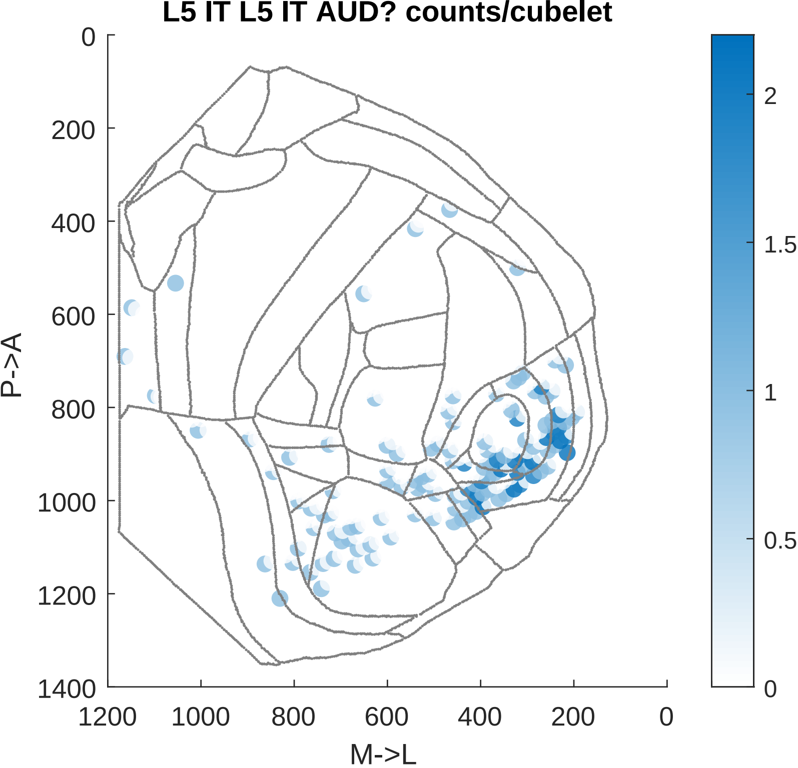

L5 IT L5 IT AUD? counts/cubelet
0
200
400
600
800
1000
1200
1400
P->A
1200
1000
800
600
400
200
0
M->L
2
1.5
1
0.5
0

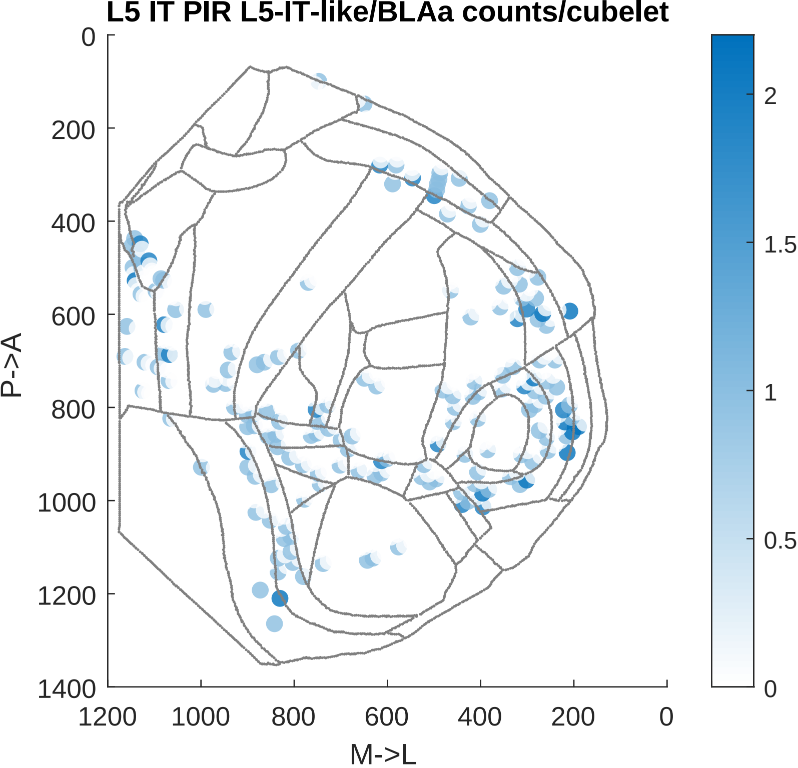

L5 IT PIR L5-IT-like/BLAa counts/cubelet
P->A
M->L
0
200
400
600
800
1000
1200
1400
1200
1000
800
600
400
200
0
2
1.5
1
0.5
0

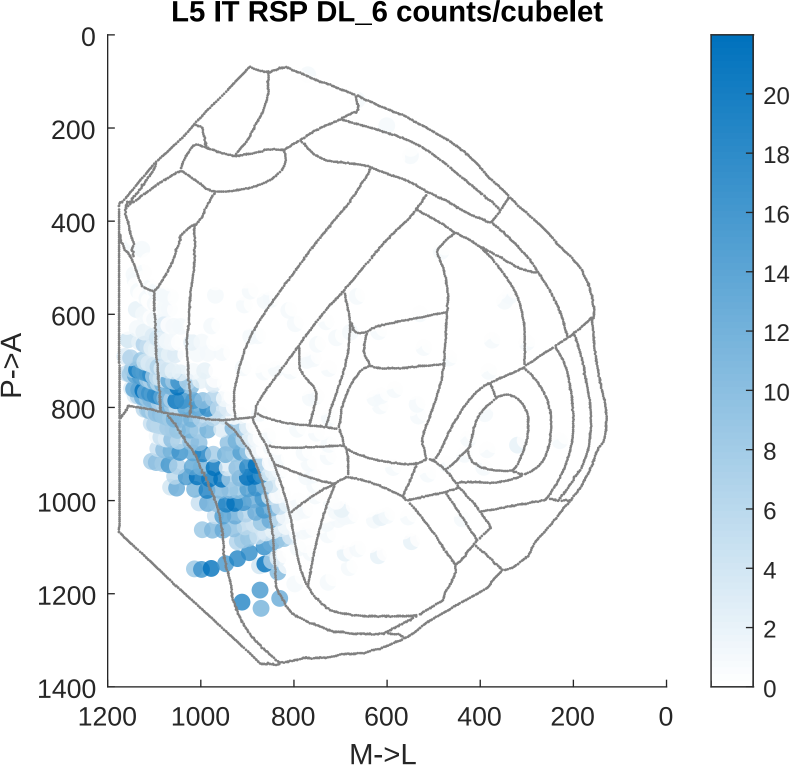

L5 IT RSP DL_6 counts/cubelet
P->A
M->L
0
200
400
600
800
1000
1200
1400
1200
1000
800
600
400
200
0
20
18
16
14
12
10
8
6
4
2
0

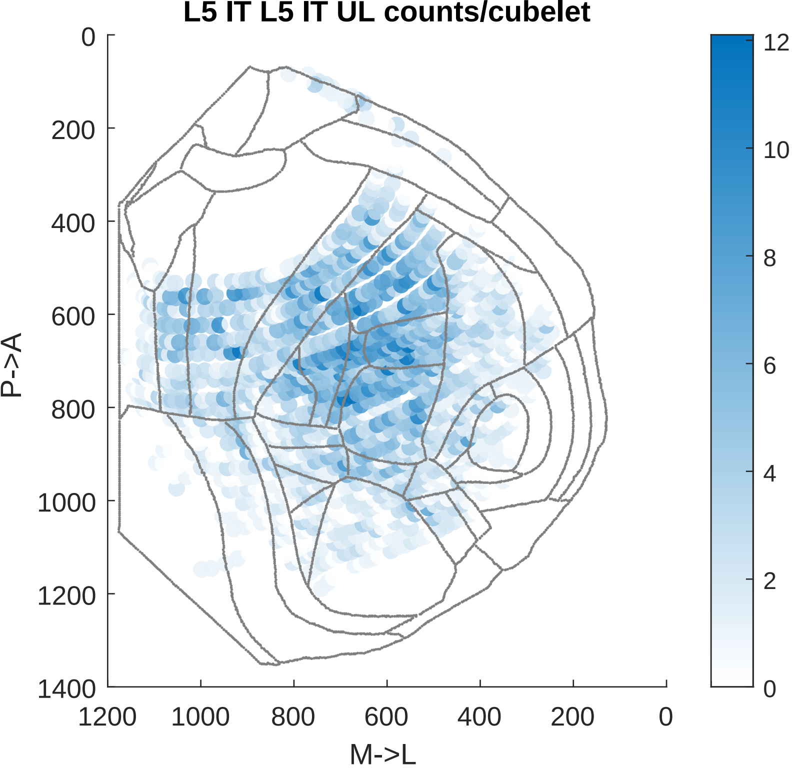

L5 IT L5 IT UL counts/cubelet
P->A
M->L
0
200
400
600
800
1000
1200
1400
1200
1000
800
600
400
200
0
12
10
8
6
4
2
0

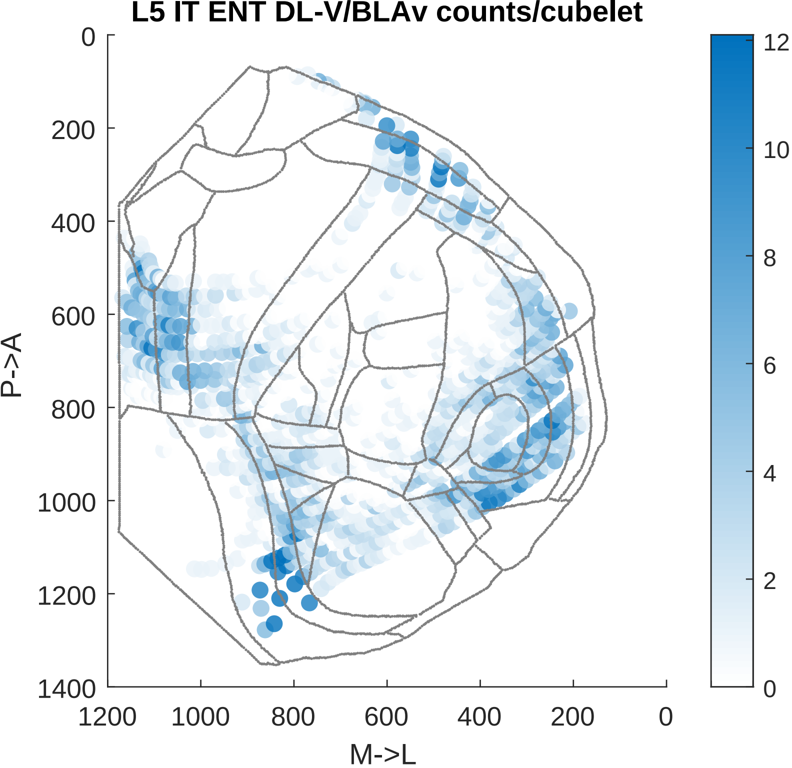

L5 IT ENT DL-V/BLAv counts/cubelet
P->A
M->L
0
200
400
600
800
1000
1200
1400
1200
1000
800
600
400
200
0
12
10
8
6
4
2
0

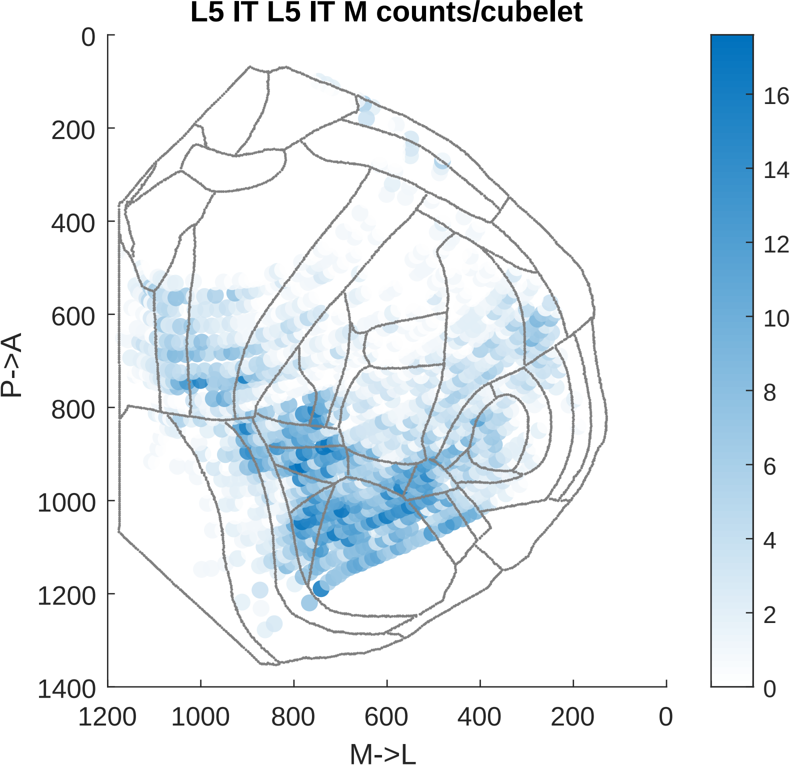

L5 IT L5 IT M counts/cubelet
0
200
400
600
800
1000
1200
1400
P->A
1200
1000
800
600
400
200
0
M->L
16
14
12
10
8
6
4
2
0

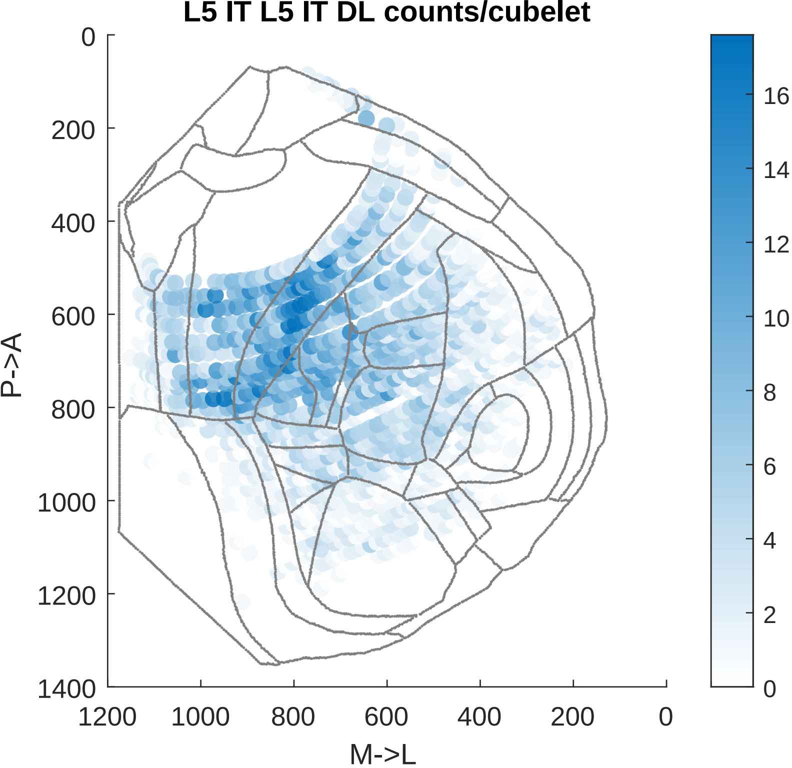
L5 IT L5 IT DL counts/cubelet
0
200
400
600
800
1000
1200
1400
P->A
1200
1000
800
600
400
200
0
M->L
16
14
12
10
8
6
4
2
0

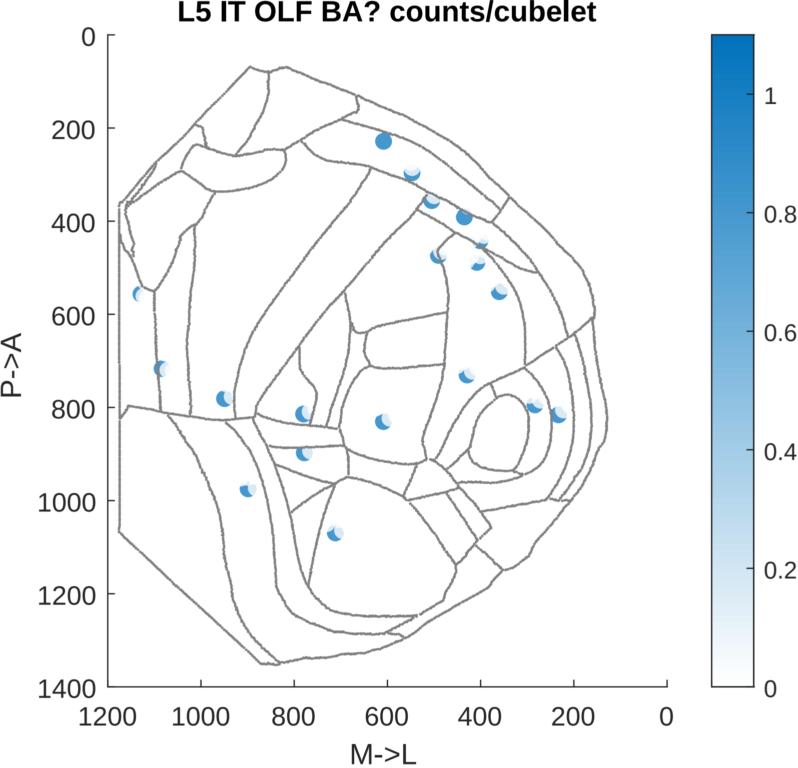

L5 IT OLF BA? counts/cubelet
0
200
400
600
800
1000
1200
1400
P->A
1200
1000
800
600
400
200
0
M->L
1
0.8
0.6
0.4
0.2
0

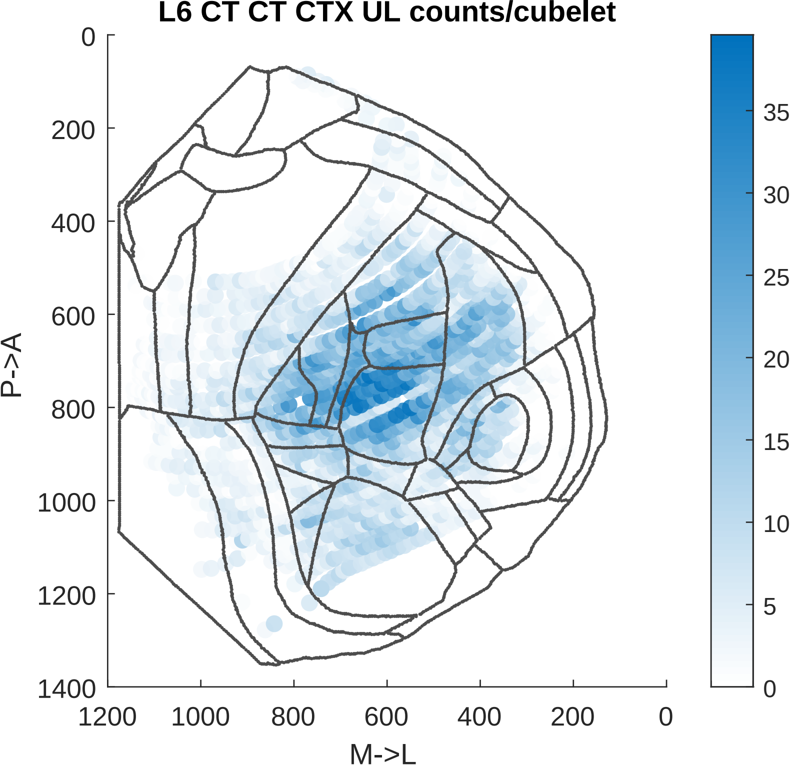
L6 CT CT CTX UL counts/cubelet
0
200
400
600
800
1000
1200
1400
P->A
1200
1000
800
600
400
200
0
M->L
35
30
25
20
15
10
5
0

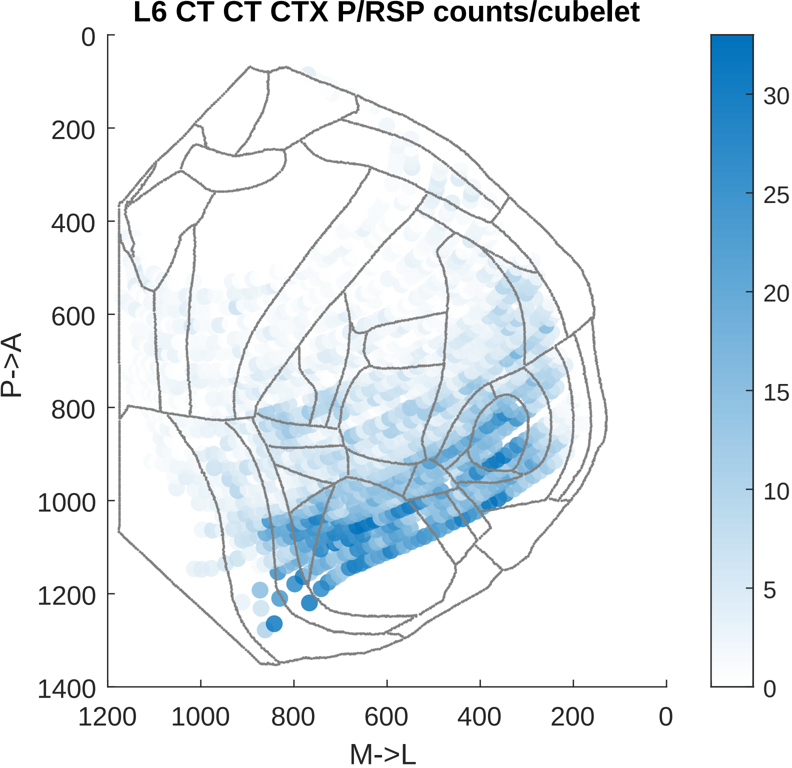

L6 CT CT CTX P/RSP counts/cubelet
0
200
400
600
800
1000
1200
1400
P->A
1200
1000
800
600
400
200
0
M->L
30
25
20
15
10
5
0

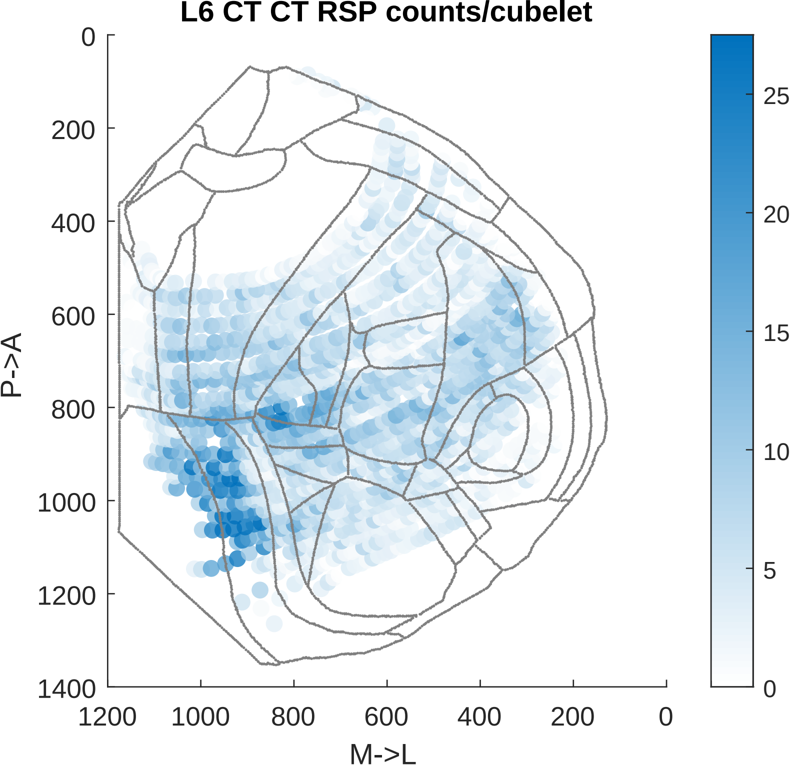

L6 CT CT RSP counts/cubelet
P->A
M->L
0
200
400
600
800
1000
1200
1400
1200
1000
800
600
400
200
0
25
20
15
10
5
0

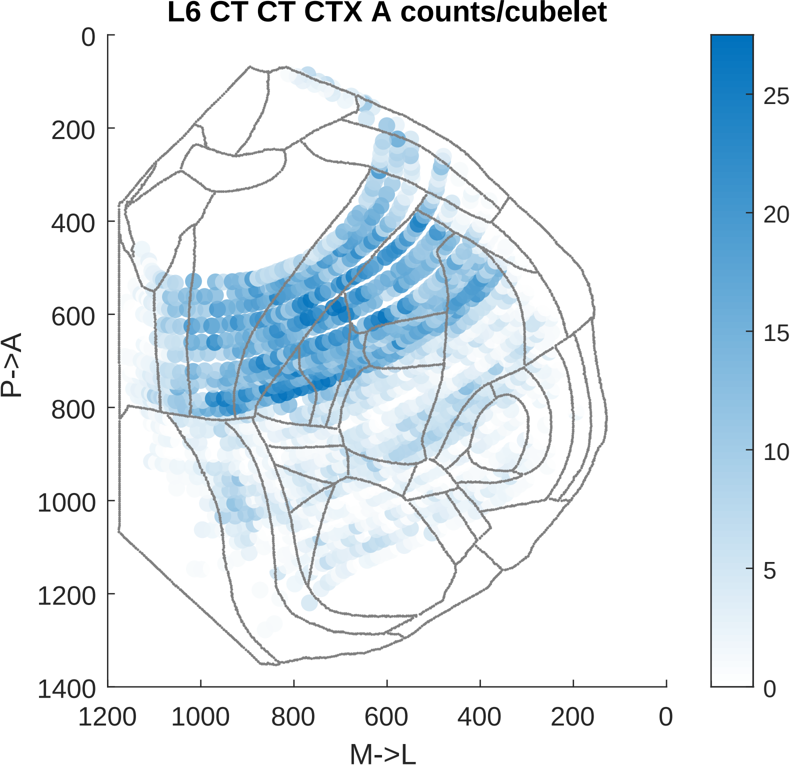
L6 CT CT CTX A counts/cubelet
P->A
M->L
0
200
400
600
800
1000
1200
1400
1200
1000
800
600
400
200
0
25
20
15
10
5
0

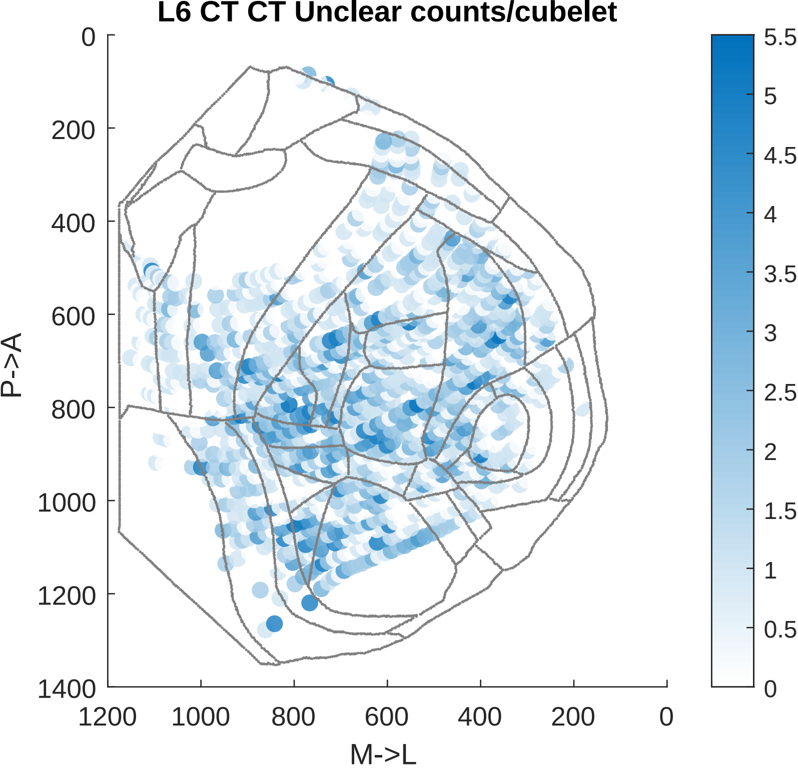

L6 CT CT Unclear counts/cubelet
P->A
M->L
0
200
400
600
800
1000
1200
1400
1200
1000
800
600
400
200
0
5.5
5
4.5
4
3.5
3
2.5
2
1.5
1
0.5
0

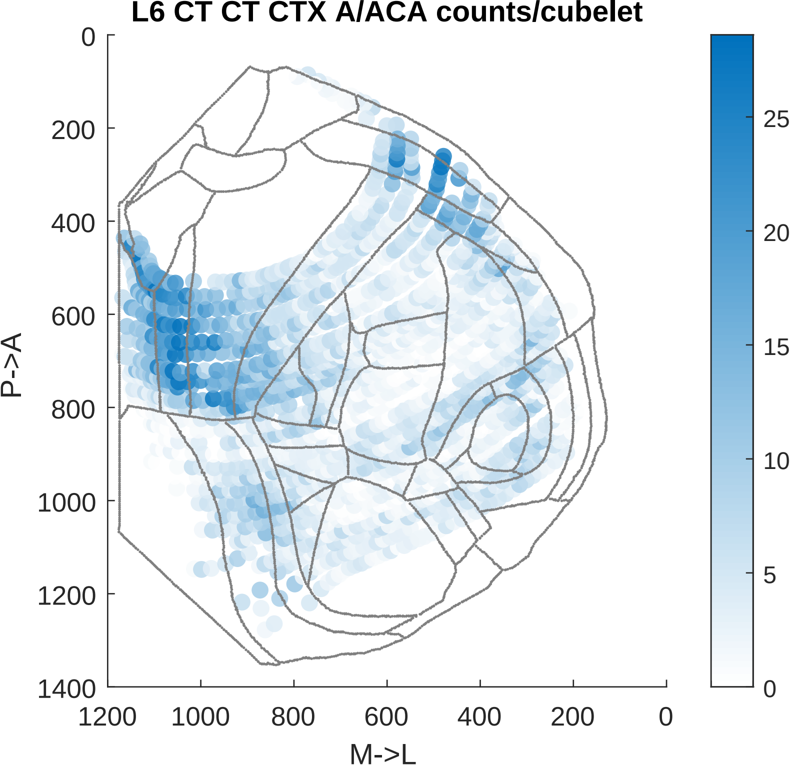

L6 CT CT CTX A/ACA counts/cubelet
P->A
M->L
0
200
400
600
800
1000
1200
1400
1200
1000
800
600
400
200
0
25
20
15
10
5
0

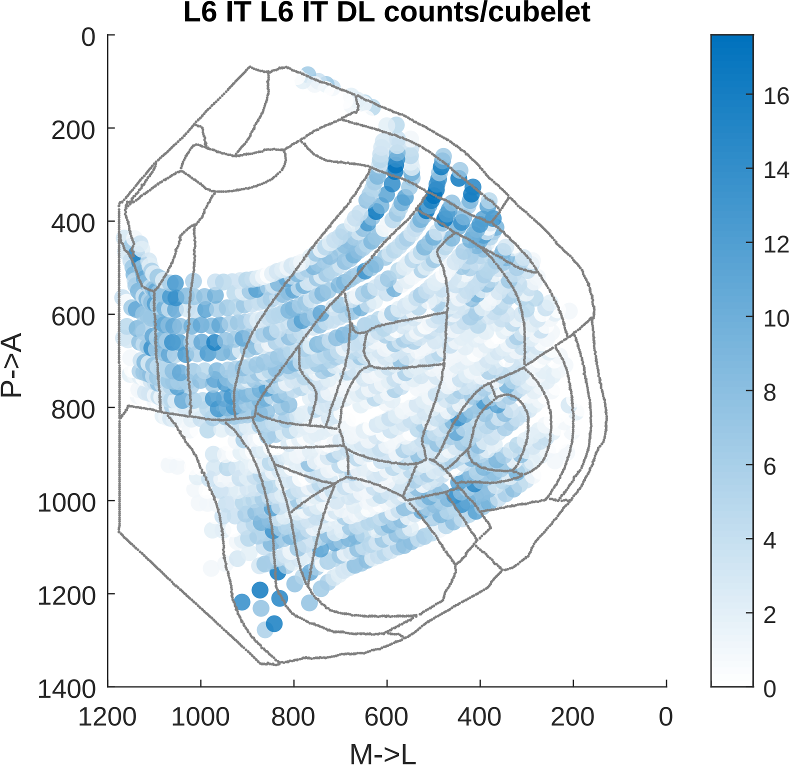

L6 IT L6 IT DL counts/cubelet
P->A
M->L
0
200
400
600
800
1000
1200
1400
1200
1000
800
600
400
200
0
16
14
12
10
8
6
4
2
0

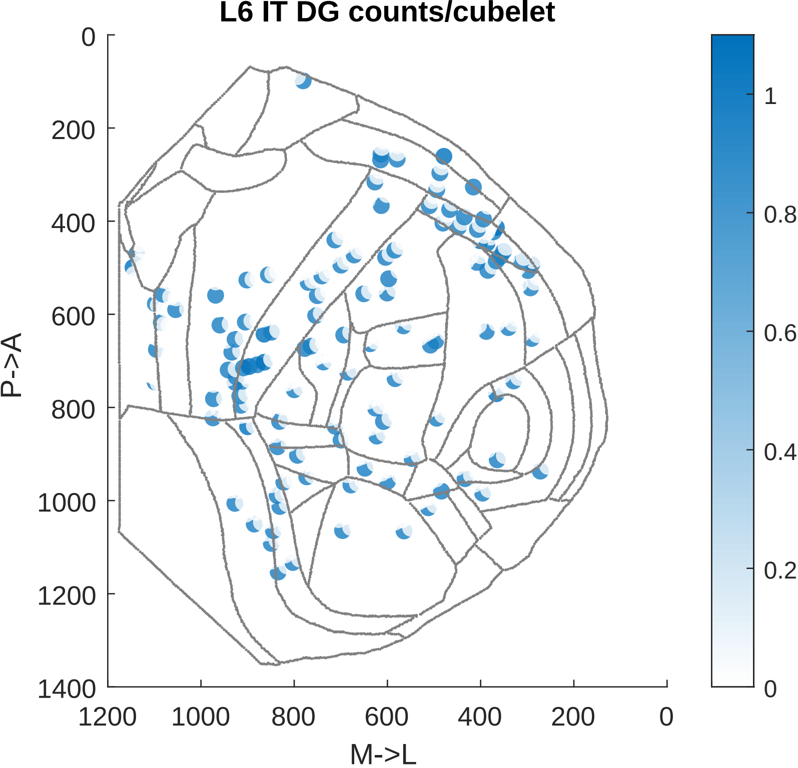

L6 IT DG counts/cubelet
0
200
400
600
800
1000
1200
1400
P->A
1200
1000
800
600
400
200
0
M->L
1
0.8
0.6
0.4
0.2
0

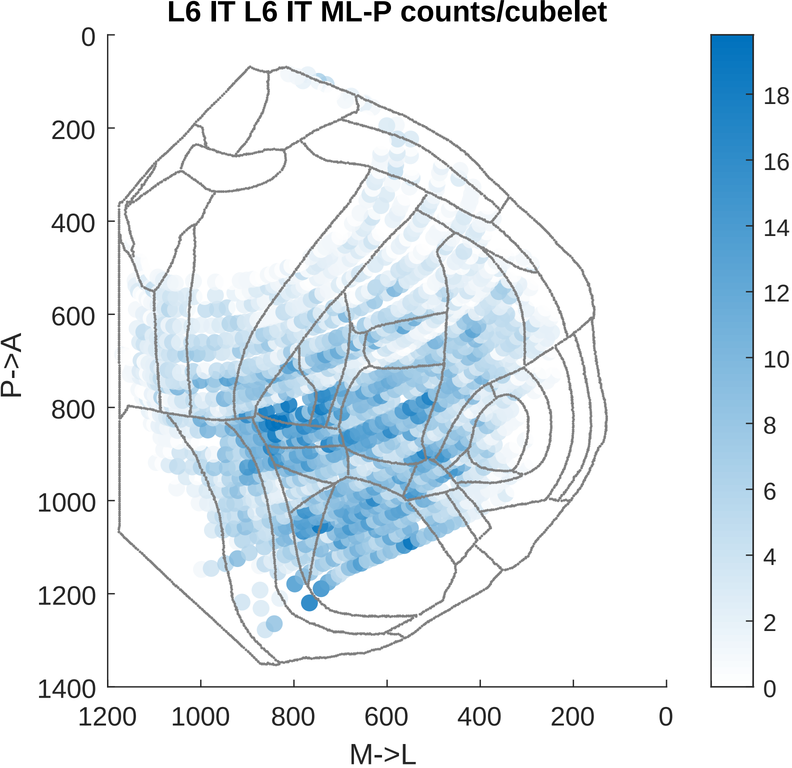
L6 IT L6 IT ML-P counts/cubelet
P->A
M->L
0
200
400
600
800
1000
1200
1400
1200
1000
800
600
400
200
0
18
16
14
12
10
8
6
4
2
0

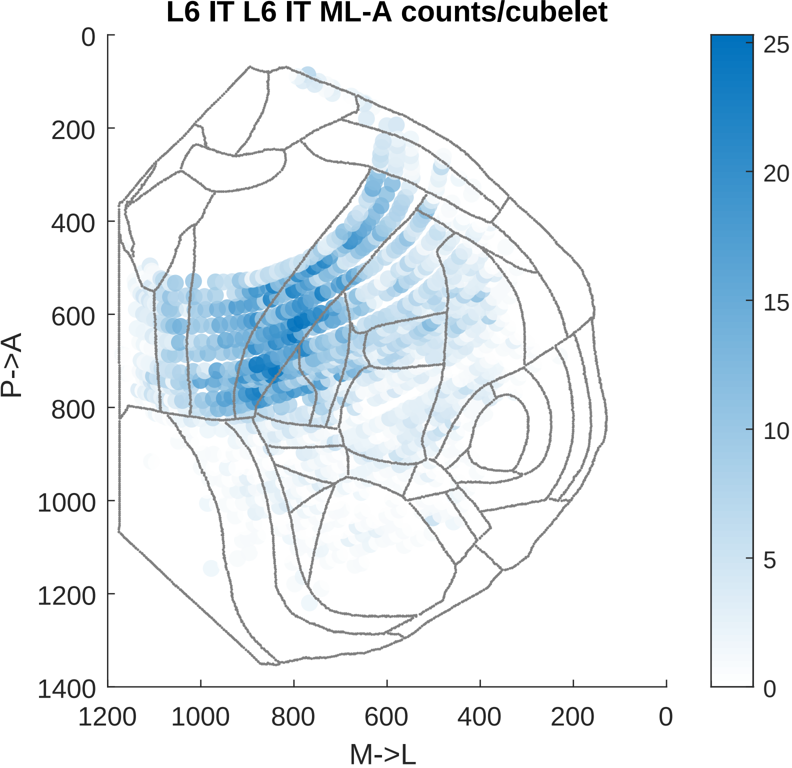

L6 IT L6 IT ML-A counts/cubelet
0
200
400
600
800
1000
1200
1400
P->A
1200
1000
800
600
400
200
0
M->L
25
20
15
10
5
0

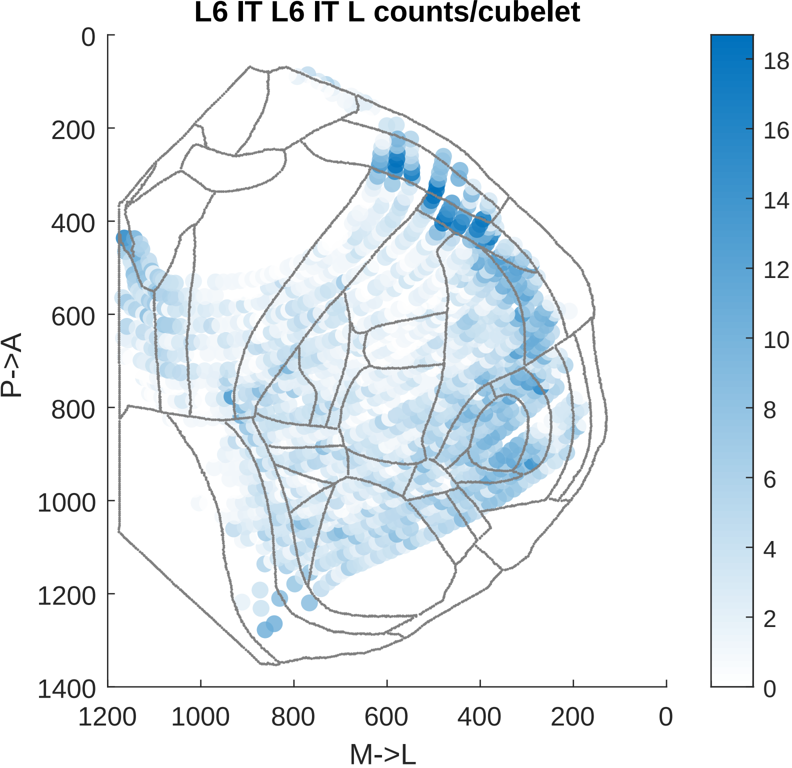
L6 IT L6 IT L counts/cubelet
P->A
M->L
0
200
400
600
800
1000
1200
1400
1200
1000
800
600
400
200
0
18
16
14
12
10
8
6
4
2
0

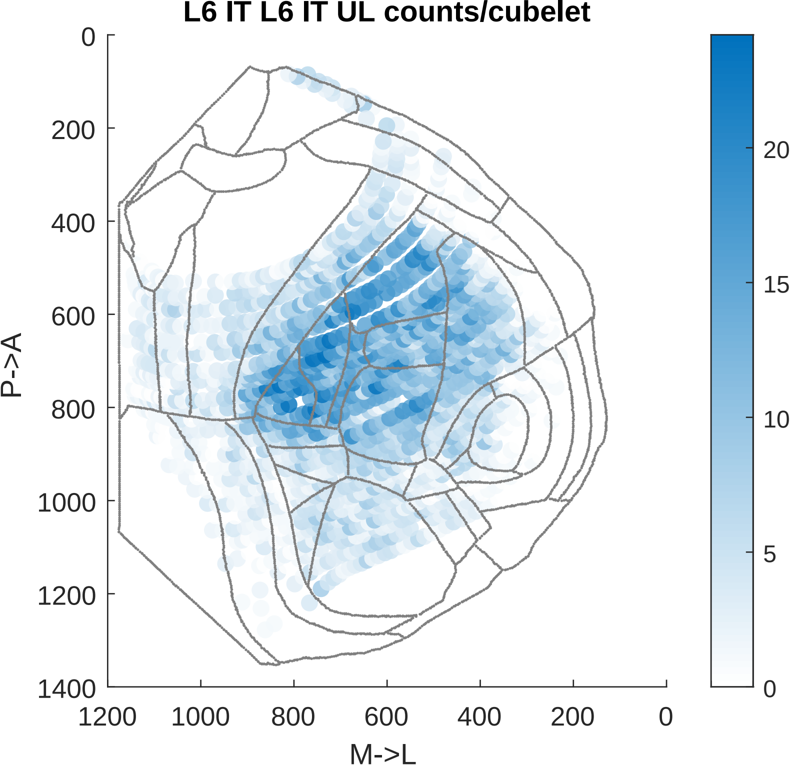

L6 IT L6 IT UL counts/cubelet
P->A
M->L
0
200
400
600
800
1000
1200
1400
1200
1000
800
600
400
200
0
20
15
10
5
0

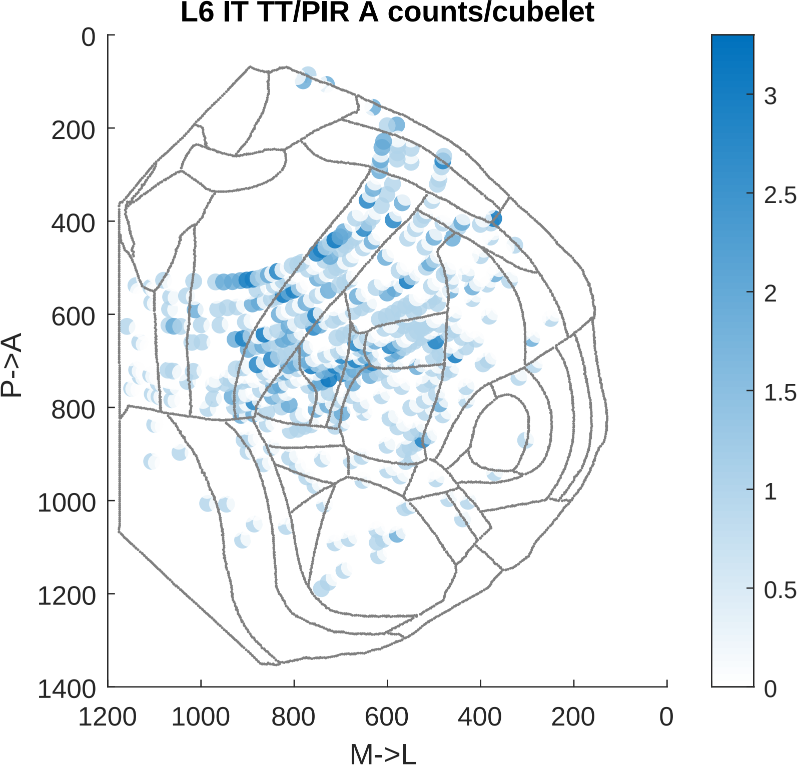
L6 IT TT/PIR A counts/cubelet
P->A
M->L
0
200
400
600
800
1000
1200
1400
1200
1000
800
600
400
200
0
3
2.5
2
1.5
1
0.5
0

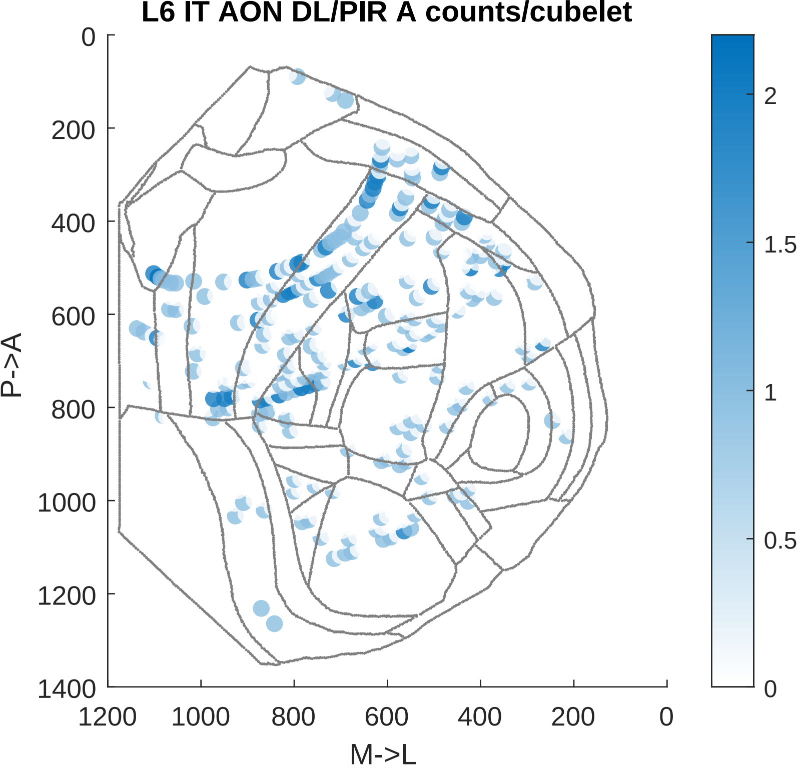

L6 IT AON DL/PIR A counts/cubelet
P->A
M->L
0
200
400
600
800
1000
1200
1400
1200
1000
800
600
400
200
0
2
1.5
1
0.5
0

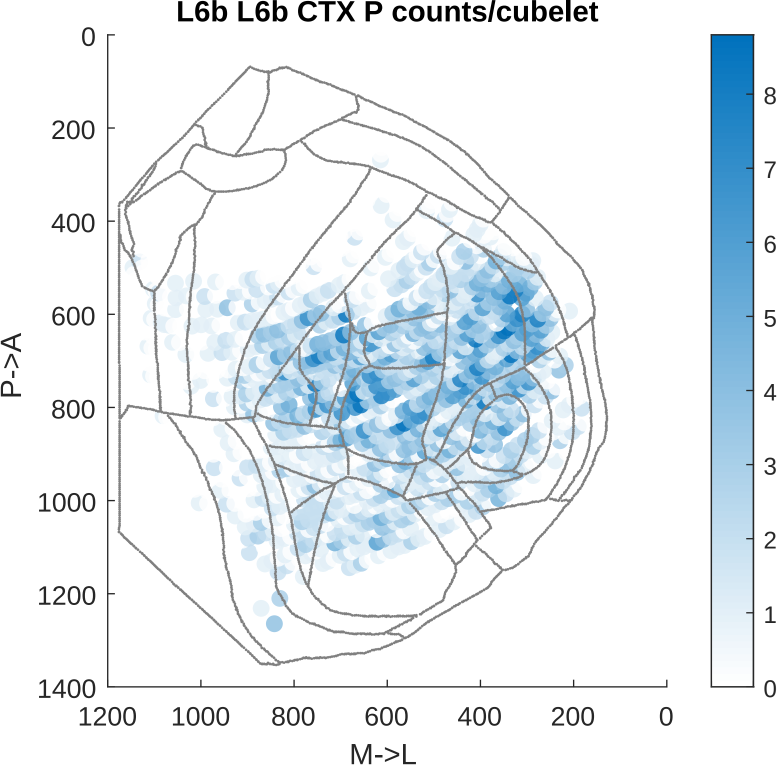

L6b L6b CTX P counts/cubelet
P->A
M->L
0
200
400
600
800
1000
1200
1400
1200
1000
800
600
400
200
0
8
7
6
5
4
3
2
1
0

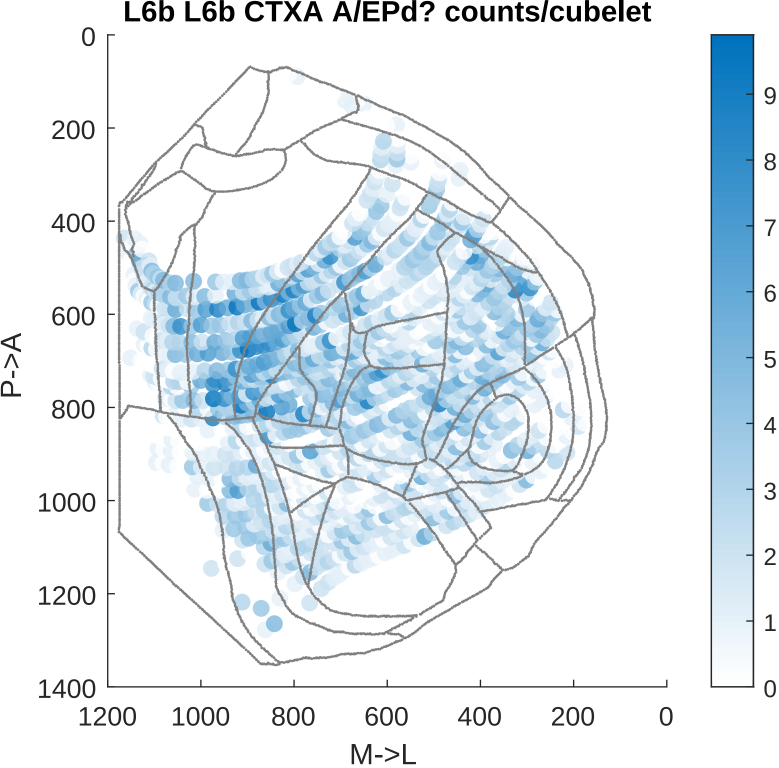

L6b L6b CTXA A/EPd? counts/cubelet
P->A
M->L
0
200
400
600
800
1000
1200
1400
1200
1000
800
600
400
200
0
9
8
7
6
5
4
3
2
1
0

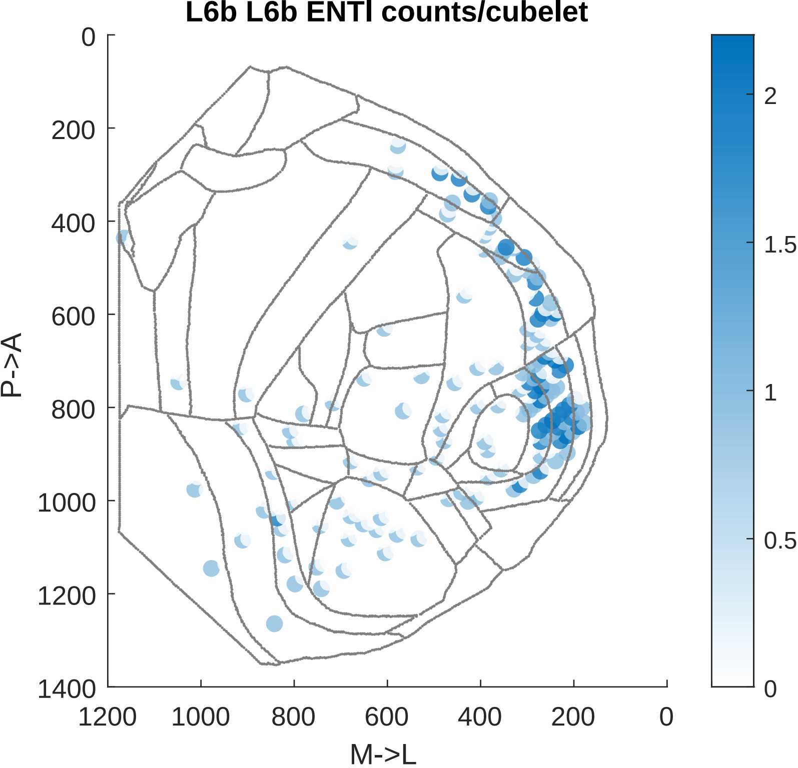

L6b L6b ENTl counts/cubelet
P->A
M->L
0
200
400
600
800
1000
1200
1400
1200
1000
800
600
400
200
0
2
1.5
1
0.5
0

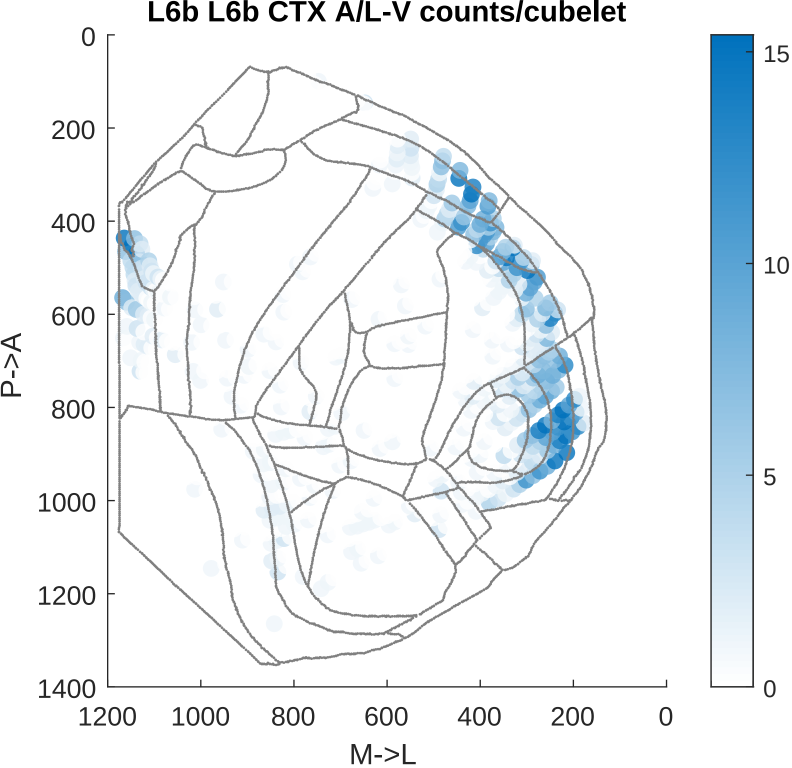

L6b L6b CTX A/L-V counts/cubelet
P->A
M->L
0
200
400
600
800
1000
1200
1400
1200
1000
800
600
400
200
0
15
10
5
0

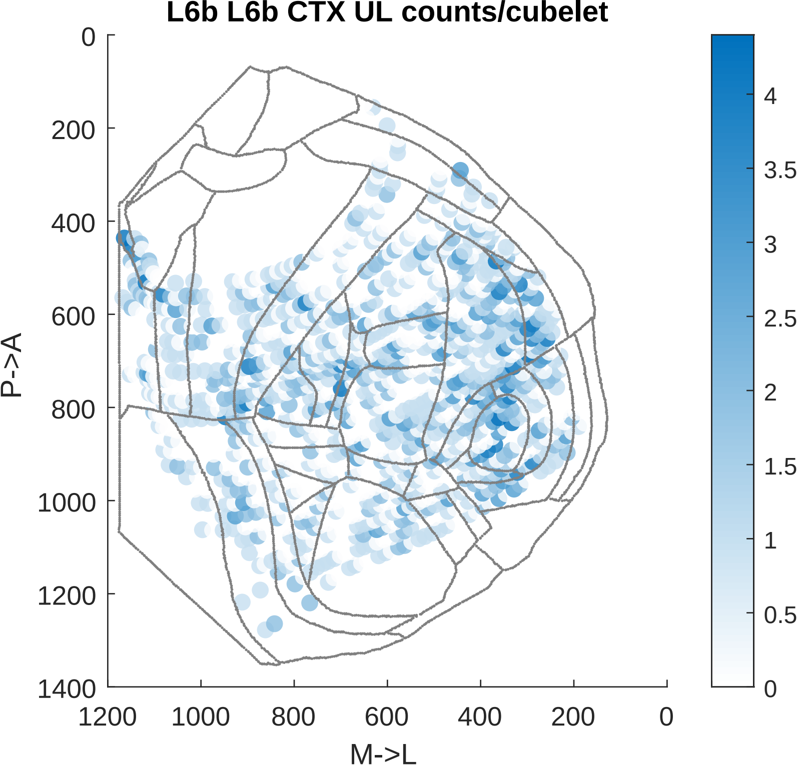

L6b L6b CTX UL counts/cubelet
P->A
M->L
0
200
400
600
800
1000
1200
1400
1200
1000
800
600
400
200
0
4
3.5
3
2.5
2
1.5
1
0.5
0

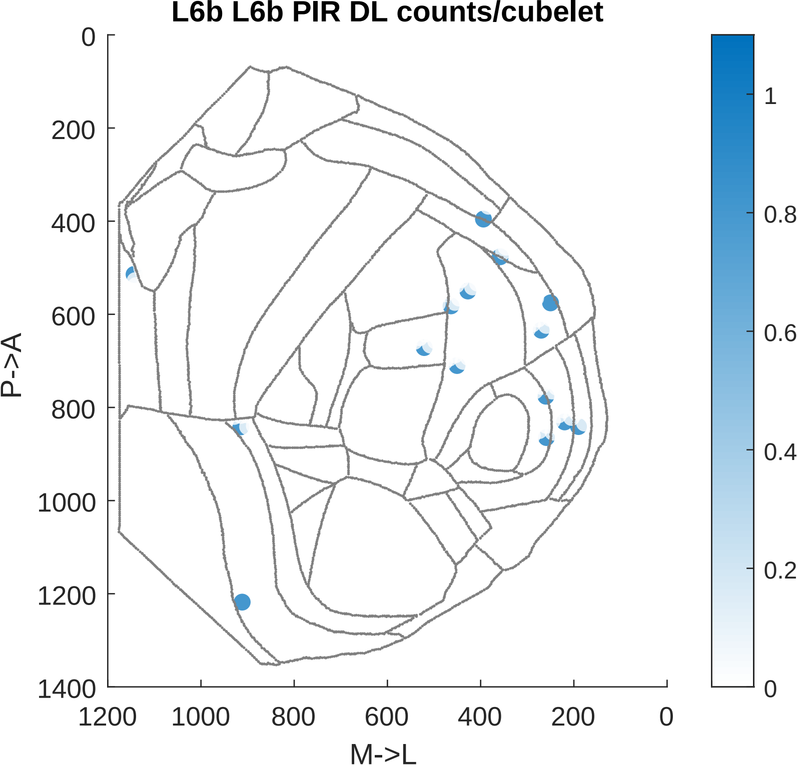

L6b L6b PIR DL counts/cubelet
0
200
400
600
800
1000
1200
1400
P->A
1200
1000
800
600
400
200
0
M->L
1
0.8
0.6
0.4
0.2
0

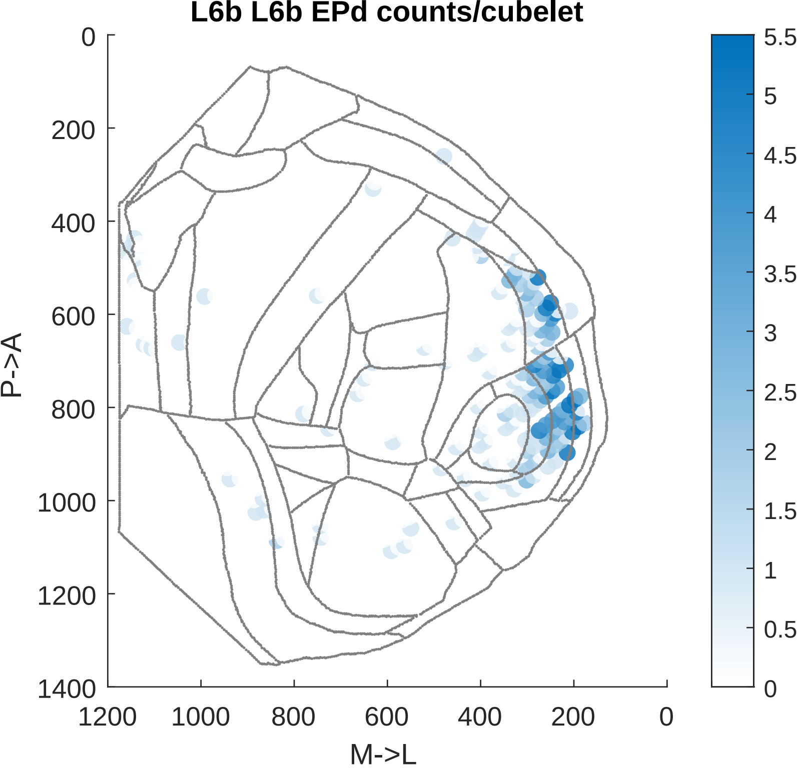
L6b L6b EPd counts/cubelet
P->A
M->L
0
200
400
600
800
1000
1200
1400
1200
1000
800
600
400
200
0
5.5
5
4.5
4
3.5
3
2.5
2
1.5
1
0.5
0

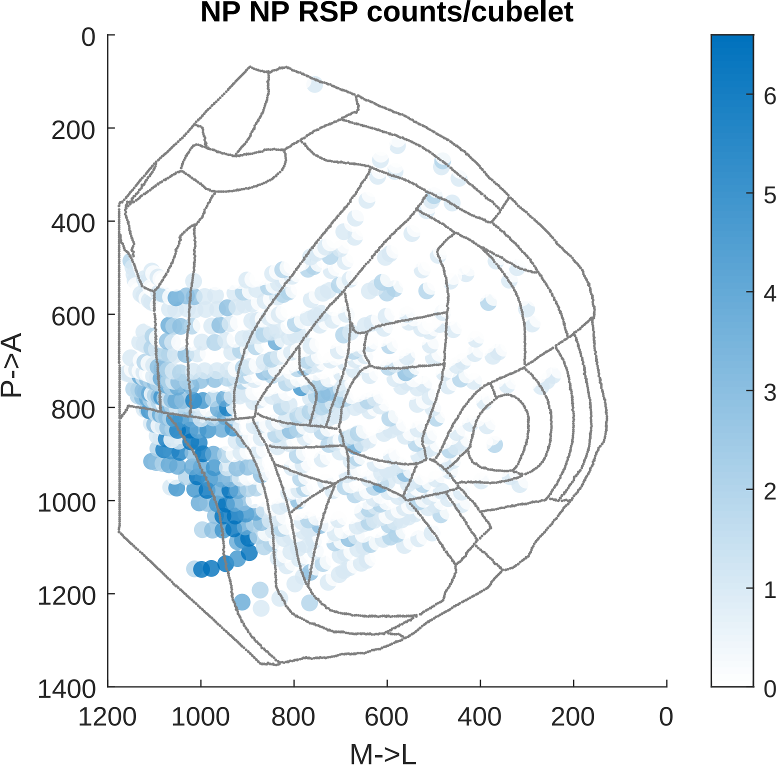

NP NP RSP counts/cubelet
0
200
400
600
800
1000
1200
1400
P->A
1200
1000
800
600
400
200
0
M->L
0
1
2
3
4
5
6

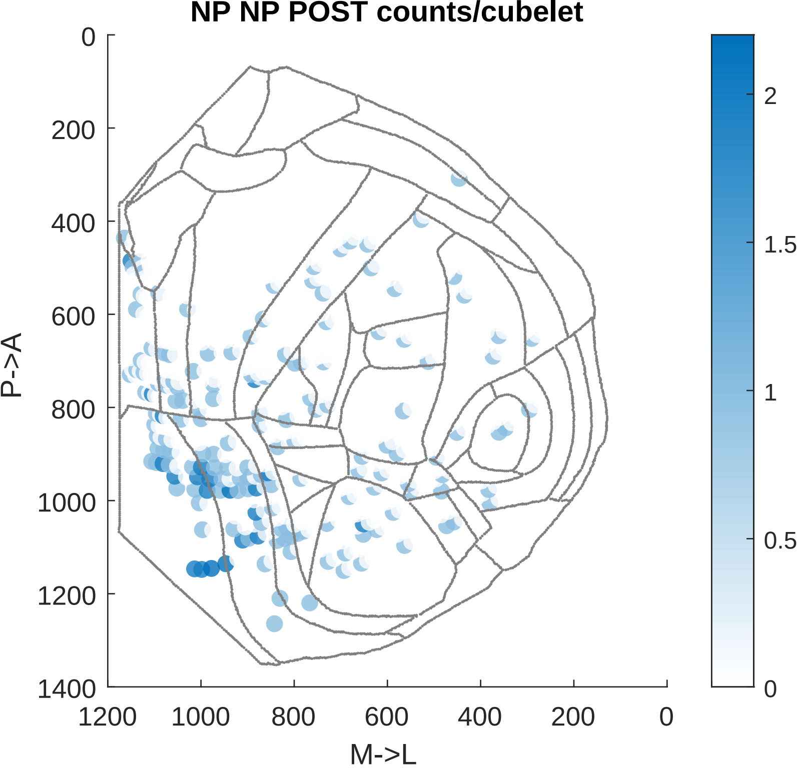

NP NP POST counts/cubelet
P->A
M->L
0
200
400
600
800
1000
1200
1400
1200
1000
800
600
400
200
0
2
1.5
1
0.5
0

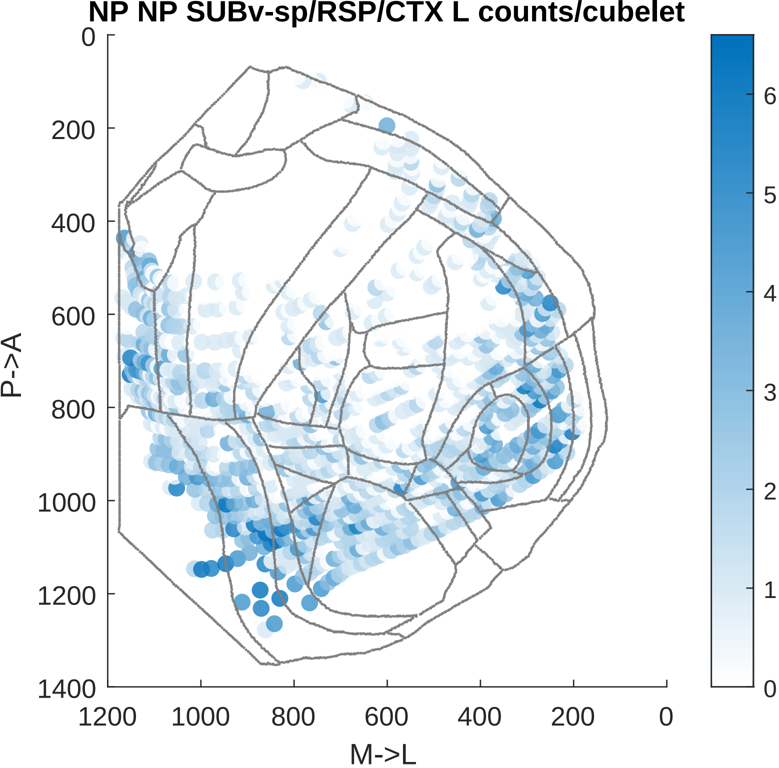
NP NP SUBv-sp/RSP/CTX L counts/cubelet
0
200
400
600
800
1000
1200
1400
P->A
1200
1000
800
600
400
200
0
M->L
6
5
4
3
2
1
0

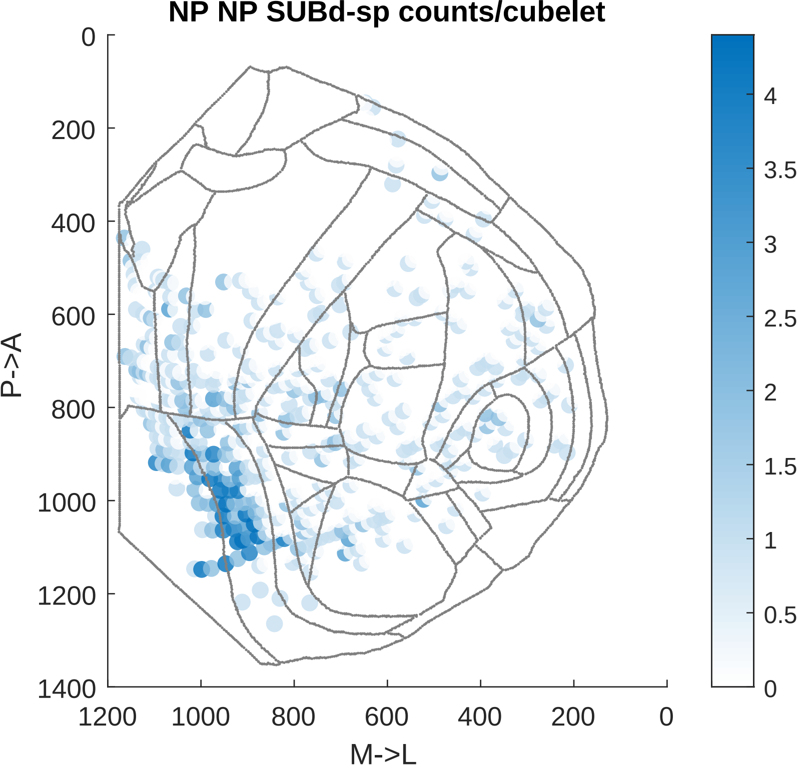
NP NP SUBd-sp counts/cubelet
0
200
400
600
800
1000
1200
1400
P->A
1200
1000
800
600
400
200
0
M->L
4
3.5
3
2.5
2
1.5
1
0.5
0

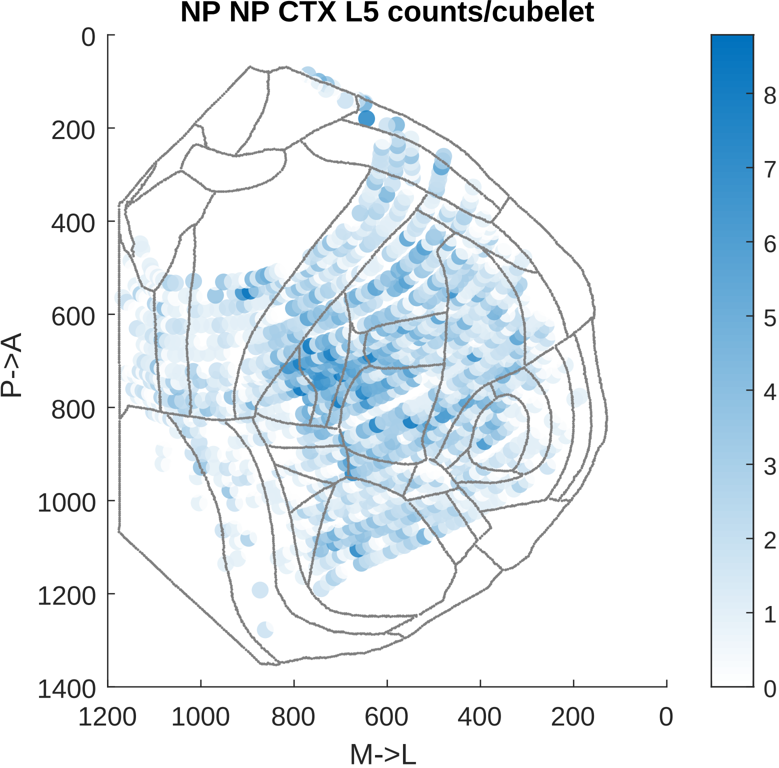

NP NP CTX L5 counts/cubelet
P->A
M->L
0
200
400
600
800
1000
1200
1400
1200
1000
800
600
400
200
0
8
7
6
5
4
3
2
1
0

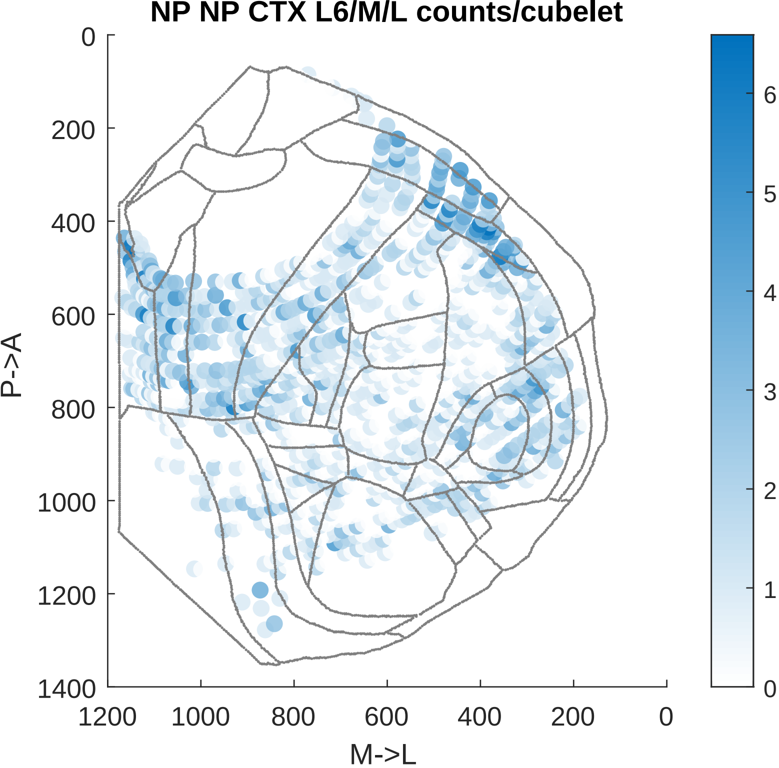

NP NP CTX L6/M/L counts/cubelet
0
200
400
600
800
1000
1200
1400
P->A
1200
1000
800
600
400
200
0
M->L
6
5
4
3
2
1
0

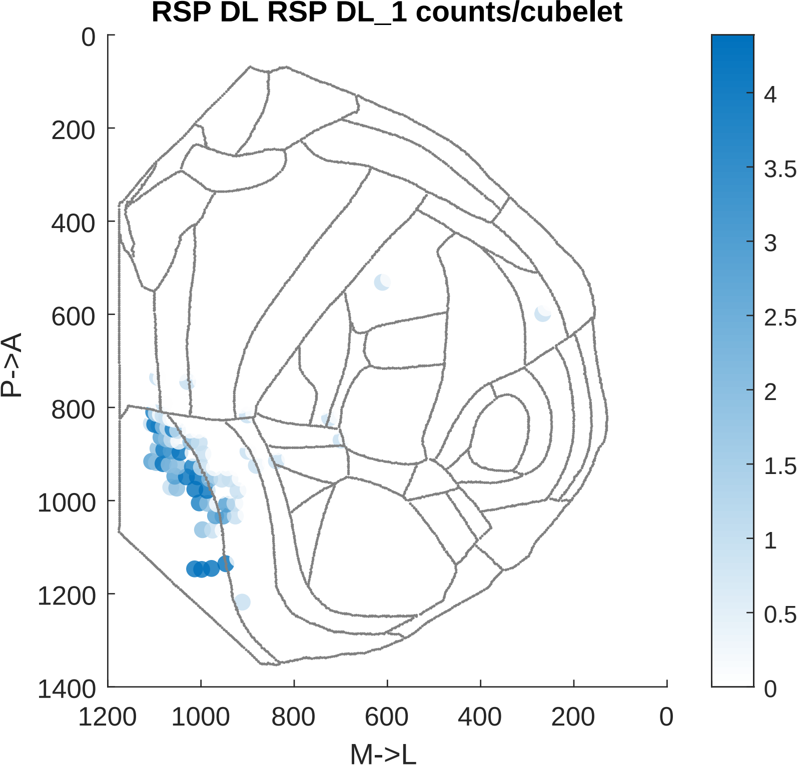

RSP DL RSP DL_1 counts/cubelet
P->A
M->L
0
200
400
600
800
1000
1200
1400
1200
1000
800
600
400
200
0
4
3.5
3
2.5
2
1.5
1
0.5
0

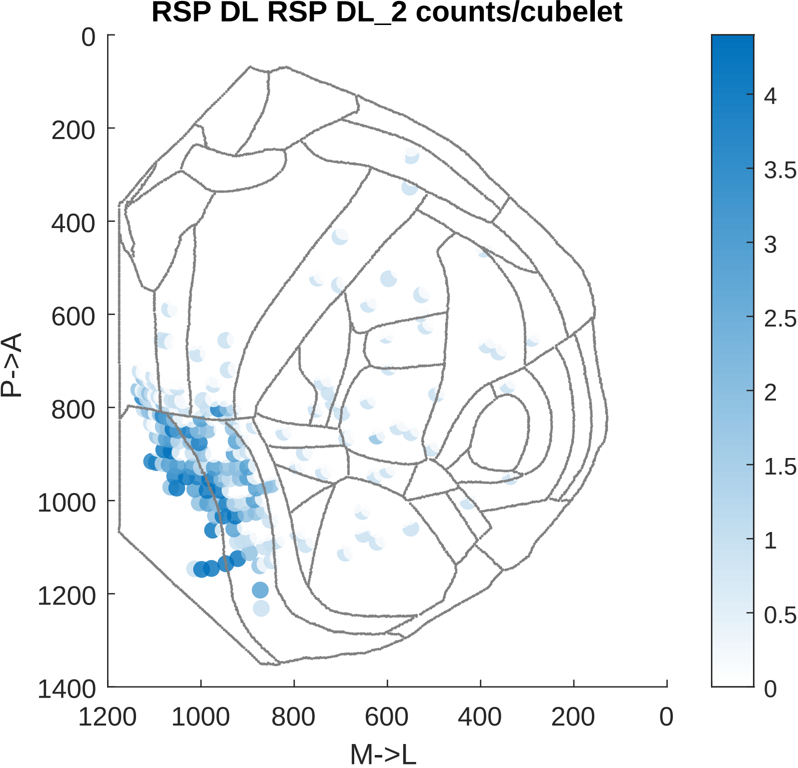
RSP DL RSP DL_2 counts/cubelet
P->A
M->L
0
200
400
600
800
1000
1200
1400
1200
1000
800
600
400
200
0
4
3.5
3
2.5
2
1.5
1
0.5
0

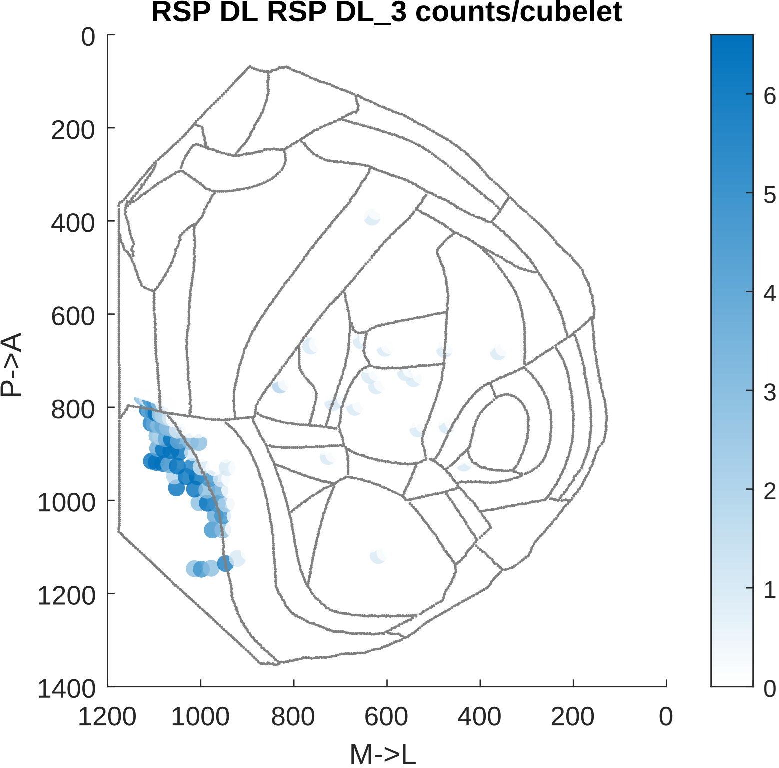
RSP DL RSP DL_3 counts/cubelet
0
200
400
600
800
1000
1200
1400
P->A
1200
1000
800
600
400
200
0
M->L
6
5
4
3
2
1
0

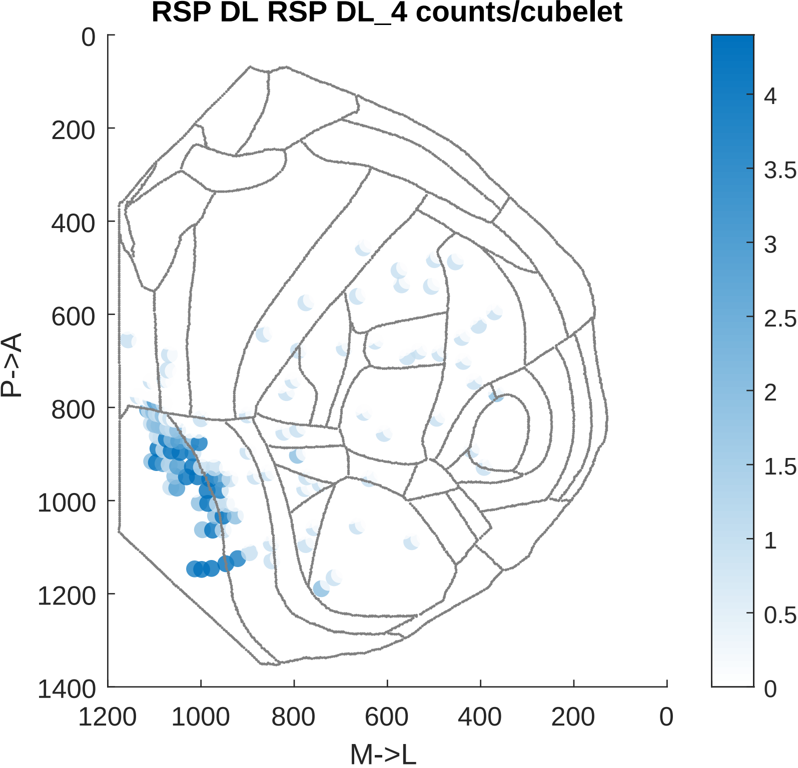

RSP DL RSP DL_4 counts/cubelet
P->A
M->L
0
200
400
600
800
1000
1200
1400
1200
1000
800
600
400
200
0
4
3.5
3
2.5
2
1.5
1
0.5
0

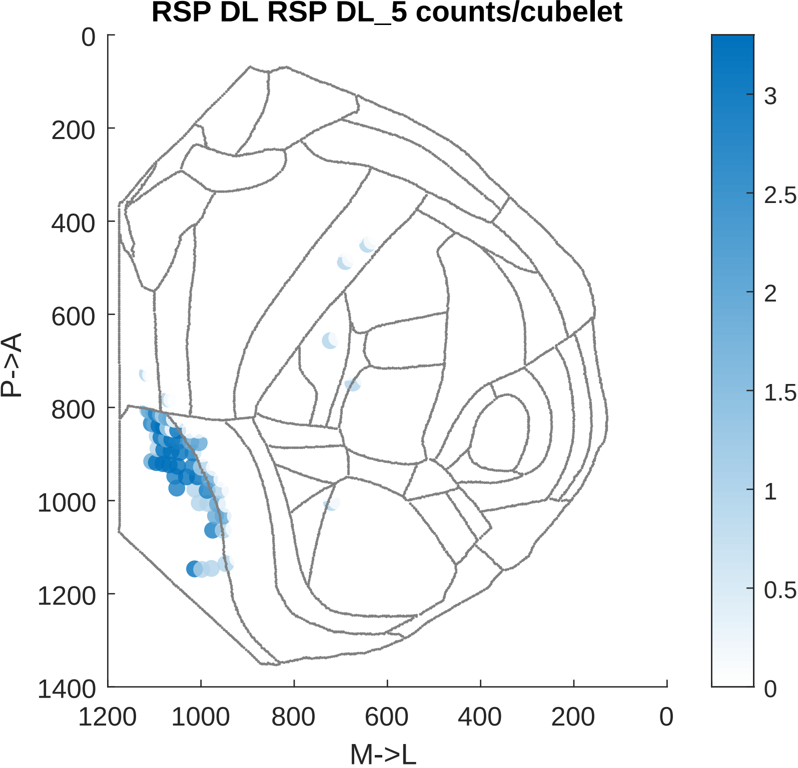

RSP DL RSP DL_5 counts/cubelet
0
200
400
600
800
1000
1200
1400
P->A
1200
1000
800
600
400
200
0
M->L
3
2.5
2
1.5
1
0.5
0

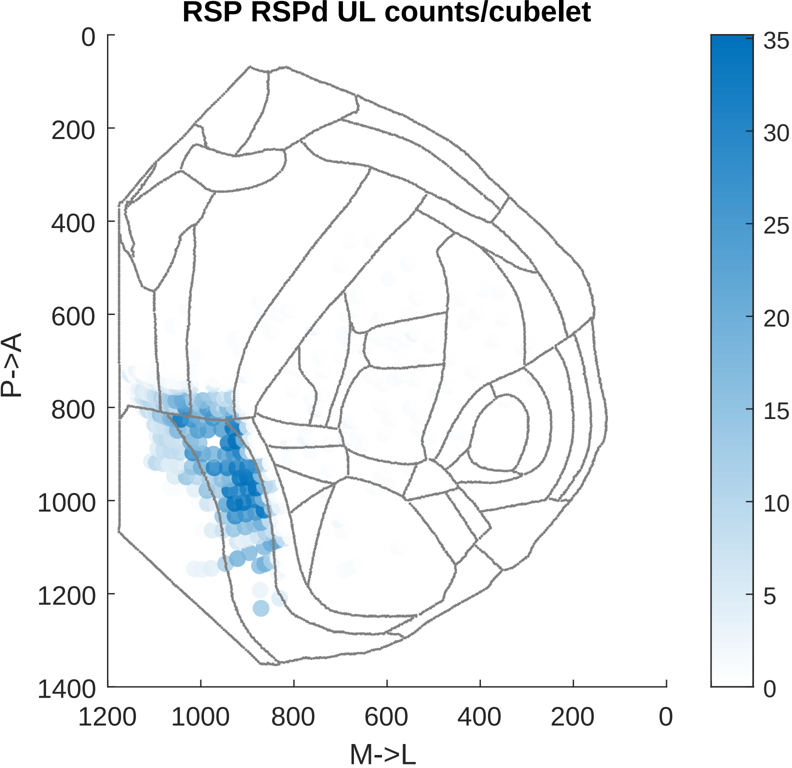
RSP RSPd UL counts/cubelet
P->A
M->L
0
200
400
600
800
1000
1200
1400
1200
1000
800
600
400
200
0
35
30
25
20
15
10
5
0

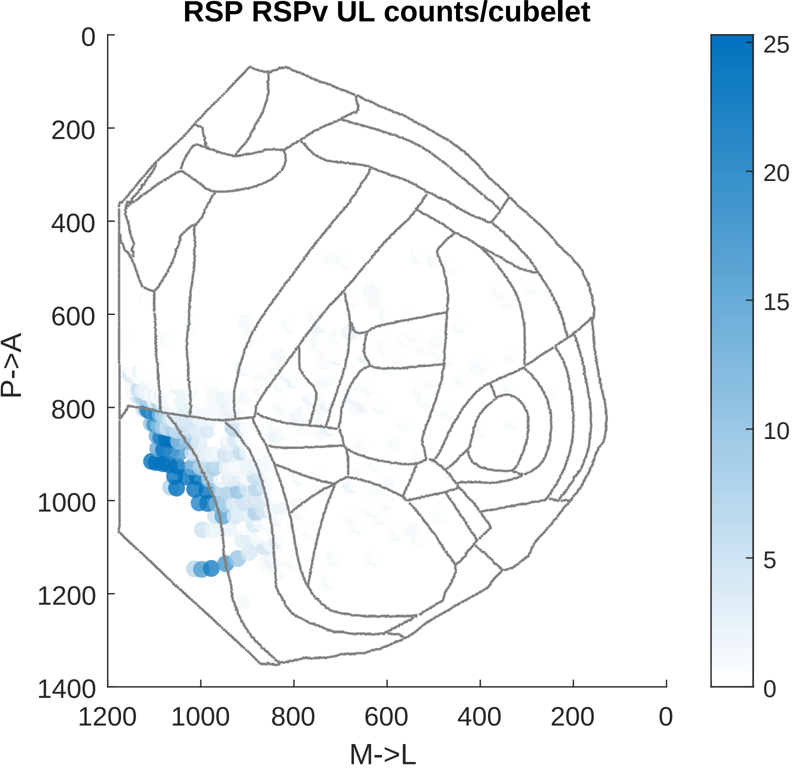

RSP RSPv UL counts/cubelet
0
200
400
600
800
1000
1200
1400
P->A
1200
1000
800
600
400
200
0
M->L
25
20
15
10
5
0
